# Genomic perspectives on the origins of Galápagos iguanas and the evolution of marine specialisation, DNA damage response and pigmentation adaptations

**DOI:** 10.1101/2025.04.13.648612

**Authors:** Julia López-Delgado, Ian M. Carr, Cecilia Paradiso, Paolo Gratton, Giuliano Colosimo, Matthijs P. van den Burg, Adolphe O. Debrot, Christian Sevilla, Mohd Noor Mat-Isa, Richard Bayliss, Matthew Batchelor, Glenn P. Gerber, Mohd Firdaus-Raih, Alan J.S. Beavan, Mary J. O’Connell, Gabriele Gentile, Simon J. Goodman

## Abstract

Island taxa provide opportunities to explore the genomic basis of species diversification and trait evolution arising from adaptation to new environmental conditions and ecological niches. Here we generate the first genomic sequences for the four endemic Galápagos iguana species to reconstruct their evolutionary history and to identify genes which may contribute to their unique adaptations. We show that the marine (*Amblyrhynchus*) and land (*Conolophus*) lineages evolved in situ on now submerged islands, following a single colonisation event 9 - 12.4 million years ago. Genomic selection scans identified genes linked to traits potentially facilitating adaptation to the Galápagos environment, including specialised pigmentation phenotypes (e.g. *ASIP*, *BCO2* and *KITLG*), DNA damage and UV irradiation inflammation responses (e.g. *MAPK14* and *BRAF*), which may contribute to increased resilience to elevated UV exposure at equatorial latitudes. A shift from eumelanin (dark) to pheomelanin (yellow-orange) dominated colouration in *Conolophus* species may be attributable to fixed substitutions in the *MC1R* gene that influence protein structure and interactions, with the depigmented skin phenotype of the pink iguana additionally linked to positive selection in regulators of *MITF* activity, melanosome transport and dermal vasculature unique to that lineage. Further genes with signatures of positive selection in marine iguanas have putative functions including hypoxia response, that may be associated with their transition to underwater foraging (e.g. *HBA*, *HMOX2*, *HIF*), as well as their unique ability to repeatedly shrink and grow in body size through tissue and bone remodelling (e.g. *AREG*, *BMP2*, *CPNE7*). Genome-wide patterns of genetic diversity indicate recent inbreeding coincident with the timing of human settlement. Our study provides insights into the origins and diversification of the iguanas and the molecular basis of adaptation to life in the Galápagos, facilitating future conservation genomic management of threatened iguanid populations.

## Introduction

Remote oceanic islands have long been recognised as natural laboratories for studying evolutionary processes. The Galápagos in particular, with its unique assemblage of species, have helped shape the development of evolutionary theory^1^. The archipelago is situated on the equator in the Pacific Ocean, approximately 930 km west of continental Ecuador. A cycle of island formation driven by volcanism, and subsequent erosion and subsidence occurs as the Nazca tectonic plate moves eastward towards South America over a stationary mantle hotspot^2^. The extant islands emerged from 30,000 to 4.0 million years ago (Mya), but island formation may have been occurring over the hotspot for around 9.1 to 16 million years with older island remnants now submerged^3^. This dynamic geological history has been central to shaping the evolution, biogeography and ecology of the species of the archipelago.

Among the islands’ iconic species, Galápagos iguanas form a monophyletic group of four species in two genera (Fig. 1a, b), the marine iguana (*Amblyrhynchus cristatus*) and land (*Conolophus*) lineage, comprising the yellow land iguana (*C. subcristatus*), the Santa Fe land iguana (*C. pallidus*) and the pink land iguana (*C. marthae*). *A. cristatus* is found on coasts around the archipelago. *C. subcristatus*’s range is spread across five islands in the central archipelago, with *C. pallidus* restricted to Santa Fe, the island from which it takes its common name. *C. marthae* occur only on Wolf Volcano on Isabela and are designated as Critically Endangered in the International Union for Conservation of Nature (IUCN) Red List^4^. Previous work suggests these endemic species diverged within the archipelago from a common Central American ancestor that rafted from the continent^5,6^. The timing of colonisation and lineage divergence within Galápagos remain uncertain, with alternate studies suggesting that differentiation occurred on the extant islands or point towards a much older split on now submerged islands ^5,7^.

**Fig. 1.**
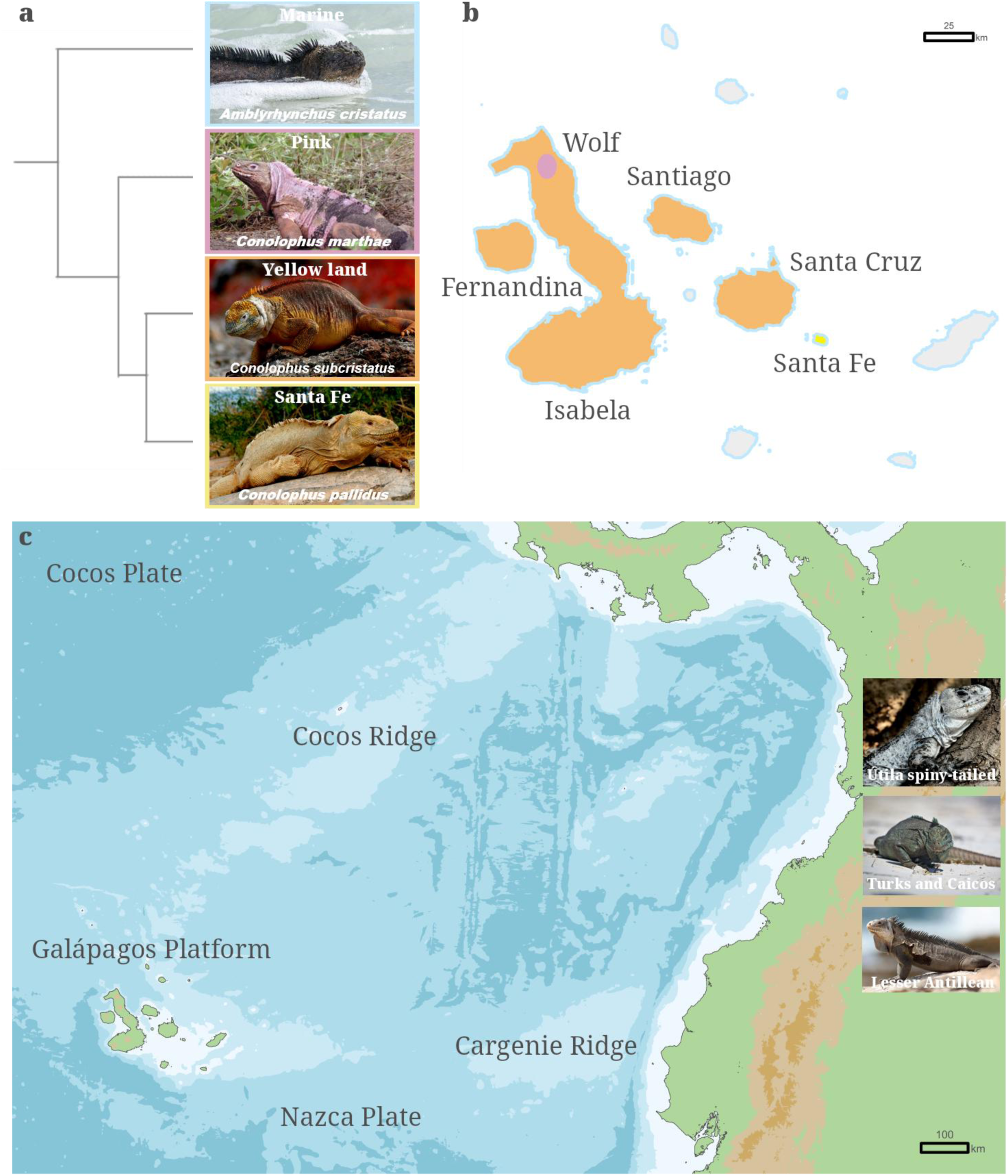
Species relationships and geographic distribution of the Galápagos iguanas. **(a)** Cladogram of the Galápagos iguanas depicting relationships between the four species: marine iguana (*Amblyrhynchus cristatus*; blue), pink iguana (*Conolophus marthae*; pink), Galápagos yellow land iguana (*C. subcristatus;* orange) and Santa Fe land iguana (*C. pallidus*; yellow). **(b)** Distribution of the iguanas on the Galápagos islands, indicated using the respective colours from the cladogram for island shading and outline. **(c)** The geographical setting of the archipelago with shaded isobaths for 500, 1,000, 1,500 and > 1,500 m, ranging from white to dark blue. Position of images do not reflect Caribbean species distributions. Photographs byM. Arechavaleta, W. Tapia, P. Wilton, P. McFarling, J. Müller, T. Sackton.

The diverse habitats, environmental gradients and climatic fluctuations (such as El Niño events) of the Galápagos have driven the evolution of remarkable adaptations in the iguanas. *A. cristatus* transitioned from a terrestrial to a marine niche by evolving a specialised foraging strategyfeeding on benthic algae, requiring extended underwater dives, an exclusive trait amongst lizards. This species also has the unique adaptation of repeatedly shrinking and regrowing in body length, an adaptive strategy to cope with resource scarcity^8^. The pigmentation of Galápagos iguanas ranges from piebaldism in *C. marthae* to black colouration in *A. cristatus*, with implications for photoprotection, thermoregulation, sexual display, camouflage, and habitat choice. Notably, to our knowledge, the Galápagos iguanas have no recorded incidences of cancer, despite prolonged exposure to extremely high levels of ultraviolet radiation^9^.

Here, we generate the first scaffold-level genome sequences for the Galápagos iguanas as well as for three representatives of closely related genera: the Turks and Caicos (*Cyclura carinata*), Lesser Antillean (*Iguana delicatissima*) and Útila spiny-tailed (*Ctenosaura bakeri*) iguanas. We then use the genome assemblies to evaluate competing hypotheses for the origins, and evolutionary and demographic histories of the species. We also compare genome-wide patterns of genetic diversity and mutational burden across Galápagos iguanas and their implications for conservation management. Finally, we analyse signatures of positive selection in gene coding sequences to identify potential molecular underpinnings of the unique adaptive traits of Galápagos iguanas.

## Results

### Origins and evolution of the Galápagos iguanas

To uncover the origins and evolutionary history of the Galápagos iguanas, we assembled and annotated the reference genomes of the four Galápagos species (marine: *Amblyrhynchus cristatus*; yellow: *Conolophus subcristatus*; Santa Fe: *Conolophus pallidus*; and pink: *Conolophus marthae*), as well as the Turks and Caicos (*Cyclura carinata*), Lesser Antillean (*Iguana delicatissima*) and Útila spiny-tailed (*Ctenosaura bakeri*) iguanas. These latter three Caribbean species, as well as being of conservation genomic interest, are hypothesised to be among the closest sister taxa to the Galápagos iguanas^5^ and their inclusion enhances power to: (i) detect signatures of selection in the Galápagos lineage and (ii) test hypotheses about their origins. The reference genomes of the Galápagos iguanas have high gene completeness (BUSCO v.4 mean score of 93.5% with the vertebrata dataset) and moderate contiguity (mean scaffold N50 4.7 Mb; Supplementary Tables S1 and S2). The seven iguana genomes had an average genome size of 2.00 Gb (sd=0.03), repetitive element content of 45.9% (sd=0.8, Supplementary Table S3) and gene count of 22,002 (sd=1,067). We also assembled mitochondrial genomes for the species, which averaged 16,724 bp in length (sd=94) and had the expected 37 genes for vertebrates. In addition, transcriptome assembly and annotation for *A. cristatus* resulted in 27,990 functionally annotated transcripts (Supplementary Data S1).

After stringent filtering to remove genes with hidden paralogy and insufficient phylogenetic signal and retain only loci exhibiting clear one-to-one orthology across all sampled taxa, we performed phylogenomic reconstructions with 229 single gene orthologs (SGOs) across 28 squamates (Supplementary Fig. S1). We also reconstructed phylogenetic relationships using 75% and 95% complete ultraconserved element (UCE) matrices, including unfiltered datasets and subsets of the 500 and 1,000 loci with the strongest phylogenetic signal. The analyses confirmed the basal position of geckos, followed by lacertids and a clade formed by snakes, iguanids and anguimorphs (Fig. 2a). Within the Galápagos clade, the phylogenies supported the initial land-marine split, with *C. marthae* forming the most basal lineage within the terrestrial clade (Fig. 2b). Divergence time estimates from the SGOs and UCE datasets were concordant, placing the origin of squamates in the Early Jurassic period and the divergence between pleurodont and acrodont iguanas in the Early Cretaceous. The split between the Galápagos land and marine iguanas, which illustrates the minimum amount of time that the species have inhabited the archipelago, was estimated at ∼12.4 Mya (95% HPD: 7.5 - 17.8 Mya) using SGOs and ∼9 Mya (95% HPD: 4.8 - 14.7 Mya) using UCEs. The analyses further suggest *C. marthae* diverged ∼5.9 Mya (95% HPD: 2.7 - 9.2 Mya) using SGOs and ∼3 Mya (95% HPD: 1.55 - 5.66) in the UCE analysis, whereas the two other land iguanas split ∼3.6 Mya (95% HPD: 1.3 - 6.5 Mya) and ∼1 Mya (95% HPD: 0.4 - 2.3 Mya), respectively. Calibration densities modelled with uniform and skewed-normal calibration distributions produced very similar divergence times for every node, whereas the Cauchy distribution resulted in much older estimates (Supplementary Tables S4-S10).

**Fig. 2.**
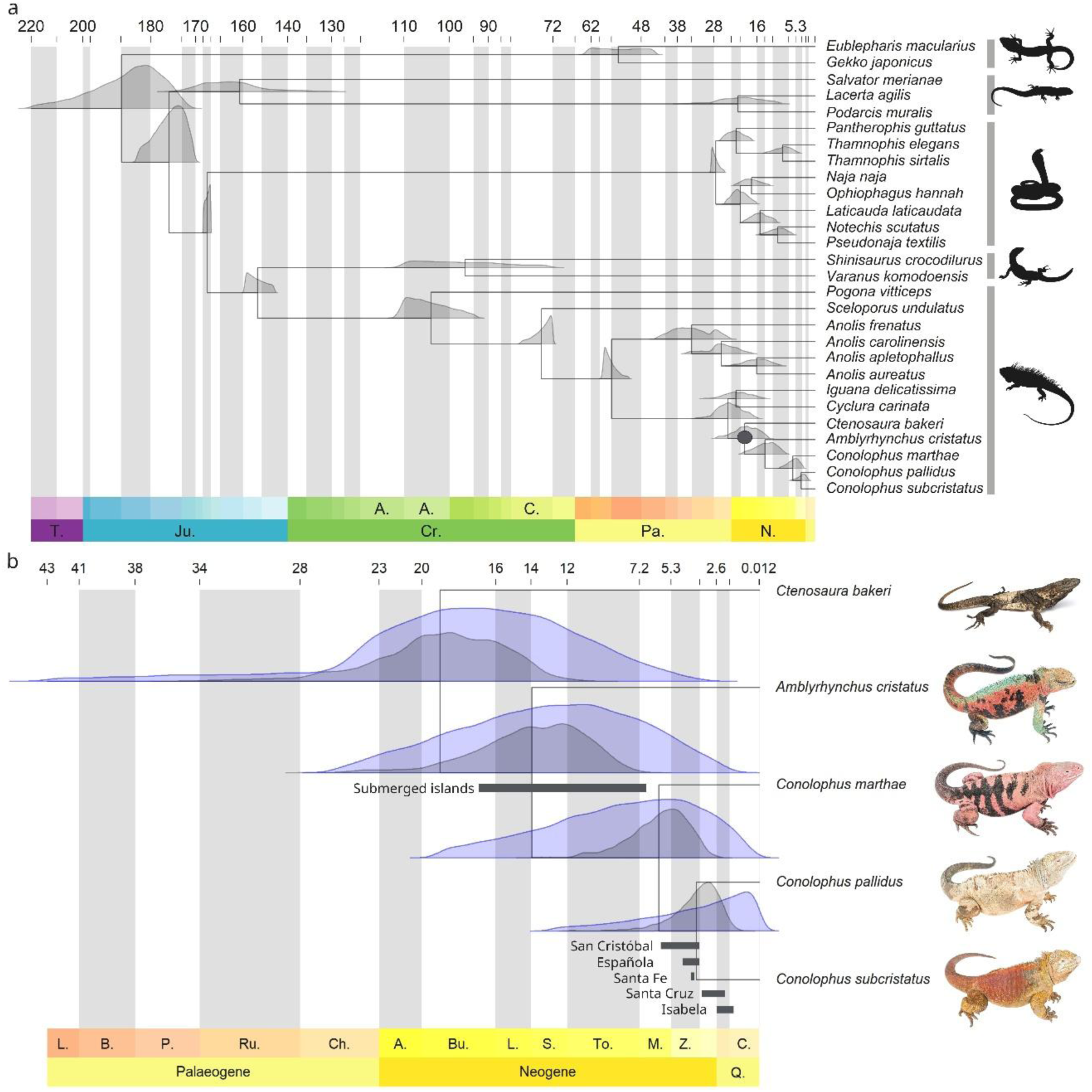
Evolutionary history of squamates and Galápagos iguanas. **(a)** Timescale of squamate evolution estimated with SGOs in MCMCtree under a skewed-normal prior probability distribution displaying the posterior distributions for each node. The geologic periods are labelled on the bottom x-axis, with their corresponding subdivisions represented by alternating white and grey bars, and time in million years are shown on the top x-axis. Calibration points are detailed in the Supplementary Information. Taxonomic subdivisions are represented by vertical bars and correspond to the squamate suborders: Gekkota, Laterata, Serpentes, Anguimorpha and Iguania. The grey circle marks the root node of the clade shown in panel b. Reptile silhouettes are from PhyloPic. (b) Timescale of Galápagos iguana evolution calculated with MCMCtree under a skewed-normal prior probability distribution with posterior distributions for node ages estimated with SGOs (grey) and UCEs (blue). Grey bars denote geological evidence for the emergence of extant and submerged islands in the archipelago. Photographs by and reproduced with permission of J. Vieira and A. Arteaga.

### Genome-wide diversity and inbreeding

We calculated genome-wide heterozygosity and identified runs of homozygosity (ROH) for the iguanid species. *A. cristatus* and *C. subcristatus* had the highest genome-wide heterozygosities (*θ* = 0.00207 and 0.00185, respectively), whereas *C. pallidus*, *C. carinata* and *I. delicatissima* had the lowest values (*θ* = 0.00020, 0.00100 and 0.00021, respectively; Fig. 3a, Supplementary Table S11). *C. marthae* and *I. delicatissima* displayed the highest proportions of the genome in ROH. The other five species had less than 2% of their genomes in ROH, with the lowest values corresponding to *A. cristatus* and *C. subcristatus*. The minimum ROH length was ≥ 1 Mb across all species, which is a common threshold that signals recent inbreeding ^10^. The number of ROH did not increase when specifying increasingly smaller window sizes (Supplementary Table S11). This suggests recent inbreeding and is supported by the low estimate of the number of generations since inbreeding occurred (Fig. 3b). Despite having reduced proportions of ROH, *C. subcristatus* and *C. bakeri* had the longest ROH, and inbreeding occurred as recently as five generations ago. *C. marthae* and *I. delicatissima* display relatively long ROH and inbreeding is predicted to have occurred 7.3 - 16.8 and 8.8 -20.2 generations ago, respectively.

**Fig. 3.**
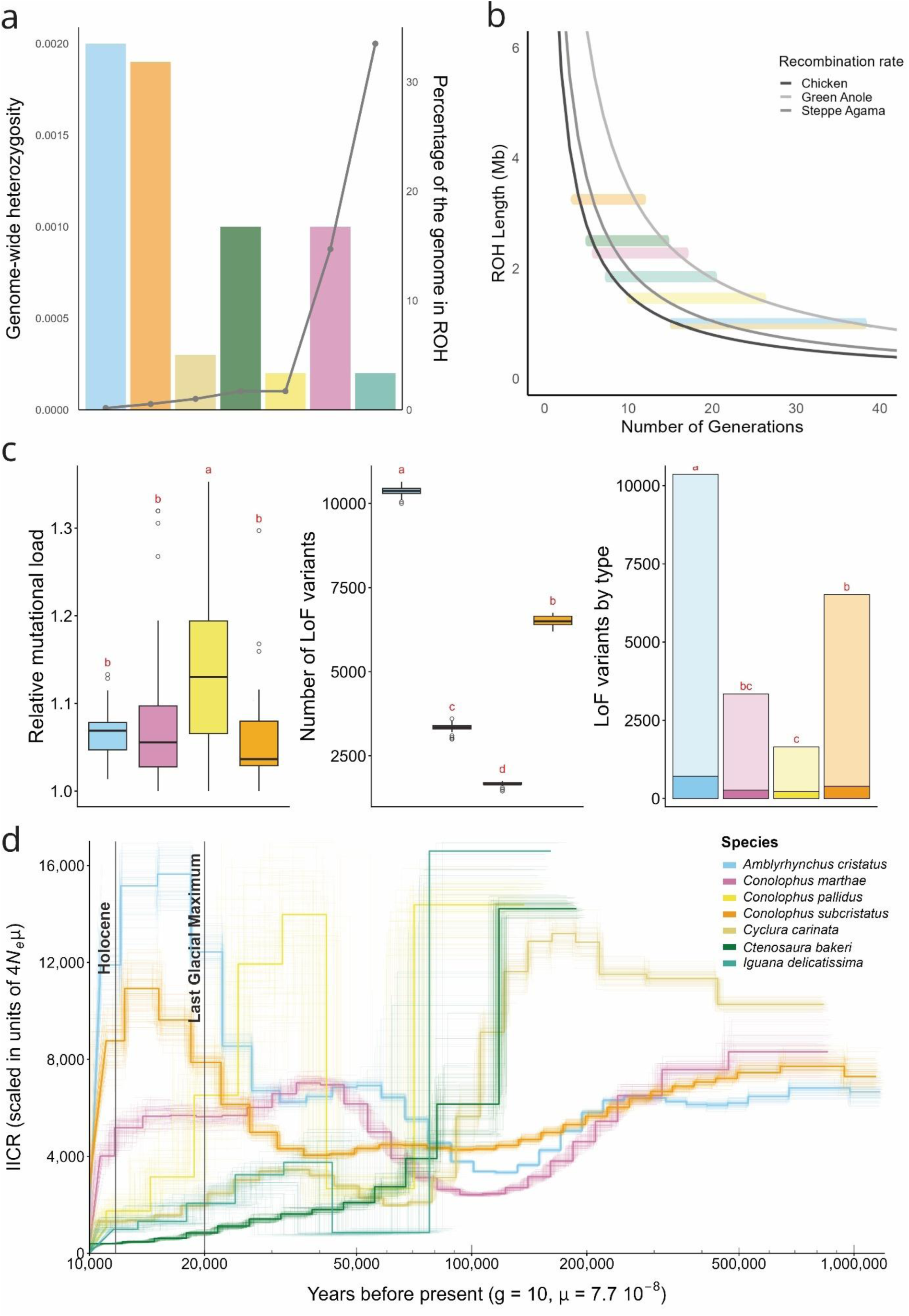
Demographic history of iguanids. **(a)** Patterns of heterozygosity reveal a recent history of inbreeding and ancient demography. Genome-wide heterozygosity (bars), calculated as Watterson’s theta (θ), on the left y-axis and percentage of the genome in Runs of Homozygosity (ROH; points) on the right y-axis and species colour as per key in panel (d). **(b)** Number of generations since inbreeding estimated based on mean ROH length using the recombination rates of the chicken, the steppe agama and the green anole. (**c**) Relative mutational load (sum of derived alleles divided by total sites); total count of Loss-of-Function (LoF) variants; distribution of LoF variants by type, with solid bars representing realized load (homozygous alternative) and semi-transparent shading showing the masked load (heterozygous). All boxplots represent variance across 50 genomic subsamples per individual. Significant differences (Tukey HSD, p < 0.05) between species are indicated by red letters. **(d)** PSMC inference of the effective population size. The x-axis shows time in years before present, calibrated using a generation time (g) of 10 years and a per site per generation mutation rate (μ) of 7.7 x 10^-9^. The y-axis represents the effective population size. The coloured lines show 100 bootstrap replicates for each species. The vertical grey lines represent the Last Glacial Maximum and the onset of the Holocene. The colour-coded species represented in panels a-d correspond to the marine (*Amblyrhynchus cristatus*), yellow land (*Conolophus subcristatus*), Santa Fe (*Conolophus pallidus*), pink (*Conolophus marthae*), Turks and Caicos (*Cyclura carinata*), Lesser Antillean (*Iguana delicatissima*) and Útila spiny-tailed (*Ctenosaura bakeri*) iguanas.

### Mutational burden

Mutational load is a key component of genomic integrity and can provide insights into the distribution of deleterious variation across taxa. Here, we measured the proportion of derived alleles across genomic blocks for the iguanids. The Galápagos taxa revealed contrasting patterns, with *C. pallidus* showing the highest genome-wide relative mutational load and the other species harbouring significantly lower and statistically similar values (Supplementary Table S12 and Supplementary Data S2). By contrast, the number of predicted loss-of-function (LoF) variants was highest in *A. cristatus* (10,363), intermediate in *C. subcristatus* (6,516), and lowest in *C. pallidus* (1,654) and *C. marthae* (3,341). Across all taxa, most LoF variants occurred in the heterozygous state, indicating that most of the deleterious variation is masked rather than realized. However, *C. pallidus* exhibited a comparatively greater fraction of homozygous LoF genotypes (13.72% realized load), whereas *A. cristatus* and *C. subcristatus* carried more LoF variants but lower realized-to-masked ratios (6.87% and 5.97% realized load, respectively).

Functional effect prediction identified high-impact variants in loci associated with pigmentation, cellular stress response, and DNA repair and genome maintenance. In *C. marthae*, frameshift variants were detected in loci homologous to pigmentation regulators, such as *FOXD3*, as well as in candidates involved in UV-related cellular damage and DNA damage response, including *ATMIN, FANCM*, *ASF1*, and *FOXO3* ^11,12^. In *A. cristatus*, putative exon-loss variants were identified in loci homologous to pigmentation-related genes (*TYRP1*, *DCT*), growth factor and metabolic regulators (*FGF9*, *FGF16*, IGFBP), and genome maintenance (*TRIM29*, *RUVBL1*, *MCM3*). Similarly, putative exon-loss variants in *C. subcristatus* affected pigment-cell regulatory (*FOXD3*) and DNA repair and oxidative stress response genes (*PARP1*, *GPX7*), and in *C. pallidus*, chromatin-associated DNA damage response factors (*ASF1B*, *KAT2B*) ^13, 14^.

### Intraspecific demographic histories

The demographic history of the seven iguanids was investigated by Pairwise Sequentially Markovian Coalescence (PSMC) analysis of their genomes (Fig. 3d). The Galápagos species show broadly similar effective population size (*N_e_*) trajectories during the mid-late Pleistocene, with relatively stable *N_e_* followed by a gradual decline around one hundred thousand years ago (100 Kya). The *N_e_* of *A. cristatus* and *C. subcristatus* increased in the period between 40 to 15 Kya and then dropped rapidly. The analysis was unable to reconstruct the *N_e_* of *C. pallidus* and *I. delicatissima* past 100 Kya, since local heterozygosity densities were probably insufficient for PSMC to infer ancient recombination events and coalescent times.

### Signatures of positive selection

To test for variation in selective pressure in the Galápagos iguanas, we considered terminal tip (individual species) and ancestral branch levels in a set of 6,682 SGOs. These SGOs were filtered to remove paralogs, with those containing all Galápagos species, at least two non-Galápagos iguanids, and two non-iguanid species retained for further analysis (Supplementary Table S13 and Supplementary Data S3). The *A. cristatus* showed the highest number of candidate genes with signatures of positive selection (421 lineage-exclusive genes; Table S14). In addition, the stem *Conolophus* lineage had more genes under positive selection than the Galápagos branch (248 and 108 lineage-exclusive genes, respectively). Gene Ontology (GO) slims were enriched with key terms including DNA repair and replication, inflammatory response and anatomical structure development in all lineages (Supplementary Fig. S2). Analysis at the gene family level revealed the Galápagos iguanas have a high number of gene gains and losses (Supplementary Fig. S3 and Supplementary Data S4), with the *A. cristatus* displaying the highest number of both gene family expansions and contractions (1,040 gene family expansions and 1,701 contractions; Supplementary Table S15).

*A. cristatus* is anatomically and physiologically specialised to thrive in the ocean ^8^. Genes with signatures of positive selection in *A. cristatus* have functions related to oxygen management, cardiovascular control, cell protection and anatomical modifications that appear to contribute towards hypoxia tolerance in this species (Fig. 4). Overrepresented GO terms across the positively selected genes are associated with swimming, adaptive thermogenesis, respiratory burst and haemoglobin biosynthetic process. Both paralogs encoding the alpha subunit of the Hypoxia-Inducible Factor (*HIF1A* and *HIF2A/EPAS1*) have evidence of positive selection. Genes linked to haemoglobin affinity and oxygen binding and transport are positively selected in this species, including *HBA*, *HMOX2* and *BPGM*. We detected positive selection in genes related to osmoregulation and antioxidation (e.g. *GPX7*, *IGF1* and *STC1*), including aquaporins and solute-carriers (Fig. 4). *A. cristatus* also shows signatures of positive selection in genes related to cardiovascular control, including genes involved in vasoregulation, coagulation and the renin-angiotensin system such as *AGT*, *APLNR*, *ENPEP* and *TFPI2*.

**Fig. 4.**
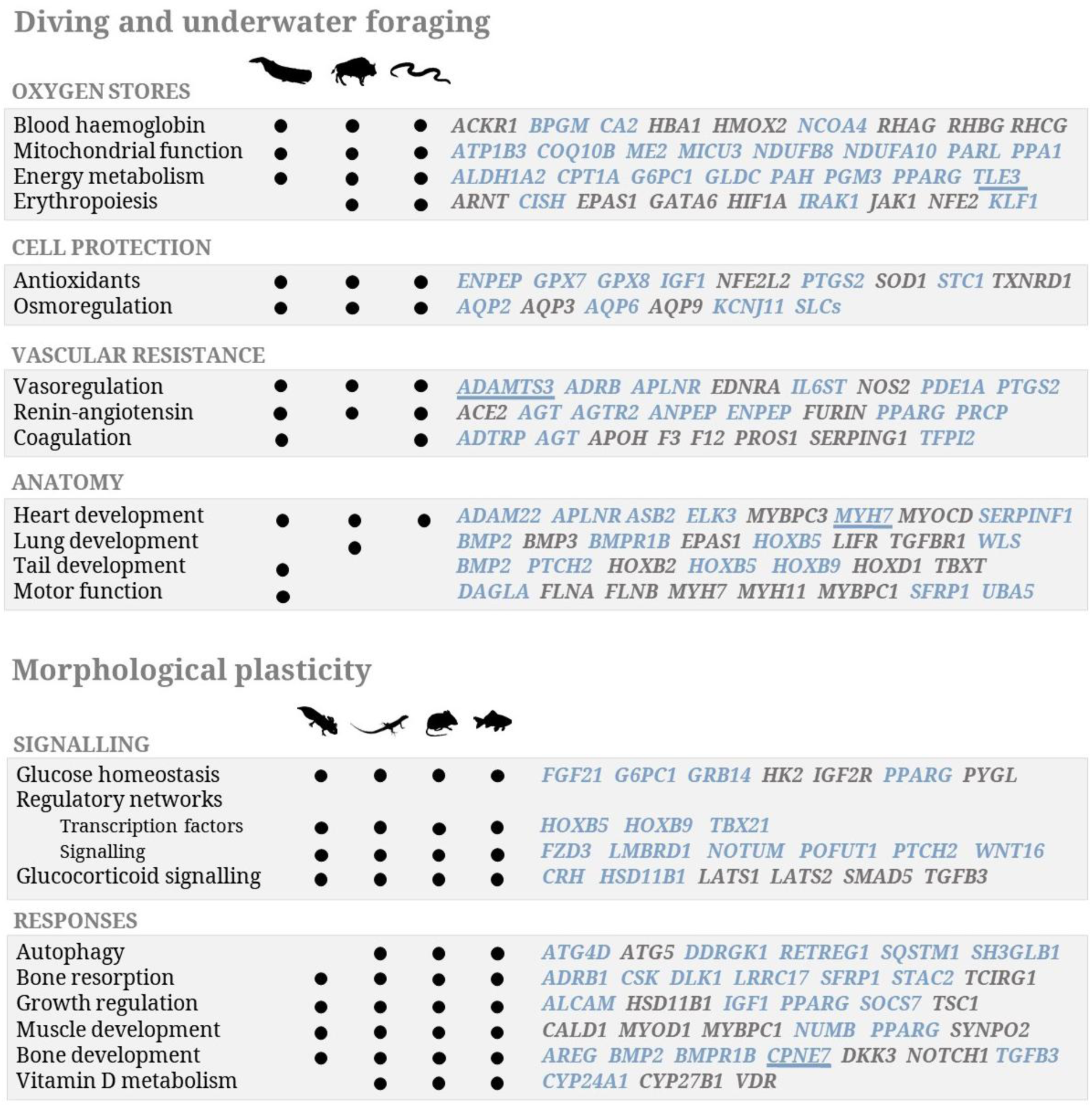
Genomic signatures of marine iguana adaptations. Top panel shows key candidate genes linked to the transition to diving and underwater foraging under positive selection in the marine iguana with known adaptive functions in marine mammals (whale icon), high-altitude species (yak silhouette) and aquatic reptiles (snake figure). Bottom panel shows the genomic signatures of morphological plasticity in the marine iguana. Key candidate genes under positive selection in the marine iguana linked to morphological plasticity with known adaptive functions in amphibians (axolotl icon), reptiles (lizard icon silhouette), mammals (mouse icon figure) and fish. Genes shown in blue are exclusively under positive selection in the marine iguana; genes in grey are also under positive selection in other lineages; and underlined genes have undergone gene family expansion or contraction.

*A. cristatus* exhibits high plasticity in body size driven by food availability^8^. Genes with signatures of positive selection in *A. cristatus* have been associated with morphological plasticity in other vertebrates, contributing to bone homeostasis, muscle development and growth regulation (Fig. 4). These phenotypic responses are mediated by molecular signalling pathways regulated by nutrient sensing and stress hormone release^15^. *A. cristatus* has genes with signatures of positive selection involved in the WNT, Hedgehog, TGF-β and Notch signalling pathways and regulatory networks, as well as in bone resorption and formation, glucose homeostasis and glucocorticoid metabolism. For instance, *AREG* promotes bone homeostasis^16^ and *CPNE7* induces osteogenic differentiation to promote bone tissue regeneration^17^. Notably, *PRKAA2*, the gene encoding the master regulator of energy homeostasis AMPK, has evidence of positive selection in *A. cristatus*.

The Galápagos iguanas also show signatures of positive selection in genes involved in DNA damage sensing, repair pathways and tumour suppression (Fig. 5a). The DNA damage sensors *ATR* and DNA-PK (encoded by *PRKDC*), which transduce DNA damage signals from UV radiation to checkpoint control proteins such as *RAD17*^18^, are under positive selection in most Galápagos species (Fig. 5a). We identified 17 Mitogen-Activated Protein Kinases (MAPKs) and MAPK kinases under positive selection in Galápagos iguanas, including the oncogenes *ARAF* and *BRAF* as well as *MAPK14*, which encodes for the critical effector in cellular stress responses p38-α^19^. p38 can be activated by the DNA damage sensors ATM and ATR or directly by UV radiation, regulating downstream targets including several kinases, transcription factors and cytosolic proteins (Fig. 5b,c). We found that the MAPK14 protein in the Galápagos iguanas has an extended N-terminal region and a unique pattern of sequence variation across its MAPK insert, which is a critical structural motif in the interaction with regulatory partners. Protein structural modelling of Galápagos iguana variants implies that the amino acid substitutions are likely to drive differences in the interaction of *MAPK14* with its upstream kinases *MAP2K3* and *MAP2K6* (Fig. 5b). In response to DNA damage, p38 phosphorylates and activates the tumour suppressor protein p53, a key regulator of genomic integrity^20^. Several genes from the p53 family were also under positive selection in the Galápagos iguanas. Additional selected genes are involved in major DNA repair systems, oncogenesis and tumour suppression (Fig. 5a).

**Fig. 5.**
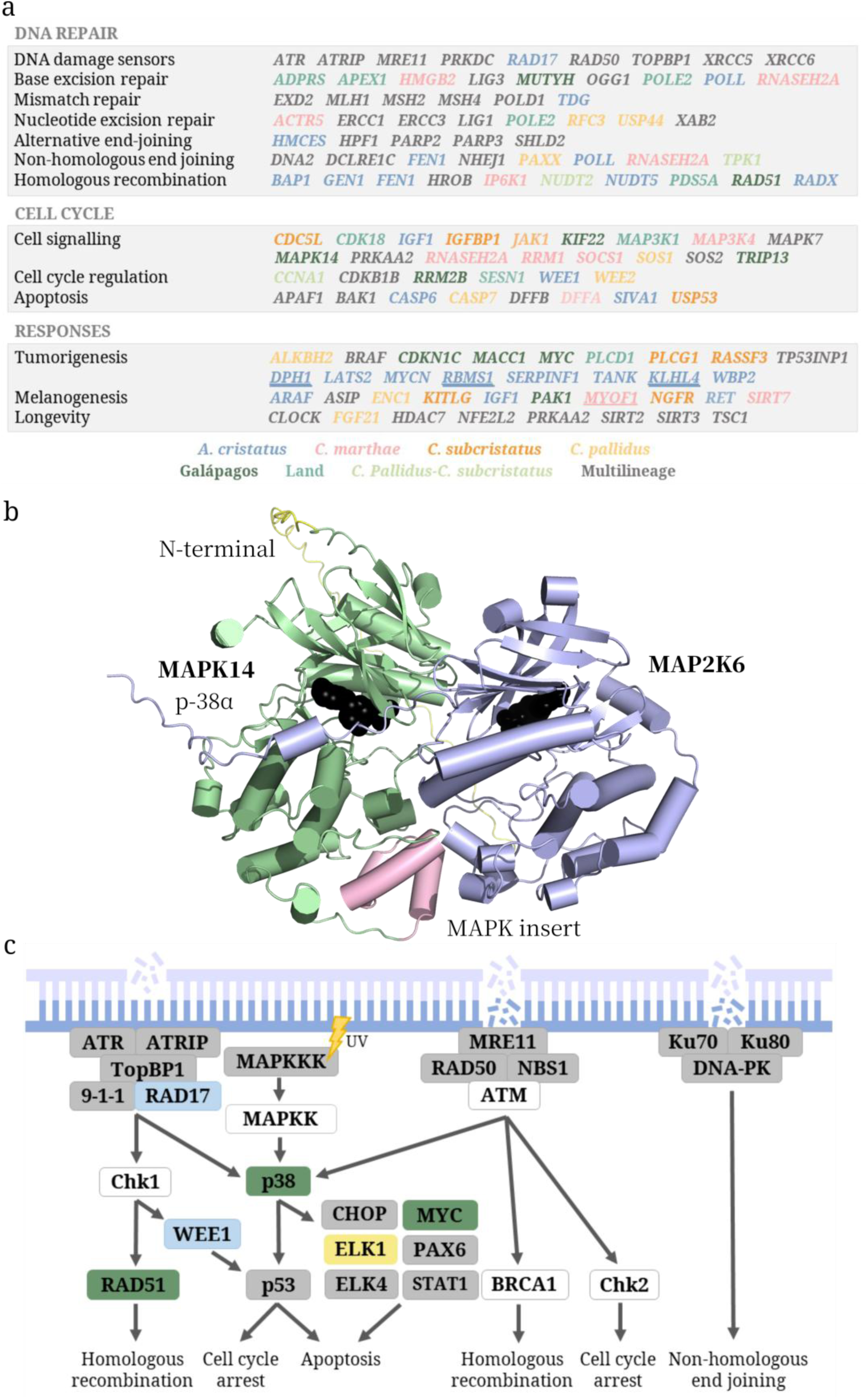
Signatures of positive selection in the DNA damage response in the Galápagos iguanas. **(a)** Key candidate genes linked to the DNA damage response involved in damage sensing, DNA repair, cellular and general responses under positive selection in each Galápagos species or branch tested. DNA repair strategies include base excision repair, nucleotide excision repair, mismatch repair, non-homologous-end-joining and homologous recombination. Underlined genes have undergone gene family expansion or contraction. **(b)** 3D protein model of the MAPK14 (p38-α, green) and MAP2K6 (purple) complex for the pink iguana, showing the N-terminal region (yellow) and MAPK insert (pink) of MAPK14. **(c)** The networks of pathways activated during the DNA damage response. Positively selected genes are coloured by lineage using the respective colours from (a), with grey indicating a gene was under positive selection in more than one lineage. Single strand breaks result in the recruitment of ATR/ATRIP, whereas double strand breaks are detected by the Ku70/Ku80 or the MRN complexes, all of which recruit and activate signal transducers.

Signatures of positive selection were also found in key genes involved in pigment cell differentiation and pigment synthesis (Fig. 6). In *A. cristatus*, many genes related to melanogenesis and melanophore differentiation showed evidence of positive selection (Supplementary Data S4). These include the key pigmentation gene *ASIP* and transcription factors that modulate the master regulator of melanocyte development *MITF* such as *TFEC*, which also plays a crucial role in chromatophore development in reptiles^21^. Essential genes for carotene uptake and metabolism, such as beta-carotene oxygenase genes, also showed signatures of positive selection in the marine species^22^. In land iguanas, genes involved in xanthophore differentiation, pteridine synthesis, purine binding and carotenoid metabolism were under positive selection. Comparative sequence analysis of the *MC1R* coding sequence revealed five amino acid substitutions unique to all *Conolophus* species (Fig.7) at positions 67 (located in intracellular loop 1, R67Q), 181 (found on the extracellular loop 2, V181I), 186 (on the extracellular loop 2, A186P), 234 (in intracellular loop 3, A234P) and 312 (on transmembrane helix 7, V312L). Structural mapping revealed that the R67Q substitution (arginine to glutamine) alters the stimulatory G protein beta subunit contact interface, while the V181I substitution (valine to isoleucine) perturbs the ligand-binding loop adjacent to the direct α-MSH pharmacophore contact residue ILE180 (3.48 Å from ARG8). Similarly, the A186P substitution (alanine to proline) introduces a rigid, helix-distorting residue close to the α-MSH binding site, potentially altering the local backbone geometry of extracellular loop 2 and influencing ligand accommodation. The A234P substitution (alanine to proline) likewise replaces a flexible residue with proline within intracellular loop 3, where Pro230 is positioned close to the Gα binding interface, suggesting a possible role in modulating the conformational dynamics required for G protein coupling. In *C. marthae*, GO terms associated with developmental pigmentation were strongly overrepresented. Evidence of positive selection was detected in transcription factors and signalling genes linked to *MITF* regulation in this species, including *TCF7* and *JAG1*, which converge on *MITF* from independent signalling axes, as well as melanosome transport and vascular tone genes (Fig. 7). Notably, two genes from the myosin superfamily were under positive selection in *C. marthae*, *MYH11* and *MYO1F*, with the latter also having undergone gene family expansion.

**Fig. 6.**
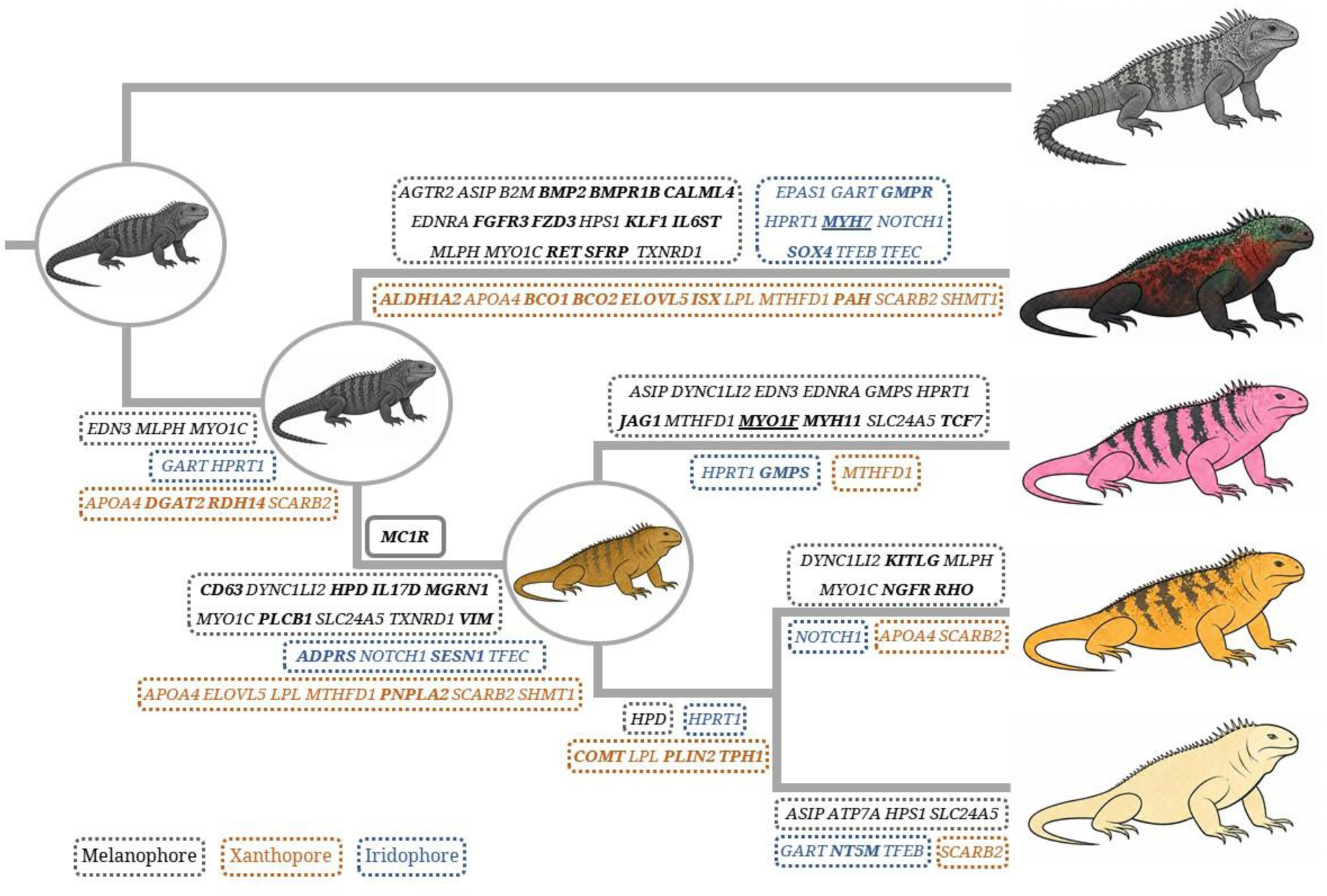
Pigmentation synthesis and transport genes show lineage-specific signatures of positive selection in the Galápagos iguanas. Key candidate genes linked to pigment synthesis and differentiation of pigment cells under positive selection in each Galápagos species or branch tested; with genes related to melanophore differentiation and melanin synthesis and transport in black, genes involved in pteridine synthesis and carotenoid uptake and deposition in orange, and genes taking part in iridophore differentiation and purine metabolism in blue. [Note: purine metabolism and binding are also involved in pteridine synthesis.] Genes in bold are under lineage-exclusive positive selection and underlined genes have also undergone gene family expansion or contraction.

**Fig. 7.**
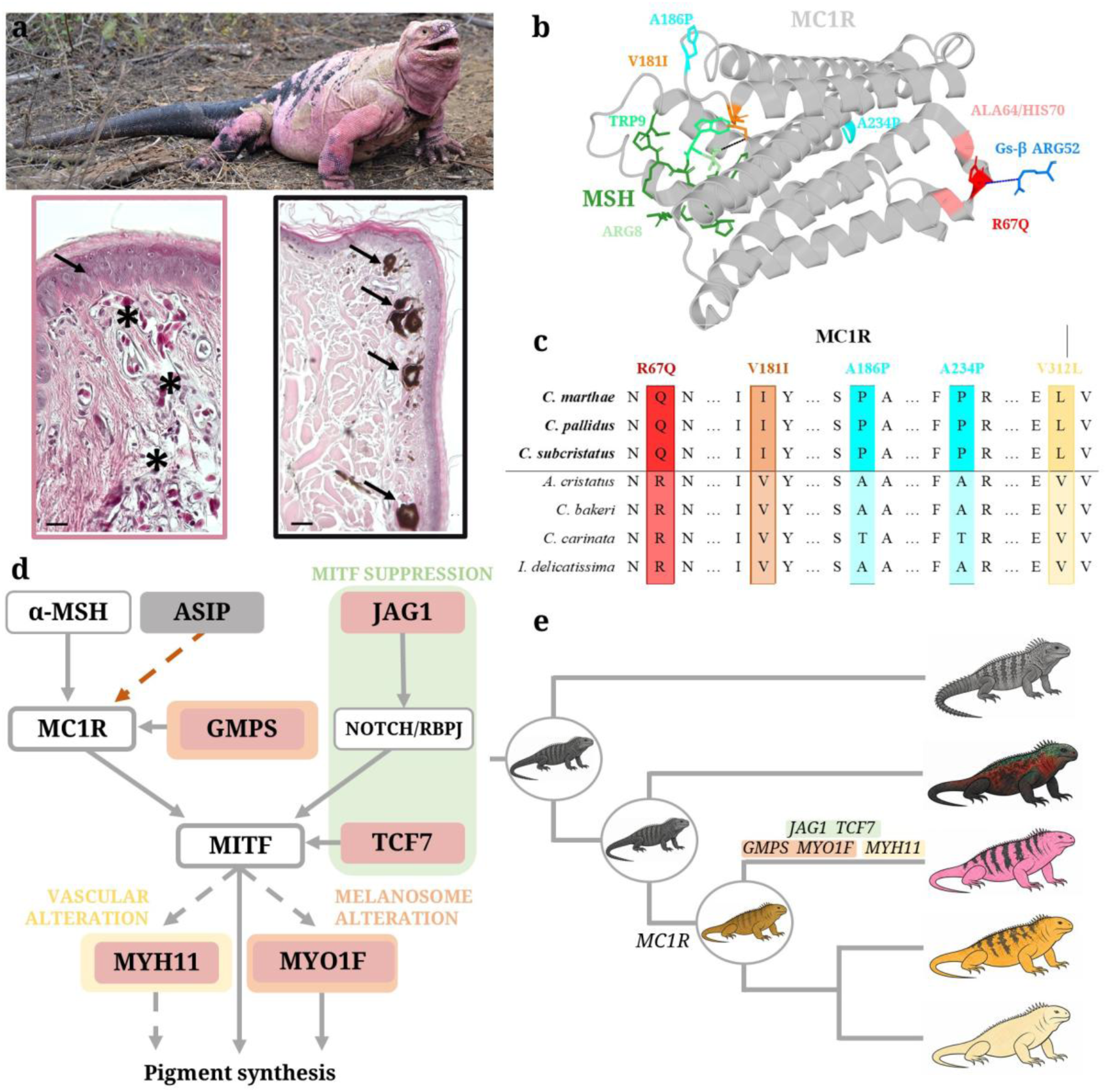
Molecular basis of depigmentation in *Conolophus marthae*. **(a)** Photograph of the pink iguana by and reproduced with permission of L. Garizio. Skin histology images of *Conolophus marthae* are from Scimeca et al. ^77^, reused with permission from the authors. Non-black-colored (pink) skin of *C. marthae* on the left characterized by a thick superficial loose dermis (arrow) and massive blood vessels (asterisks). Black-colored-skin of *C. marthae* on the right displays numerous melanocyte cells (arrows). Scale bar = 100 μm. **(b)** Structural mapping of *Conolophus*-unique *MC1R* substitutions onto its cryo-EM structure (PDB: 9K3P). The receptor is shown in grey. α-MSH is shown in green, with the HFRW pharmacophore residues ARG8 (pale green) and TRP9 (lime green) highlighted. Mutation sites are shown as: ILE181 (orange, V181I) and the adjacent contact residue ILE180 (dark orange), which contacts MSH ARG8 at 3.48 Å (black dashed line); GLN67 (red, R67Q) with intrachain H-bond partners ALA64 and HIS70 (salmon). Blue sticks indicate Gs-β subunit ARG52, which contacts R67Q at 3.83 Å (blue dashed line). Human residue numbering is used throughout. **(c)** Left side displays the multiple sequence alignment of *MC1R* showing the three amino acid positions uniquely derived in all *Conolophus*. Alignment positions 67 (R67Q), 181 (V181I), 186 (A186P), 234 (A234P) and 312 (V312L) are highlighted; the substituted residue in *Conolophus* (Q, I and L respectively) differs from the outgroup state (R, V and V) at all three sites. Mutation site colours correspond to those in panel (a). **(d)** Pigmentation regulatory network underlying the depigmentation phenotype of *Conolophus marthae*. Three convergent mechanisms are proposed to drive the pink depigmentation phenotype: (1) ***MITF* suppression** (green shading): *JAG1* (via NOTCH/RBPJ) and *TCF7* independently converge to suppress *MITF* transcriptional activity, reducing melanocyte differentiation; (2) **melanosome alteration** (orange shading): *MYO1F* modifies melanosome transport downstream of *MITF*, while *GMPS* modulates GTP pool availability upstream of MC1R, flanking the core signalling node; (3) **vascular alteration** (yellow shading): *MYH11* may contribute to the pink coloration through vascular smooth muscle remodelling, independently of melanocyte biology. Nodes represent genes or signalling complexes organised by functional tier. Pink fill indicates genes under positive selection exclusively in *C. marthae*; grey fill in *ASIP* indicates positive selection across *A. cristatus, C. marthae* and *C. subcristatus*. Solid arrows indicate direct activation, dashed arrows indicate indirect or regulatory interactions, and the orange dashed arrow indicates inhibition. (e) Cladogram showing the molecular basis of depigmentation in the pink iguana, facilitated by loss-of-function mutations in *MC1R* in the *Conolophus* lineage.

## Discussion

Here, we generated the first genome assemblies for the Galápagos, Turks and Caicos, Lesser Antillean and Útila spiny-tailed iguanas. These resources represent an important advancement in sequencing squamate genomes, which remain under-represented in genomic repositories, and support our focal phylogenomic and molecular evolution analyses for Galápagos taxa. Our reconstructions of squamate relationships based on protein-coding genes and UCEs confirmed the basal position of geckos, followed by lacertids and the Toxicofera clade, in line with recent phylogenetic analyses^23,24^. Divergence time estimation suggests that squamates originated in the Jurassic, concordant with previous studies^25,26^. Whilst our estimates on the origins of squamate groups largely agree with previous dating from molecular phylogenies, at the rank of suborder to genus we propose an older split for snakes and anguimorphs^25^. This finding likely results from our calibration having included the oldest known fossil snake, which was dated 70 My earlier than previous estimates for the origin of snakes^27^, which highlights the importance of incorporating high-quality fossil calibrations in divergence time estimation^28^. We applied stringent filtering and tested multiple marker and calibration point subsets to ensure that phylogenies and divergence-time estimates were based on true SGOs and informative UCEs, an approach shown to produce robust phylogenetic inferences across diverse taxa^29,30^. Although this filtering necessarily reduced the number of loci available for reconstruction, our results are congruent with previous phylogenies of the group^6,7,29,30^. As more genomes become available, the resolution of future dedicated phylogenomic studies could be enhanced by incorporating additional taxa and genomic regions, such as rapidly evolving long exons, that have shown high phylogenetic informativeness in squamates^31,32^.

The monophyly of the Galápagos lineage and divergence time estimates imply that the iguanas evolved from a single colonisation event soon after land became available in the Galápagos platform. We estimate the initial divergence among Galápagos iguanas occurred 9 - 12.4 Mya according to UCE and SGO estimates, respectively. This is consistent with previous findings from RADseq data, mitochondrial rRNA sequences, genome-wide exons and ultraconserved elements, immunological comparisons of serum albumins and electrophoresis of plasma proteins support a similar timescale^6,7,33,34^. The divergence time estimates for the Galápagos iguanas do not overlap with extant island ages^2^ (< 4 Mya), implying that the species originated and diversified into the marine and land lineages on now submerged islands. This scenario is supported by palaeogeographical data on the ontogeny of proto-Galápagos islands^3^, with direct evidence for subaerial erosion of sunken seamounts to the southeast of the extant islands 9.1 - 16 Mya^35,3^. In addition, plate motion reconstructions suggest that significant subaerial landscapes existed for at least the past 20 My along the Carnegie-Cocos hotspot tracks leading from America to the archipelago^3^ (Fig. 1c). The Galápagos also host other endemic taxa with reported divergence times older than the oceanic islands, including the *Galapaganus* weevils^36^, *Phyllodactylus* geckos^37^ and *Pseudalsophis* racer snakes^38^.

Following the initial colonisation, the iguanas likely spread onto the new islands in a northwest direction parallel to island formation, as reflected in the population structure patterns of marine and land iguanas^39,40^. Notably, the pink iguana (*C. marthae*), the most ancient species in the land lineage is restricted to the youngest island^2^ (0.5 - 0.8 Kya). The estimated origin of the *C. marthae* at 3 - 6.1 Mya based on our genomic data, suggests the pink and yellow land iguanas (*C. subcristatus*) that inhabit Isabela must have colonised the island separately. Similarly, data from MacLeod et al.^41^ (four single-copy protein-coding nuclear genes) and Paradiso et al.^7^ (30,672 SNPs from RADseq data) are also consistent with allopatric speciation followed by separate colonisation events of Isabela.

Our estimates for the marine - land split are older than those estimated by MacLeod et al.^41^ (2.8 - 6.7 Mya), Malone et al.^5^ (2.6 - 8.6 Mya) and Scarpetta et al.^6^ (4.8 - 8.1 Mya). Taxon composition, number and type of loci, calibration point choices and strategies, can all influence divergence times recovered in phylogenomic analyses^6^. The older studies of MacLeod et al.^41^ and Malone et al.^5^ were constrained on the number of loci available to them, and calibrated their molecular clock analyses by placing most of the prior probability mass near the estimated age of fossils descending from each node and with fossil age as a hard boundary for the minimum node age, which could have favoured an underestimation of divergence times^42^. Scarpetta et al.^6^ inferred divergence times from a large multilocus dataset (including rapidly evolving long exon capture, anchored-hybrid enrichment, UCEs, and Squamate Tree of Life loci), but their taxon sample focused on Iguania, using three fossil calibration points, with a shallower total time depth compared to our phylogeny. These factors along with other methodological differences may contribute to the variation in estimates among studies^42^.

Interestingly, the SGO HPD estimate for the *C. subcristatus* - *C. pallidus* divergence (3.6 Mya), is a close match for the estimated emergence of Santa Fe (2.9 Mya)^2^. For splits within the land lineage, we recover older times than Paradiso et al.^7^. While Paradiso et al.^7^ applied an average substitution rate for lizards to scale the species tree, we used a fossil-calibrated relaxed molecular clock, which can yield older divergence estimates by incorporating lineage-specific rate variation and paleontological constraints^43^. The youngest calibration point available for the current study is 23 - 28.4 Mya within the snake clade, and an absence of more recent points may lead to an overestimation of node ages close to the tree tips^44^. Additionally, differences in assumptions around *N_e_* and gene flow between phylogenomic and coalescent-based approaches^7^ may also affect estimates of divergence times. Further whole genome studies at the population level, and a wider exploration of the influence of calibration point availability will help to clarify divergence times within the land lineage^7^.

We applied best practice for fossil calibration^45^, used stringent filtering for the inclusion of SGOs, and explored a range of model and parameter settings. To evaluate the effect of marker choice, we also estimated divergence times with UCEs. While UCEs yielded younger posterior ages compared to SGOs, the HPD intervals overlap, indicating that overall, our results are robust to marker and methodological choices and are a reliable reflection of the phylogenomic signal of the data.

We find low genome-wide heterozygosity in *C*. *marthae* (*θ* = 0.00105) and *C. pallidus* (*θ* = 0.00020), comparable with the values of other endangered reptiles such as Komodo dragons (*θ* = 0.00013) and the Indian gharial (*θ* = 0.00021)^46,47^. *A. cristatus* and *C. subcristatus* have moderate to high levels of heterozygosity (*θ* = 0.00207 and 0.00185, respectively), in the range typical for non-endangered vertebrates. In addition, *A. cristatus* and *C. subcristatus* carried a higher number of predicted loss-of-function variants but lower realized-to-masked ratios, consistent with a larger pool of deleterious alleles segregating in heterozygous form ^48^. In line with its reduced genomic diversity, *C. pallidus* showed the highest relative mutational load and a comparatively greater proportion of homozygous loss-of-function genotypes, indicating a higher realized mutational load^48^. Both *C. marthae* and *C. pallidus* exist as single populations restricted to very small geographic areas, with the pink iguana consisting of only 150 - 270 adults^49^, putting them at risk from stochastic catastrophes and further erosion of genetic diversity and adaptive potential. Prolonged population bottlenecks in these species may have driven the homozygosity or fixation of deleterious alleles, thereby increasing realized load and its potential fitness consequences. These processes may increase genomic vulnerability and inbreeding risks^47^, with potential fitness reductions, particularly in *C. pallidus*. Paradiso et al.^50^ found that the purging coefficient was significantly lower in *C. pallidus* than other Galápagos iguanas, consistent with weaker purifying selection that allowed deleterious mutations to accumulate in this species.

In the Galápagos iguanas, the timing of ROH coalescence coincides with the arrival of whalers and pirates in the 18^th^ century and colonisation in the 19^th^ century. Humans introduced invasive species including cattle, cats, dogs and rats. These have had a strong impact on iguana population sizes via resource competition, habitat alteration and predation, since iguanas have negligible anti-predation behaviour due to having evolved in the absence of terrestrial predators^51^. Despite having high heterozygosity, *C. subcristatus* showed the highest average ROH length (3.25 Mb), which is potentially indicative of very recent inbreeding. This pattern has also been reported in the blue-tailed skink^52^ and the California condor^53^. Our *C. subcristatus* specimen was sampled from Wolf Volcano, where predation pressure from non-native animals is particularly high^51^ and could have led to a reduction in population size and gene flow that resulted in inbreeding within the last century. Future population genomic investigations based on whole genome resequencing, with increased sample sizes and marker numbers that allow the exploration of structural variation and coalescence times, will help refine understanding of demographic histories. These will provide improved resolution of recent fluctuations in effective population size, admixture and migration events. Such data will enhance evaluation of conservation status by providing more comprehensive population differentiation metrics, inbreeding coefficients, and better quantification of genetic load and inbreeding depression potential.

PSMC analysis revealed broadly similar *N_e_* trajectories among Galápagos iguanas during the mid–late Pleistocene, consistent with the relatively stable climatic conditions in the equatorial islands during the period^2^. The subsequent decline towards 100 Kya may have been driven by increased climatic variability and sea-level changes during the onset of the last glacial cycle, which could have altered habitat availability, configuration and connectivity among islands^54^. The recent increases in *N_e_* for *A. cristatus* and *C. subcristatus* could reflect transient expansions during late-glacial environmental shifts or secondary contact among island populations^2^, although recent dynamics are difficult to resolve with PSMC. Divergence among species trajectories may reflect differences in resource dependence and island-specific histories of colonisation and habitat turnover. Caution must be exercised when interpreting the timing and magnitude of predicted changes and absolute values of *N_e_*, since PSMC estimates become less reliable in the very ancient and recent past^55^.

Positive selection signatures in the Galápagos iguanas are consistent with lineage-specific adaptations. We identified a larger proportion of genes under positive selection in *A. cristatus*, likely associated with the transition from a land-dwelling ancestor to the marine environment. In other terrestrial species adapting to aquatic niches, traits experiencing selection include apnea, hypoxia tolerance, biomechanical and energetic adaptations, elaborated cardiovascular control and maintenance of body water balance^56^. Selection scans of air-breathing marine vertebrates with highly developed dive responses, or those adapted to hypoxic conditions at high altitude, have detected similar sets of positively selected genes to those identified in *A. cristatus*. The master regulator of oxygen homeostasis, the Hypoxia-Inducible Factor (HIF), is a notable target of hypoxia adaptation in highland species including Andean humans and Tibetan horses, sheep and wolves^57,58,59^. We found both *HIF-1α* and *HIF-2α* (*EPAS1*) to be under positive selection in *A. cristatus*. In addition, we identified positively selected genes linked to osmotic and cardiovascular regulation, haemoglobin metabolism and oxygen binding and transport. Molecular dynamics simulations suggest that the oxygen affinity of the haemoglobin tetramer Hb(αA)₂(βII)₂ is higher in the marine iguana than in the land species ^60^. Such genes are also under positive selection or differentially expressed in Tibetan human populations, sea snakes, alpine sheep, highland geladas and northern elephant seals^56,61,62^.

*A. cristatus* is the only vertebrate known to repeatedly shrink and grow its body size as a strategy to manage energetic overheads during resource limiting conditions^8^. This morphological plasticity starts with body size reductions during El Niño events, caused by bone mass reduction via resorption and loss of cartilage and connective tissue, and is followed by regrowth once favourable conditions are restored^9^. Similar reductions in bone density and strength are reported in astronauts and osteoporotic mammals^63^, whereas bone and cartilage growth have been widely described in amphibian and reptilian limb regeneration^58^. Genes under positive selective pressure and with putatively high-impact functional variants in the marine iguana have known functions related to growth regulation, bone homeostasis and muscle development in vertebrates. Furthermore, we identified genes under positive selection that are involved in nutrient sensing and stress hormone release. Glucocorticoids are the main stress hormones secreted as a response to starvation and have marked effects on bone metabolism by enhancing bone resorption and decreasing bone formation and differentiation of osteoblastic cells^64^. Severe environmental conditions during El Niño have been documented to increase glucocorticoid concentrations in marine iguanas due to food shortages^65^. We uncover key genes potentially involved in this remarkable adaptation, which is driven by nutrient availability and induced by the dysregulation of energy metabolism and signalling cascades. Such genes may be promising candidates to explore in future as biomedical targets for tissue regeneration and bone density therapies.

Our analyses reveal signatures of positive selection in multiple genes involved in DNA damage sensing and signal transduction. DNA damage sensors, such as the positively selected genes encoding ATR and DNA-PKs, detect genotoxic stress and coordinate downstream responses involved in DNA repair and cell cycle regulation ^18^. The positively selected *MAPK14* plays a central role in the MAP kinase signal transduction pathway, which mediates signal transduction cascades and modulates cellular responses to extracellular stimuli, including UV radiation^66^. We identified unique sequence variation in the MAPK insert and an extended N-terminus in *MAPK14*, fixed in the in Galápagos iguana lineage, altering its interaction with upstream kinases *MAP2K3* and *MAP2K6* and potentially affecting cell stress signalling^67^. Postive selection was identified in other key genes involved in DNA repair mechanisms and tumorigenesis, including important targets of cancer therapies such as *BRAF* (a frequently mutated oncogene in melanomas). *C. marthae* has skin patches lacking protective melanin, but experiences increased UV intensity (up to 600 µW/cm^2^) at altitudes exceeding 1,700m on Wolf Volcano^9^. The pink skin areas are devoid of pigment cells and result from blood flowing through a network of superficial dermal vessels (Fig. 7a)^68^. This depigmentation leads to higher rates of erythrocyte nuclear DNA damage in response to UV radiation compared to the other Galápagos iguana species^69^. UV induced DNA damage is a risk factor for melanoma development, but to our knowledge, Galápagos iguanas have no reported incidences of cancer. It is possible that the observed variation in these genes may contribute to increased resilience to UV damage and cancer resistance, evolved in response to high UV exposure while basking. Our *in silico* structural analyses indicates that variation in Galápagos *MAPK14* sequences has potential to be functionally relevant. Determining actual functional effects, mechanisms and any potential wider biomedical relevance will require extensive further experimental work. These results add to previously identified adaptive signatures reported for DNA repair and cancer suppression related genes in species including the naked mole rat (*Heterocephalus glaber*), bowhead whale (*Balaena mysticetus*), elephants (*Loxodonta* and *Elephas* sp.) and Galápagos tortoises (*Chelonoidis* sp.) ^70,71^.

Our analysis revealed species-specific signatures of positive selection in key pigmentation genes. *A. cristatus* are generally black and have a thick epidermal layer packed with melanocytes, which is reflected in the number of positively selected genes involved in melanocyte development and melanin synthesis and transport (Fig. 6). In addition, males can present hues of red and green during the reproductive season, which could be associated with the positive selection in key carotene pigment metabolism genes. In land iguanas, genes involved in xanthophore differentiation, pteridine synthesis and carotenoid metabolism were under positive selection. Xanthophores are pigment cells that contain carotene and pteridine pigments, which result in yellow to red colouration. Previous studies confirmed the presence of carotenoids in *Conolophus* blood, supporting their role in skin colour ^72^. Structural analysis revealed that amino acid substitutions in *MC1R* are consistent with a partial reduction in the receptor’s signalling efficiency in land iguanas (Fig. 7b,c). In particular, the R67Q substitution disrupts a conserved intrachain hydrogen bond and alters the Gs-beta subunit contact interface, while the V181I substitution perturbs the loop geometry adjacent to ILE180, the direct contact residue with the *α-MSH* pharmacophore. Fixed substitutions in *MC1R* are consistent with reduced eumelanin and increased pheomelanin production, in line with the transition from black marine to yellow-orange land iguanas (Fig. 7 d,e). Analogous cases have been reported in other reptiles, including experimentally validated *MC1R* substitutions in the V181I of *Phrynocephalus erythurus* at position 183, immediately adjacent to the *Conolophus* mutation, which produced lighter pigmentation^73^. The A186P and A234P substitutions introduce proline residues near the α-MSH binding site and the Gα-binding interface, respectively, where the constrained backbone geometry of proline is expected to rigidify and distort the local loop conformation, altering receptor-ligand and receptor-G protein coupling^74^. Proline substitutions at homologous functional positions have documented pigmentation effects in other vertebrates by altering α-MSH responsiveness, as confirmed by direct functional assays^74,75^. Additionally, intracellular loop mutations in *MC1R* are established drivers of pigmentation change in vertebrates, suppressing MITF transcription and limiting signal transduction rate by impairing stimulatory G protein coupling^76^.

*C. marthae* is one of the world’s most distinctive reptiles owing to its unique pink colouration. Hatchlings exhibit a maculated pattern on a green background as an anti-predatory adaptation, although this pigmentation is lost during ontogeny^77^. Our study provides evidence that the pink iguana phenotype has evolved through coordinated modifications at multiple nodes of the melanocyte differentiation programme acting in concert with altered *MC1R* signalling. First, positive selection found in *TCF7* and *JAG1*, regulators of *MITF* transcription via the independent Wnt/β-catenin and Notch signalling axes respectively, suggest convergent suppression of the master melanocyte transcription factor from multiple upstream inputs. Second, positive selection and gene family expansion in *MYO1F* may have altered melanosome transport, while positively selected *GMPS* modulates GTP pools available for stimulatory G GTPase activity, representing potential downstream and upstream modifiers flanking the *MC1R* signalling node. Third, selective pressure on *MYH11*, a smooth muscle myosin, may have contributed to the characteristic increased vasculature in the histological dermal arrangements of the pink iguana (Fig. 7a)^69^. Collectively, this provides the first genomic and protein structural evidence for the basis of pigmentation loss in the pink iguana, resulting from reduced *MC1R* signalling efficiency, convergent suppression of MITF and altered melanosome transport, along with increased dermal vasculature enhancing pink colouration.

Interactions between different selection pressures and trade-offs, including sensitivity to UVB-induced damage, thermoregulation, vitamin D metabolism, habitat and mate choice are likely to influence the evolution of the land iguana pigmentation traits^49^. Recently, Garizio et al.^78^ suggested that *C. marthae* avoids ecological competition with *C. subcristatus* on Volcan Wolf by preferentially using microhabitats with higher vegetation cover, offering greater shade and lower UVB exposure. This imposes a thermoregulatory cost, which *C. marthae* offsets through depigmentation and increased dermal vascularisation to enhance heat exchange efficiency, at the expense of increased UVB sensitivity.

However, this raises a question of evolutionary directionality: did this habitat shift relax selection on pigmentation allowing the pink phenotype to arise through drift, or did an emerging pink phenotype facilitate behavioural and ecological adaptation? The complex genomic architecture of positive selection across multiple pigment regulatory pathways and gene family expansion in *C. marthae* strongly favours that the pink phenotype evolved through selection rather than drift, and preceded or co-evolved with the ecological displacement. While the basal, putative, DNA repair adaptations (e.g. *MAPK14*, *BRAF* and *RAD51*) may partially offset the *MC1R-*mediated reduction of melanin-associated photoprotection in the other land iguana species, the additional UVB vulnerability imposed by the pink phenotype ultimately necessitated a compensatory shift to sheltered habitats, driving a complex eco-physiological trade-off.

Overall, we provide a foundation for future population genomics research and conservation genomic management of these endangered reptiles. We contribute to the broader understanding of the genomic makeup of squamates, which remain relatively underrepresented in genomic datasets compared to other vertebrates. Our analyses showcase how sequencing data can be leveraged to reconstruct evolutionary and demographic histories as well as to identify the molecular basis of adaptive traits, some of which may be of broader biomedical interest. Finally, our results provide evidence for the need to implement strong conservation measures to reduce the risk of erosion of genetic diversity in Galápagos iguanas.

## Materials and Methods

### Sampling and Extraction

Samples were collected from a single individual of *Amblyrhynchus cristatus* on Plaza Sur, *Conolophus pallidus* on Santa Fe, and both *C. marthae* and *C. subcristatus* on Wolf Volcano in Isabela. *Cyclura carinata* was sampled on Little Water Cay in the Turks and Caicos Islands. *Ctenosaura bakeri* blood was collected from an individual housed in the Bioparco di Roma in Italy that originates from the island of Útila in Honduras. The *Iguana delicatissima* sample was obtained from an individual housed at Rotterdam Zoo in The Netherlands that originates from the island of Sint Eustatius. Briefly, blood was collected from the caudal vein of adult males and preserved in a near-saturated ammonium sulphate solution buffered with sodium citrate and EDTA acid.

Blood lysates were centrifuged at 1,000 rpm for 3 minutes and the transparent supernatant was discarded. High molecular weight DNA was extracted for all species from the cell fraction using a phenol-chloroform protocol ^79^. In addition, RNA was extracted for the marine iguana using the Direct-zol RNA MiniPrep kit (Zymo Research, Irvine, CA, USA) following the manufacturer’s protocol and eluted in a final volume of 50 μl.

### Sequencing

Illumina paired-end and mate-paired sequencing of *A. cristatus* was carried out at the Malaysia Genome and Vaccine Institute (Kajang, Malaysia). Paired-end libraries were prepared based on 500 bp fragment size using Illumina TruSeq PE kit, and mate-paired libraries were prepared from 3 kbp and 8 kbp fragments using Nextera XT kit. Marine iguana libraries were sequenced on an Illumina HiSeq 2000. Illumina paired-end sequencing of *C. pallidus, C. marthae, C. subcristatus, C. carinata, C. bakeri* and *I. delicatissima* was performed at Novogene (Cambridge, UK) on an Illumina NovaSeq 6000, with paired-end libraries prepared based on 500 bp insert size using a NEBNext® DNA Library Prep. The sequencing library for long read sequencing was constructed using the SQK-LSK109 kit from Oxford Nanopore Technologies. Long-read ONT sequencing was performed at the Next Generation Sequencing Facility at the University of Leeds for *A. cristatus, C. pallidus, C. marthae, C. subcristatus, C. bakeri* and *I. delicatissima* on R9.4 minION flowcells using a MinION sequencer, and the raw fast5 data was basecalled with Guppy v.3.2.10 (Oxford Nanopore Technologies, UK). Additionally, RNA libraries for the marine iguana were constructed using a NEBNext® Ultra™ Directional RNA Library Prep Kit and sequenced on a NovaSeq 6000 at Novogene (Cambridge, UK).

### Read quality assessment and filtering

Short and long read sequences were generated for all iguana species except for *C. carinata*, for which only Illumina short read sequences were produced. Short-read quality was examined using FastQC v.0.11.9^80^. We used Trimmomatic v.0.39^81^ to remove lower quality reads with the following flags “-phred 33 ILLUMINACLIP:TruSeq3-PE.fa:2:30:10 SLIDINGWINDOW:4:30 LEADING:30 TRAILING:30 MINLEN:50”. Long-read quality was analysed using NanoPlot^82^. Long ONT reads were left untrimmed as this resulted in higher quality assemblies.

### Genome assembly

Eleven short, hybrid and long genome assembly pipelines were tested under a variety of assembly parameters for each method. For short-read assembly, MaSuRCA v.4.0.2^83^, SOAPdenovo v.2.04^84^ and Sparse v.20160205^85^ were tested. The hybrid assemblers that incorporate short and long read data used were dbg2olc v. 20180222^86^, MaSuRCA v.4.0.2 ^83^ and Wengan v.0.2^87^, as well as the long-read assemblers Flye v. 2.8.3^88^, MECAT v.1.0^89^, Miniasm v.0.3^90^, Shasta v.0.7.0^91^ and wtdbg2 v.2.5^92^. The output from the best performing long-read assembler Flye was polished with short reads using Racon v.1.4.11 ^93^ and Pilon v.1.23^94^, as well as with long reads using Medaka v.1.2.3 (https://github.com/nanoporetech/medaka). The genome assembly statistics for all pipelines tested can be found in Supplementary Table S1.

### Genome annotation

Repetitive elements were identified using RepeatModeler v.2.0.2a^95^. Custom repeat libraries were created for each species and were combined with repeat sequences from *Anolis carolinensis* and were classified using the module RepeatClassifier. Genomes were soft masked with RepeatMasker v.4.1.2 (http://www.repeatmasker.org/) using the repeat libraries. BRAKER v.2.1.6^96,97^ was used to predict protein-coding genes using RNA-seq data and protein homology information. Firstly, gene prediction was performed in BRAKER1 using RNA-seq data from marine iguanas, including the newly generated data and publicly available Ion Torrent RNA-seq reads (NCBI SRA at BioProject PRJNA602224). The RNA-seq datasets were aligned to the soft masked genomes using STAR^98^. Given the high alignment rate of marine iguana reads to the other iguanas’ genomes (>85% uniquely mapped reads) and the lack of RNA-seq data for the other species, this data was used as evidence for this gene prediction step in all species. Secondly, an independent gene prediction was run in BRAKER2 using protein homology information from the OrthoDB v.10 database^99^. Subsequently, the two gene predictions were integrated using TSEBRA v.1.0.3^100^ with default settings. TSEBRA selects transcripts from the BRAKER1 and BRAKER2 annotations to generate a joint prediction based on both RNA-seq and protein evidence.

Functional annotation of the protein-coding genes was performed combining the search for full-sequence similarity using BLAST against the NCBI squamate protein database and targeted characterisation of functional elements through the InterProScan pipeline^101^. The BLAST and InterProScan outputs were loaded onto the genome annotations using the script agat_sp_manage_functional_annotation.pl from the AGAT package (https://github.com/NBISweden/AGAT). To obtain ‘high-confidence’ sequences from the gene annotations, Diamond v.0.9.24^102^ searches were conducted against the *Anolis carolinensis* reference proteome, the UniProtKB/Swiss-Prot database, and the UniProtKB/TrEMBL database (The UniProt Consortium, 2021). The BUSCO^103^ scores were computed in the ‘protein’ mode for each of the subsets derived from the unique matches obtained when searching against the three databases. The subsets with the highest BUSCO scores across all species corresponded to the unique matches against the UniProtKB/TrEMBL database.

### Mitogenome assembly and annotation

*De novo* assembly of the mitochondrial genome was performed with the adapter-trimmed Illumina sequencing reads using the GetOrganelle pipeline^104^. GetOrganelle was run under default options with a range of k-mers of 21, 45, 65, 85 and 105. Mitogenome annotation and visualisation was performed using the MITOS WebServer^105^.

### Transcriptome assembly and annotation

Both *de novo* and reference-based transcriptome assemblies were performed for *A. cristatus*. For the reference-guided assembly, the RNA-seq reads were mapped onto the genome assembly of *A. cristatus*. The two assembly strategies were performed using Trinity v.2.14.0^106^ under default settings with the ‘trimmomatic’ flag. Publicly available raw Ion Torrent RNA-seq reads from heart, muscle, blood, skin and lung tissues of a marine iguana from Genovesa island were also added to the dataset (NCBI SRA at BioProject PRJNA602224). The *de novo* and genome-guided, trimmed and untrimmed assemblies were performed on both the newly generated data and on the concatenation with the publicly available reads. The assemblies were filtered to remove redundant and poorly constructed transcripts using TransDecoder v.5.5.0 (https://github.com/TransDecoder/TransDecoder) and CD-Hit v.4.8.1^107^.

The transcriptome was annotated using Trinotate v.3.2.2^108^. Coding sequences were translated into amino acid sequences using TransDecoder v.5.5.0. Trinotate used BLAST^109^ for homology search and HMMER3^110^ for sequence feature annotation. rnammer v.1.2^111^ was run for RNA classification, SignalP v.5.0^112^ for signal peptide identification and tmHMM v.2.0c^113^ for the prediction of transmembrane domains. The databases queried were the UniProtKB/Swiss-Prot database (The UniProt Consortium, 2021), the Pfam protein family database^114^, the SignalP signal peptide database, the eggNOG database of nested orthologous gene groups^115^, the tmHMM transmembrane domain database, the Gene Ontology (GO) knowledgebase (The Gene Ontology Consortium, 2021) and the Kyoto Encyclopedia of Genes and Genomes^116^ (KEGG).

### Ortholog identification and filtering

The predicted proteomes or coding sequences of all squamate reptile species with assembled genomes available in April 2023 were obtained (Supplementary Table S13). The coding sequences were ‘cleaned’ (to remove those sequences that were not divisible by 3) and translated into amino acid sequences using the vespa_clean.py and vespa_translate.py commands from VESPA^117^. The longest transcript variant per gene was extracted for each proteome using the custom script *p_LongestIsoformInFastaFile.py*. OrthoFinder v.2.5.4 was used to infer orthogroups among the squamate dataset^118^.

The 439 SGOs identified were filtered to obtain true orthologs with informative signals. Firstly, alignments were constructed for each single-gene orthogroup using MAFFT v. 7.505^119^, MUSCLE v.5.1^120^ and PRANK v. 170427^121^ using default settings. The best fitting alignment per orthogroup was selected based on the highest normalized mean distance score^122^ and the lowest number of gaps. The alignments were then trimmed using TrimAL v. 1.4.1^123^ with the ‘automated1’ flag. Sequence saturation was calculated using the ‘saturation’ function in PhyKIT^124^ and orthogroups with values below 0.8 were removed resulting in 401 filtered orthogroups. Clan_check (https://github.com/ChrisCreevey/clan_check) was used to reduce noise from hidden paralogy and genes that violated uncontroversial splits at the suborder level were filtered out and not used in further analysis. A total of 311 orthogroups were retained and the 90 single gene families containing more than one clan violation were discarded. Likelihood mapping was performed using the command ‘lmap’ in IQ-TREE^125^ to remove genes with insufficient phylogenetic signal. In total, 231 single gene families that contained sufficient signal ccording to this metric were retained. Lastly, compositional heterogeneity was assessed for each orthogroup using a chi-squared test in IQ-TREE. Ortholog filtering resulted in a dataset of 229 SGOs across 28 squamates.

### UCE harvesting and filtering

UCE loci were harvested from the genome assemblies of 28 squamate taxa (Supplementary Table S13). PHYLUCE v.1.7.3^126^ was used to match the Tetrapods-UCE-5Kv1 bait set (https://www.ultraconserved.org/) to the genome sequences and generate a database using ‘phyluce_probe_run_multiple_lastzs_sqlite’, with minimum identity and coverage thresholds of 80%. Matching loci were extracted using ‘phyluce_probe_slice_sequence_from_genomes’, retaining 400 bp of flanking sequence. Extracted contigs were then re-matched to the probe set using ‘phyluce_assembly_match_contigs_to_probes’ to build a relational database of UCE loci. Loci present across taxa were identified using ‘phyluce_assembly_get_match_counts’, and corresponding sequences were extracted with ‘phyluce_assembly_get_fastas_from_match_counts’. Individual loci were aligned with MAFFT^115^ via ‘phyluce_align_seqcap_align’ and trimmed using Gblocks (phyluce_align_get_gblocks_trimmed_alignments_from_untrimmed) to remove poorly aligned regions^127^.

From these alignments, 75 and 95% completeness matrices were constructed. The 75% matrix comprised 4,663 loci represented in 21 - 28 taxa with a mean length of 866 bp (519-1502 bp range) and a mean of 333 informative sites per UCE. The 95% subset contained 4,153 loci with 26 - 28 taxa with a mean length of 868 bp (519 - 1502bp) and a mean of 329 informative sites per locus. Finally, UCE loci were ranked using the R package genesortR^128^ under both default settings and a systematic-bias-only filtering scheme to identify loci predicted to carry greater phylogenetic signal while minimizing potential bias. The gene properties applied in the systematic-bias-only filtering were average pairwise patristic distance, compositional heterogeneity, level of saturation and root-to-tip variance. For each completeness matrix, the full unfiltered matrices as well as subsets of the top 500 and top 1,000 loci generated under both filtering criteria were selected for subsequent phylogenetic analyses.

### Phylogenomic reconstructions and divergence time estimation

Phylogenomic analyses on the squamate SGOs and UCEs were performed using maximum likelihood in IQ-TREE with automatic model selection, Bayesian analysis in PhyloBayes under the CAT+GTR model with a gamma distribution of four categories and a supertree approach in ASTRAL-III^129,130^.

Divergence time estimation was performed within a framework of Bayesian inference. Three sets of analyses were used to test the robustness of calibration choice. Strategy 1 modelled the calibrations as uniform distributions with soft upper bounds, strategy 2 modelled the nodes using a skewed-normal distribution, and strategy 3 used a truncated Cauchy distribution with a long tail. Then, ten fossil calibration points that covered most of the deep branches within the tree were identified (Supplementary Table S17). MCMCtree from the PAML v.4.10.6 package^131^ was used to obtain the posterior time estimates on the fixed tree topology produced by the phylogenetic reconstructions. Two independent MCMC chains were run with a 2,000 generation burn-in, subsequently sampling every 10 generations until 20,000 samples were collected. MCMCtree estimated the posterior mean divergence times and the 95% highest posterior density credible intervals on divergence times for the three calibration strategies. Additionally, the analyses were run without sequence data to generate the time prior, and the samples from the prior and the posterior were compared. The divergence time estimates were plotted with the ‘MCMC.tree.plot’ function from the MCMCtreeR package in R v.4.1.1^132^. We found that the choice of prior calibration distributions had an impact on the posterior estimates of divergence times. The concordance in age and substitution rate estimates between the uniform and skewed-normal distributions, along with the older estimates produced by the Cauchy distribution, suggest that the first distributions are a closer representation of the true ages of the splits.

### Genome-wide diversity and inbreeding

Genome-wide heterozygosity levels and runs of homozygosity (ROH) were computed for each iguanid using ROHan^133^. Firstly, the raw Illumina reads were mapped onto each conspecific genome assembly using BWA-mem v.0.7.17^134^. ROHan was run three independent times on the mapped bam files using three different window sizes (500 kbp, 1 Mb, 2 Mb), under default options for the other parameters.

The number of generations since inbreeding events was calculated using the formula G = 100/ 2rL, whereby G = number of generations, r = recombination rate and L = length of ROH in Mbp^135^. This model assumes that ROH length decreases over time and that recombination rate is constant across the genome. Published estimates of reptile recombination rates were used as an approximation since genome-wide recombination rates are unavailable for Iguanidae, including chicken^136^, green anole^137^ and steppe agama^52^. The minimum, average and maximum lengths of the ROH used to calculate the number of generations since inbreeding were estimated by ROHan.

### Genomic variant annotation

Mutational load was estimated for each species using SnpEff v.5.4c^138^ on variants called with bcftools v.1.22 mpileup using the call command ^139^. Annotation categories were parsed and quality filtered with the ‘QUAL > 20 & GEN[0].DP > 10’ flags using SnpSift^140^. Putative deleterious variants classified as *high* impact were retained. SnpEff defines high-impact variants as those assumed to have high disruptive effect on protein function, likely causing protein truncation, loss of function or nonsense-mediated decay, including the gain of stop codons, splice donor variant, splice acceptor and loss of start codon. For each retained site, genotypes were translated into alternative allele counts.

We implemented a block-based subsampling framework to generate replicate estimates across the genome. Variants were divided into 50 equally sized blocks, for which we computed relative mutational load as the total number of alternative alleles divided by the number of sites and the total count of predicted LoF variants. LoF variants were further partitioned by genotype class into masked load, corresponding to heterozygous genotypes, and realized load, corresponding to homozygous alternative genotypes. Pairwise differences among species were evaluated using Tukey HSD tests.

### Coalescent models of demographic history

PSMC analysis was applied to estimate the demographic history of the seven iguanid species^123^. Illumina reads were aligned to the corresponding reference genome assembly using Minimap2^141^. Variants were called with bcftools v.1.22 mpileup using the call command^139^. Annotated repetitive regions identified with RepeatMasker during the genome annotation stage were masked by excluding these intervals during diploid consensus sequence generation with bcftools v.1.22 using the view command^139^. The resulting VCF files were converted into diploid consensus FASTQ files using VCFtools^139^, setting the minimum read depth ‘-d’ to a third of the average read depth and the maximum read depth ‘-D’ to twice the average depth. The FASTQ files were then converted to a ‘psmc’ file using the PSMC command fq2psmcfa with the ‘-q 20’ flag^121^. *N_e_* was inferred across 64 atomic time intervals (4+25*2+4+6) using the ‘-p’ parameter with the options ‘-N 25 –t20 -r5’. 100 bootstrap replicates were performed for each species by splitting the ‘psmcfa’ file and randomly sampling from these regions. The model was scaled using a generation time of ten years to account for the variability within the species and a substitution rate of 7.7 x 10^-8^ per year per site per generation, based on the average neutral substitution rate estimated for squamate reptiles^142^.

### Scanning for signatures of positive selection

The squamate database generated for phylogenomic reconstruction was filtered to remove the proteomes with BUSCO scores <85%, resulting in a set of 22 species (Supplementary Table S13). OrthoFinder v.2.5.4 was run under default settings to infer orthologous groupings among the dataset^112^, which identified 21,990 orthogroups. Orthogroups representing less than seven species were discarded, as likelihood ratio tests performed with less than seven species have low power in detecting positive selection at amino acid sites^143^. Multiple sequence alignments were constructed for each orthogroup using MAFFT v. 7.505, MUSCLE v.5.1 and PRANK v. 170427. Alignments for each orthogroup were selected based on the highest normalized mean distance score and the lowest number of gaps. Multi-gene orthogroups were paralog-filtered to derive subtrees with 1:1 orthologous relationships, this was done using phylogenetic tree-based orthology inference in PhyloPyPruner with the maximum inclusion method (https://github.com/fethalen/phylopypruner). Following, the isolation of putative single-copy orthologs, including those identified by OrthoFinder and those obtained by paralogy pruning, were tested for hidden paralogy. Firstly, IQ-TREE was run with automatic model selection on a gene-by-gene basis using ModelFinder^144^ and 1,000 ultrafast bootstrap replicates. Robinson-Foulds distances between each gene tree and the canonical species tree were estimated using Clann^145^ and only genes with distances below 0.5 were retained.

The single-copy orthologs were converted to nucleotide format using the map_alignments command from the VESPA package. The orthologs were further filtered using the custom python script *p_geneFilter.py* to ensure that each gene family retained for further analysis contained: all Galápagos species, at least two non-Galápagos iguanids, and two non-iguanid species. This yielded 6,682 single-copy genes for the selective pressure analysis. Signatures of positive selection were tested on the ortholog set using the aBSREL method from the HyPhy package^146,147^. Selection was evaluated on a lineage-specific level for multiple lineages by annotating the input trees with LabelTrees (https://github.com/veg/hyphy-analyses/tree/master/LabelTrees). The foreground lineages specified were the four Galápagos species along with the three internal nodes within the Galápagos clade (Galápagos root node, land iguana node and *Conolophus subcristatus – C. pallidus* node). aBSREL adjusts p-values by applying the Holm-Bonferroni correction to account for multiple branch testing. To account for multiple gene testing, we applied an additional correction by calculating the false discovery rate (FDR) using the *p.adjust* function in R v.4.1.1, following the Benjamini-Hochberg procedure, with an FDR threshold of 0.05.

Gene family expansion and contraction were examined at the Iguanidae family level with Computational Analysis of gene Family Evolution (CAFE) v.5 using a Poisson distribution and estimating the rate of change of evolution (lambda parameter) for each species^148^.

### Functional characterisation of candidate genes under positive selection

The R library biomaRt v.2.50.3 was used to retrieve the gene and gene ontology (GO) information from the BioMart databases for the green anole, common wall lizard, mainland tiger snake and eastern brown snake^149^. The 6,682 single-copy genes were tested for functional enrichment of GO IDs, GO slims and KEGG terms. Fisher’s exact tests were performed with Bonferroni multiple test correction to estimate GO ID and GO slim term over-representation with the script *GOEnrichment.R*. REVIGO was used to summarise the GO terms based on similarity^150^. The *enrichKEGG* function from the R package clusterProfiler v.4.2.2 was used to assess whether genes were significantly enriched for a particular KEGG pathway, under default Benjamini-Hochberg adjustment of p-values^151^. Lastly, the PANTHER (Protein ANalysis THrough Evolutionary Relationships) classification system was used to classify the HyPhy input genes and positively selected genes by biological process, molecular function and protein class^152^. The positively selected genes were searched in the literature to identify genotype-phenotype associations, prioritising the functionally enriched set and genes with demonstrated functional associations with the adaptive traits in other vertebrates. Gephebase was interrogated to identify genetic variants associated with trait variation^153^. Silhouettes for the figures were taken from PhyloPic (https://www.phylopic.org/).

Molecular model of *Conolophus marthae* MAPK14-MAP2K6 complex and *Conolophus* MC1R, α-MSH and Gα complex (Supplementary Figure 4) was generated using AlphaFold3 (https://alphafoldserver.com), with high confidence in the interface between the kinases (pairwise iPTM 0.72). Image was produced using PyMOL 3.7 (http://www.pymol.org).

## Ethics declaration

Animal manipulation and blood sampling were performed according to a protocol that minimized animal stress, in accordance with the European Community guidelines and with the approval of the Galápagos National Park Directorate and the Turks and Caicos Islands Department of Environment and Coastal Resources. *Conolophus* and *Amblyrhynchus* samples were exported under permits granted by the Galápagos National Park and the CITES-Authority to GG (permit no. 050-03-PNG and 004-05-PNG; EC9119940 and 0211914 CITES EXPORT). Samples were imported under the CITES permits IT/IM/2003/MCE/02328 and IT/IM/2006/MCE/07225 to GG. *Cyclura carinata* samples were exported under permits granted by the Turks and Caicos Department of Environment and Coastal Resources and the CITES authority to GG (permit no. PLS-W-2021 8); samples were imported under the CITES permit IT/IM/2022/MCE/03130 to GG.

## Data and resource availability

The sequencing data (genomic and transcriptomic raw reads) generated in this study are available through NCBI repositories linked to BioProject accession number PRJNA1338454. The positive selection and gene family evolution results, transcriptome annotation SQL database and bioinformatic scripts are available from the Research Data Repository at the University of Leeds (https://doi.org/10.5518/1772).

## Acknowledgements

We thank the Galápagos National Park Directorate and park rangers for their invaluable support. We are grateful to Morag Raynor and Carolina Lascelles for library preparation and sequencing at the Next Generation Sequencing Facility of the University of Leeds. We thank the Rotterdam Zoo for sourcing the Lesser Antillean iguana sample. We thank the Bioparco of Rome for sourcing the Útila spiny-tailed iguana sample. JLD thanks Elizabeth Duncan from the University of Leeds for constructive feedback. JLD also thanks Bede Constantinides from the University of Oxford and Peter Mulhair from Trinity College Dublin for sharing code for the phylogenomic analysis. Bioinformatics work was performed on the University of Leeds High Performance Computing ARC4 cluster.

## Author contributions

JLD, MJO’C, GG and SJG designed the research. CP, GC, GG and GPG collected the samples. JLD and CP performed the DNA and RNA extractions. JLD assembled and annotated the reference genomes, with bioinformatics support from IMC. JLD performed the evolutionary, demographic and selection pressure analyses, with guidance from MJO’C, PG, GG and SJG. JLD and RB carried out the protein structure modelling. JLD wrote the primary manuscript, and all authors edited and revised the final version.

