## Supplementary Information for "Genomic perspectives on the origins of Galápagos iguanas and the evolution of marine specialisation, DNA damage response and pigmentation adaptations"

### Supplementary Figures

**a**

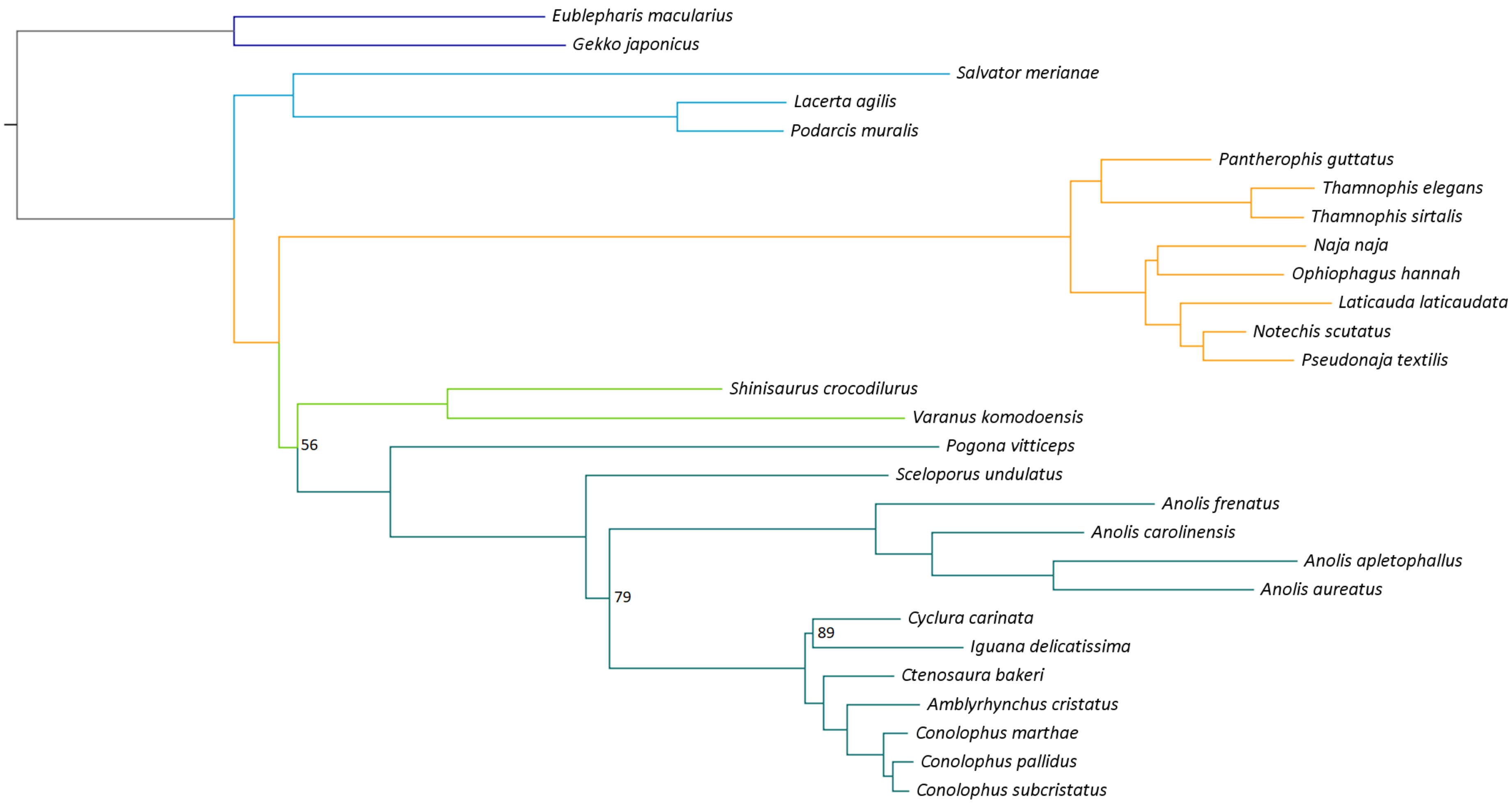
Eublepharis macularius
Gekko japonicus
Salvator merianae
Lacerta agilis
Podarcis muralis
Pantherophis guttatus
Thamnophis elegans
Thamnophis sirtalis
Naja naja
Ophiophagus hannah
Laticauda laticaudata
Notechis scutatus
Pseudonaja textilis
Shinisaurus crocodilurus
Varanus komodoensis
56
Pogona vitticeps
Sceloporus undulatus
Anolis frenatus
Anolis carolinensis
Anolis apletophallus
Anolis aureatus
79
Cyclura carinata
89
Iguana delicatissima
Ctenosaura bakeri
Amblyrhynchus cristatus
Conolophus marthae
Conolophus pallidus
Conolophus subcristatus

**b**

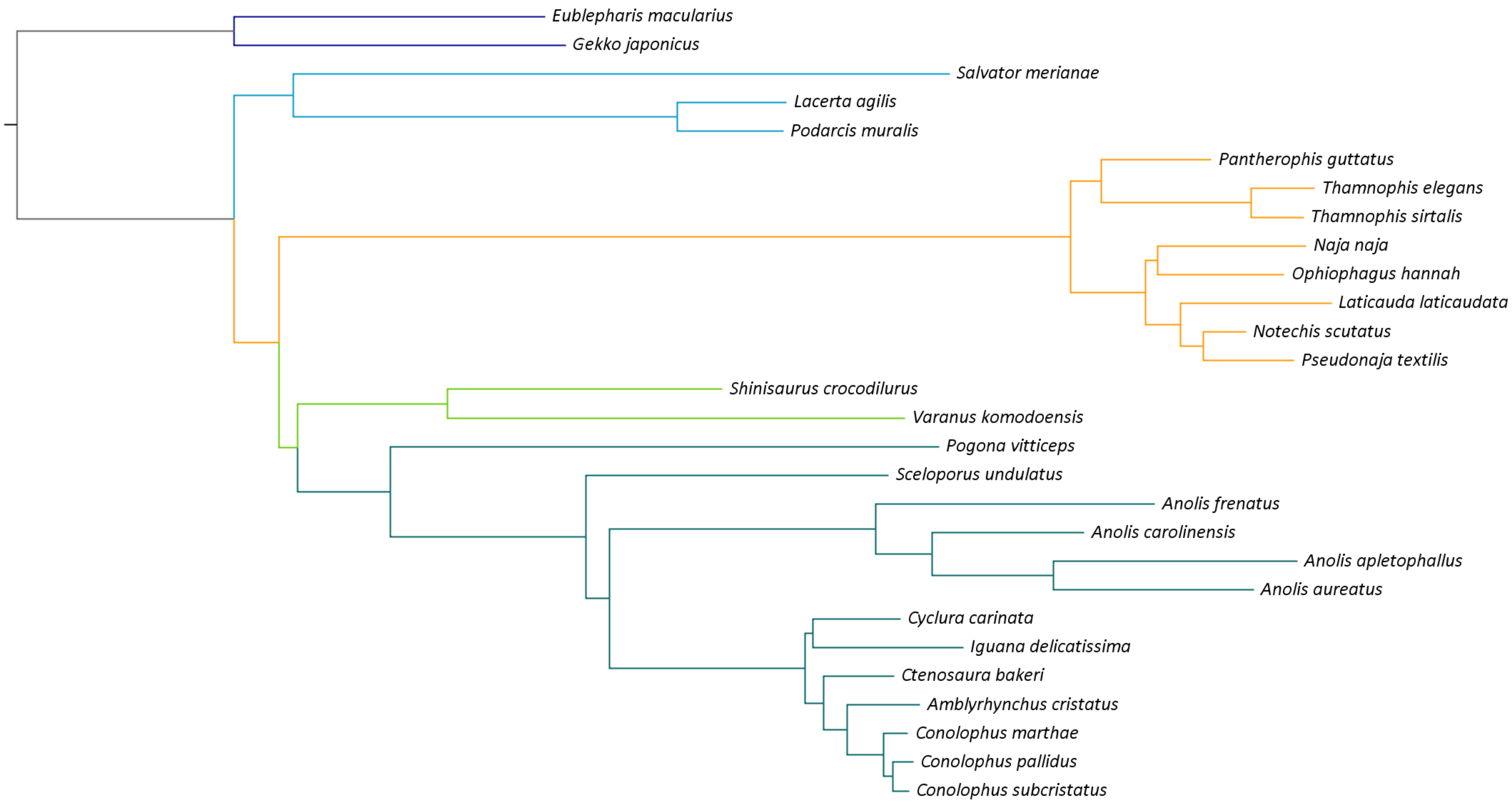
Eublepharis macularius
Gekko japonicus
Salvator merianae
Lacerta agilis
Podarcis muralis
Pantherophis guttatus
Thamnophis elegans
Thamnophis sirtalis
Naja naja
Ophiophagus hannah
Laticauda laticaudata
Notechis scutatus
Pseudonaja textilis
Shinisaurus crocodilurus
Varanus komodoensis
Pogona vitticeps
Sceloporus undulatus
Anolis frenatus
Anolis carolinensis
Anolis apletophallus
Anolis aureatus
Cyclura carinata
Iguana delicatissima
Ctenosaura bakeri
Amblyrhynchus cristatus
Conolophus marthae
Conolophus pallidus
Conolophus subcristatus

c

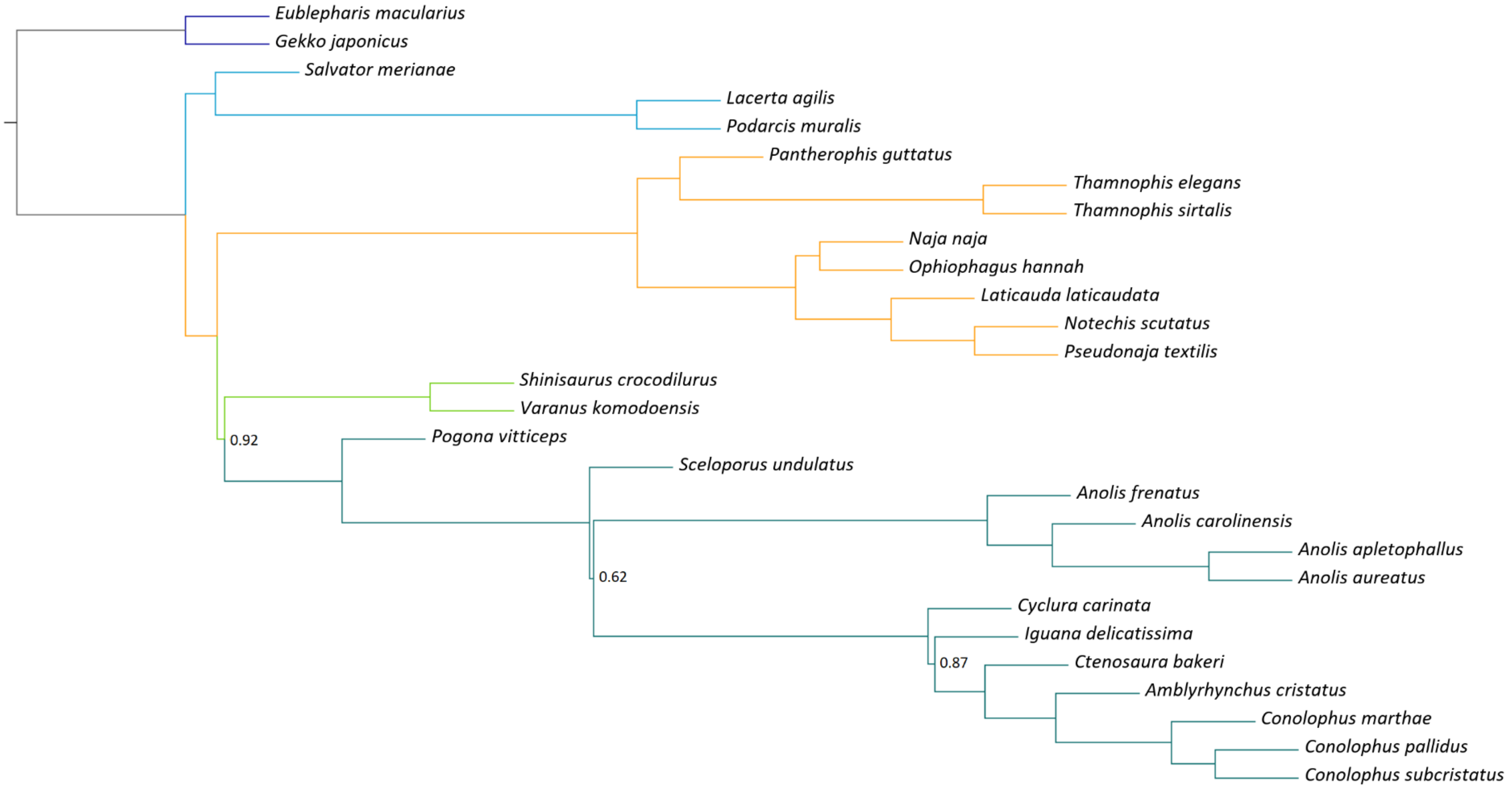


**Figure S1: Phylogenomic reconstructions for the squamate dataset with single-gene orthologs. a)** Reconstruction using maximum likelihood in IQ-TREE. Branch support is represented by ultrafast bootstrap supports from 1,000 replicates, with values equal to 100 not shown. **b)** Reconstruction using Bayesian inference in PhyloBayes. Branch support is represented by posterior probabilities; all values were equal to 1. **c)** Reconstruction using the ASTRAL supertree approach. Branch support is represented by local posterior probabilities, with values equal to 1 not shown. In all trees, dark blue represents Gekkota, light blue for Laterata, yellow for Serpentes, light green for Anguimorpha, and dark green for Iguania.

**a**

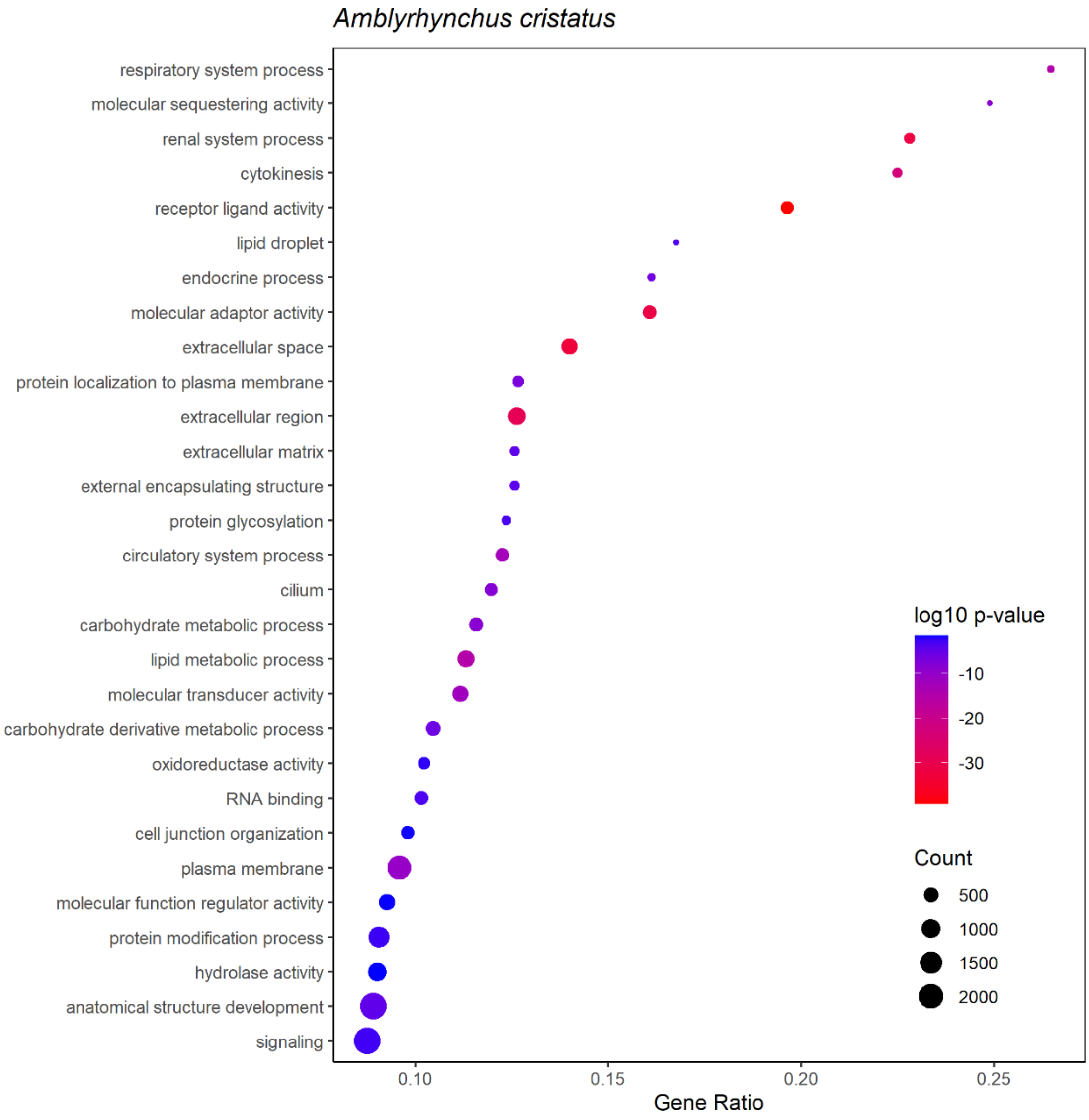

Amblyrhynchus cristatus
respiratory system process
molecular sequestering activity
renal system process
cytokinesis
receptor ligand activity
lipid droplet
endocrine process
molecular adaptor activity
extracellular space
protein localization to plasma membrane
extracellular region
extracellular matrix
external encapsulating structure
protein glycosylation
circulatory system process
cilium
carbohydrate metabolic process
lipid metabolic process
molecular transducer activity
carbohydrate derivative metabolic process
oxidoreductase activity
RNA binding
cell junction organization
plasma membrane
molecular function regulator activity
protein modification process
hydrolase activity
anatomical structure development
signaling
0.10
0.15
0.20
0.25
Gene Ratio
log10 p-value
-10
-20
-30
Count
500
1000
1500
2000

**b**

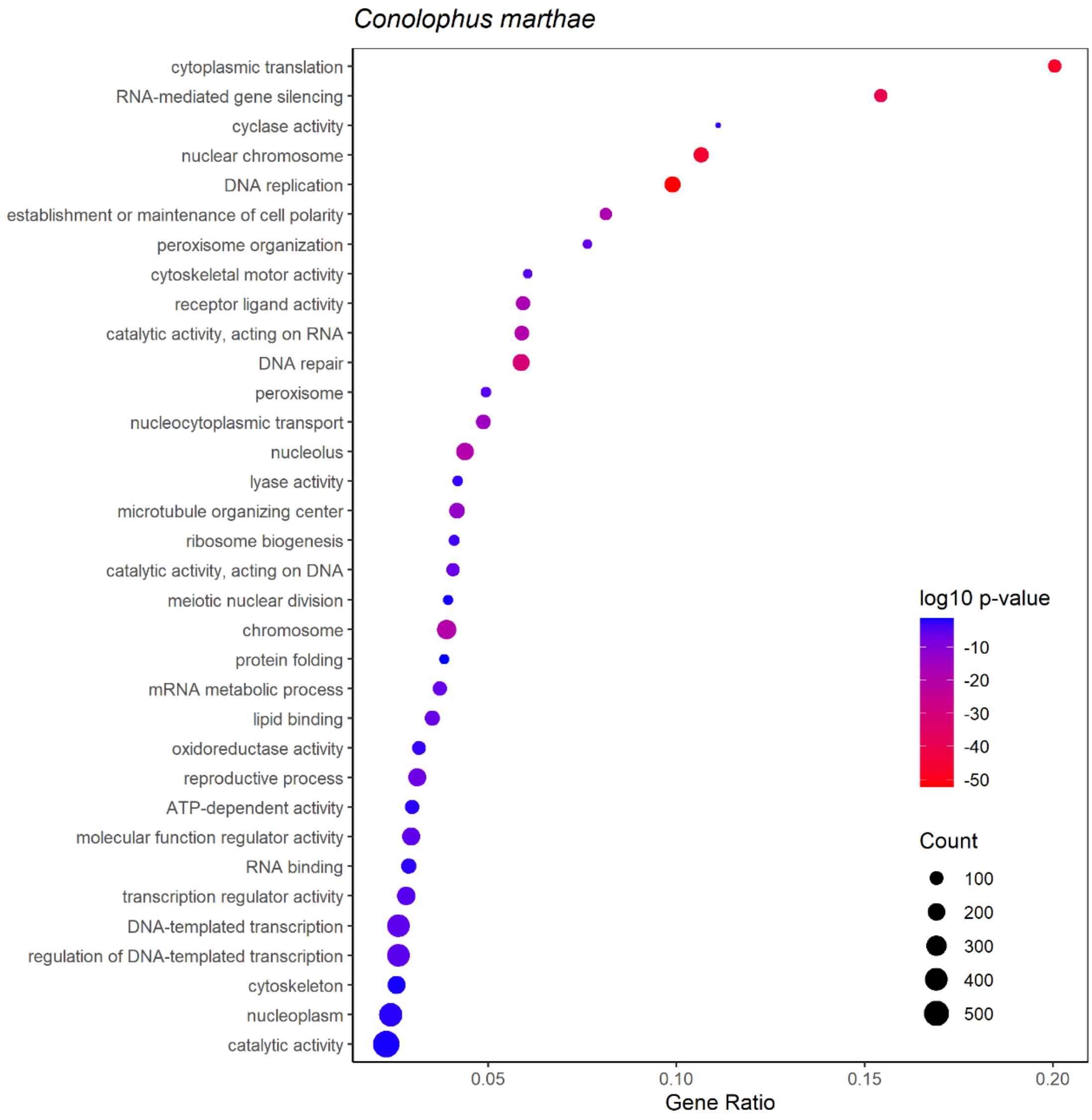
Conolophus marthae
cytoplasmic translation
RNA-mediated gene silencing
cyclase activity
nuclear chromosome
DNA replication
establishment or maintenance of cell polarity
peroxisome organization
cytoskeletal motor activity
receptor ligand activity
catalytic activity, acting on RNA
DNA repair
peroxisome
nucleocytoplasmic transport
nucleolus
lyase activity
microtubule organizing center
ribosome biogenesis
catalytic activity, acting on DNA
meiotic nuclear division
chromosome
protein folding
mRNA metabolic process
lipid binding
oxidoreductase activity
reproductive process
ATP-dependent activity
molecular function regulator activity
RNA binding
transcription regulator activity
DNA-templated transcription
regulation of DNA-templated transcription
cytoskeleton
nucleoplasm
catalytic activity
0.05
0.10
0.15
0.20
Gene Ratio
log10 p-value
-10
-20
-30
-40
-50
Count
100
200
300
400
500

**c**

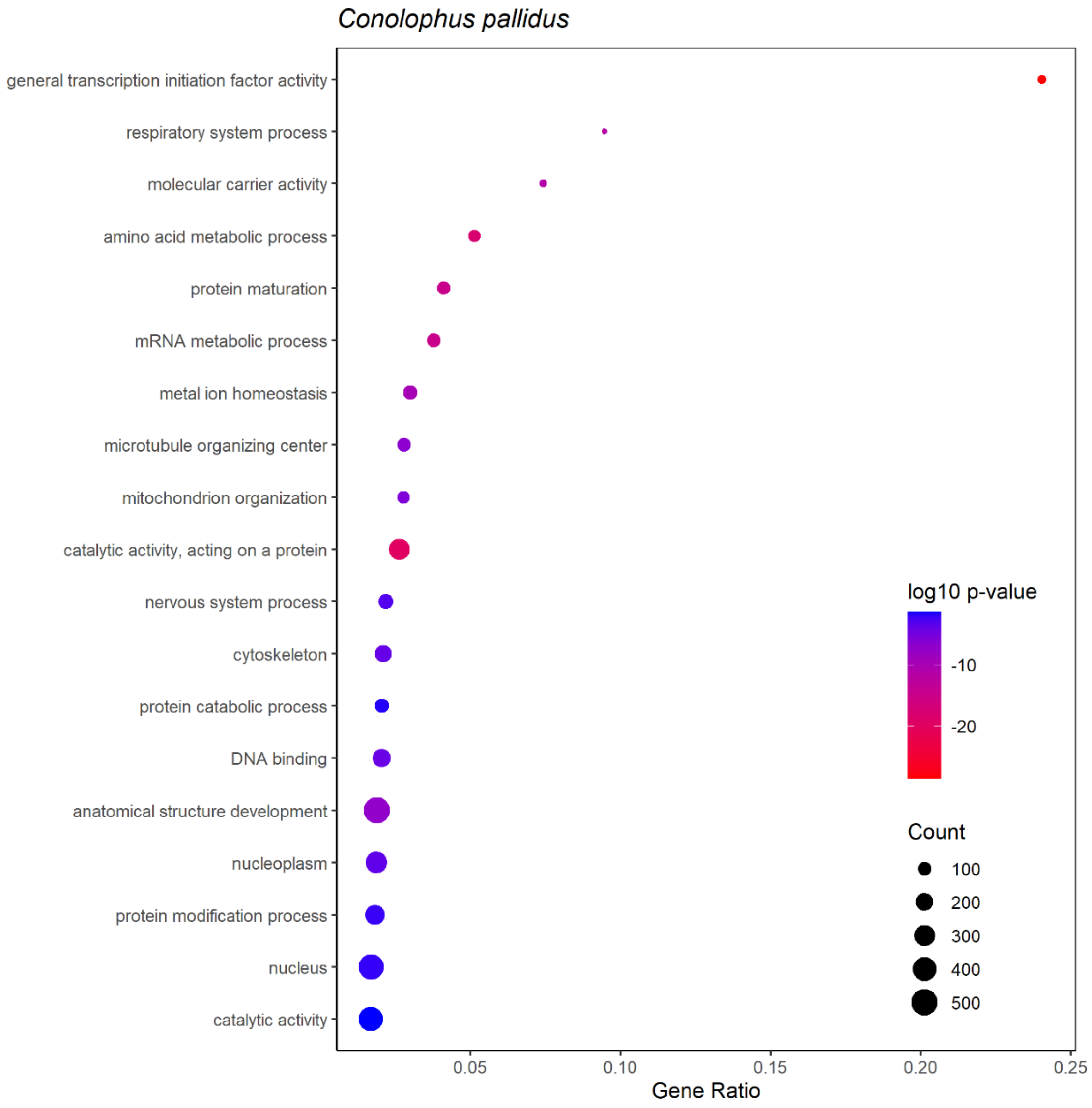

Conolophus pallidus
general transcription initiation factor activity
respiratory system process
molecular carrier activity
amino acid metabolic process
protein maturation
mRNA metabolic process
metal ion homeostasis
microtubule organizing center
mitochondrion organization
catalytic activity, acting on a protein
nervous system process
cytoskeleton
protein catabolic process
DNA binding
anatomical structure development
nucleoplasm
protein modification process
nucleus
catalytic activity
0.05
0.10
0.15
0.20
0.25
Gene Ratio
log10 p-value
-10
-20
Count
100
200
300
400
500

d

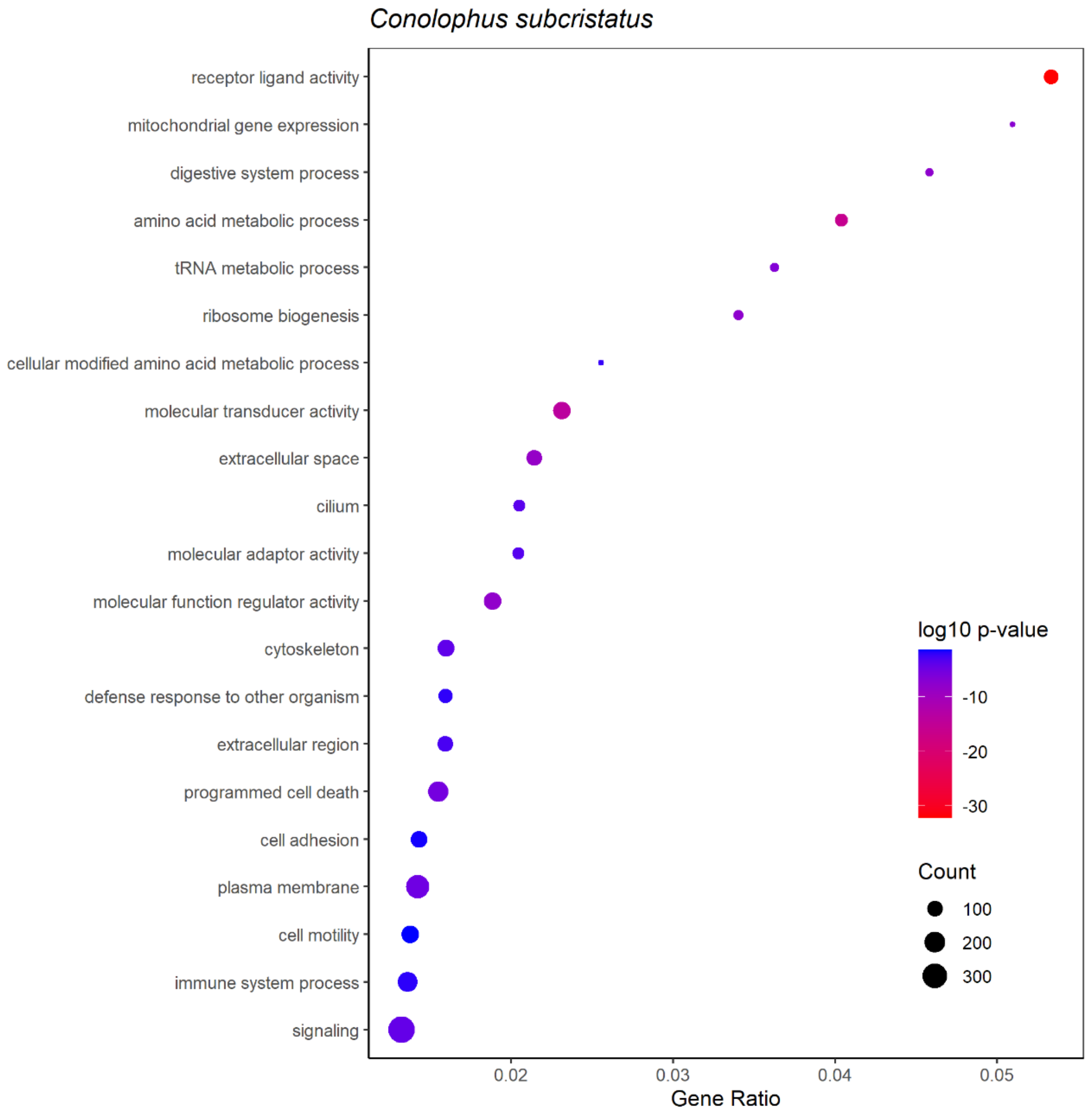

Conolophus subcristatus
receptor ligand activity
mitochondrial gene expression
digestive system process
amino acid metabolic process
tRNA metabolic process
ribosome biogenesis
cellular modified amino acid metabolic process
molecular transducer activity
extracellular space
cilium
molecular adaptor activity
molecular function regulator activity
cytoskeleton
defense response to other organism
extracellular region
programmed cell death
cell adhesion
plasma membrane
cell motility
immune system process
signaling
log10 p-value
-10
-20
-30
Count
100
200
300
0.02
0.03
0.04
0.05
Gene Ratio

e

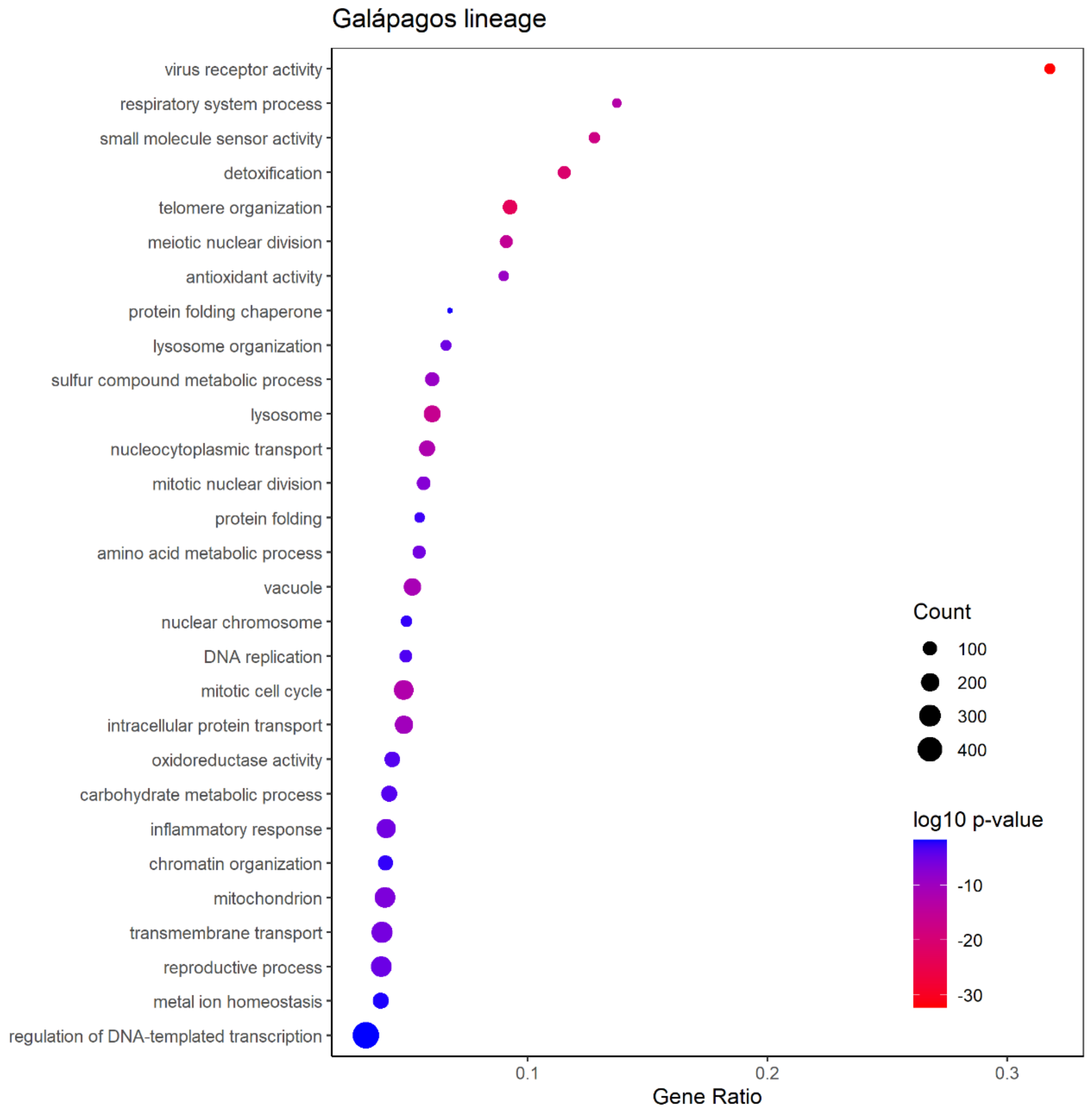
Galápagos lineage
virus receptor activity
respiratory system process
small molecule sensor activity
detoxification
telomere organization
meiotic nuclear division
antioxidant activity
protein folding chaperone
lysosome organization
sulfur compound metabolic process
lysosome
nucleocytoplasmic transport
mitotic nuclear division
protein folding
amino acid metabolic process
vacuole
nuclear chromosome
DNA replication
mitotic cell cycle
intracellular protein transport
oxidoreductase activity
carbohydrate metabolic process
inflammatory response
chromatin organization
mitochondrion
transmembrane transport
reproductive process
metal ion homeostasis
regulation of DNA-templated transcription
0.1
0.2
0.3
Gene Ratio
Count
100
200
300
400
log10 p-value
-10
-20
-30

**f**

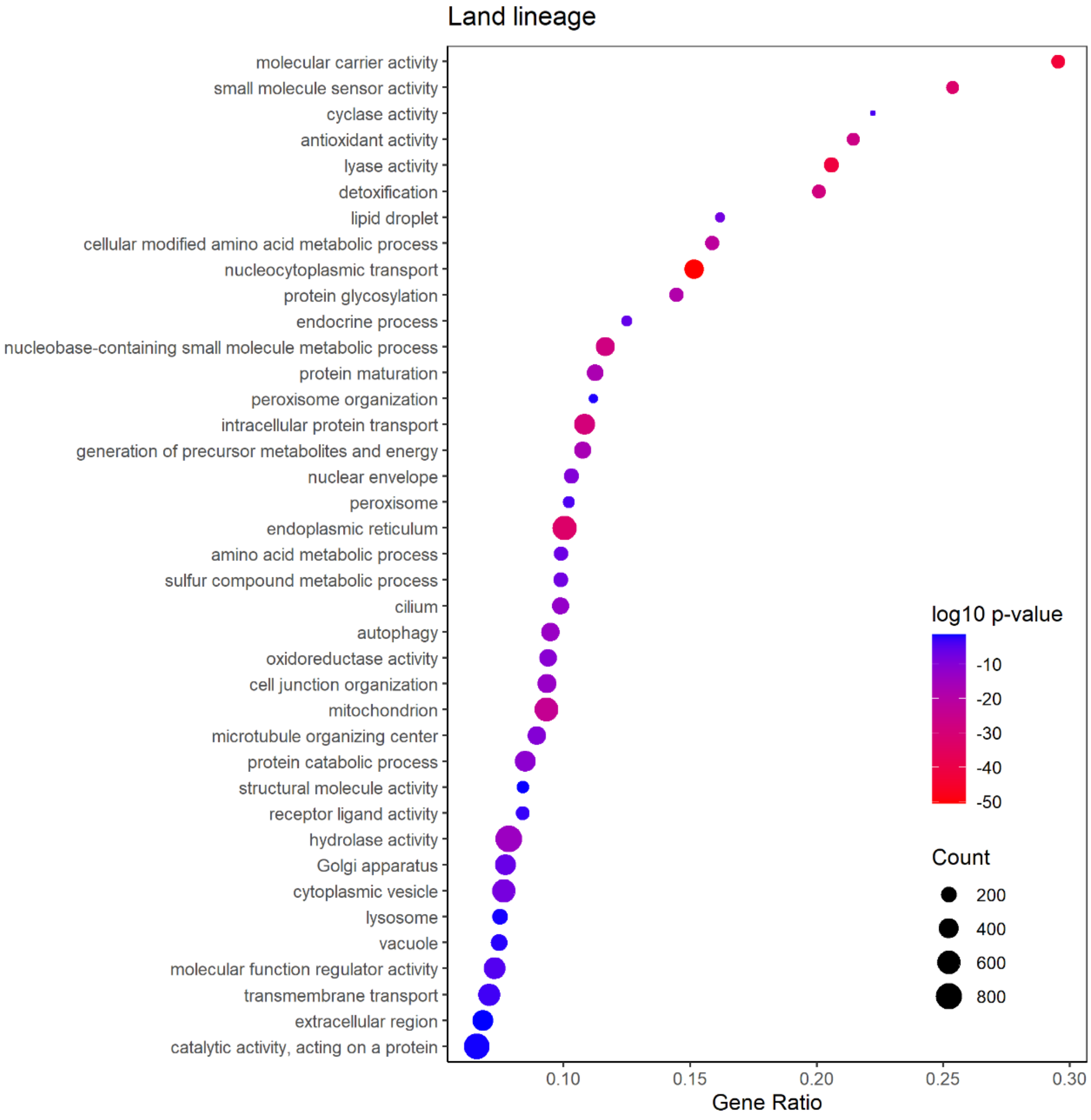
Land lineage
molecular carrier activity
small molecule sensor activity
cyclase activity
antioxidant activity
lyase activity
detoxification
lipid droplet
cellular modified amino acid metabolic process
nucleocytoplasmic transport
protein glycosylation
endocrine process
nucleobase-containing small molecule metabolic process
protein maturation
peroxisome organization
intracellular protein transport
generation of precursor metabolites and energy
nuclear envelope
peroxisome
endoplasmic reticulum
amino acid metabolic process
sulfur compound metabolic process
cilium
autophagy
oxidoreductase activity
cell junction organization
mitochondrion
microtubule organizing center
protein catabolic process
structural molecule activity
receptor ligand activity
hydrolase activity
Golgi apparatus
cytoplasmic vesicle
lysosome
vacuole
molecular function regulator activity
transmembrane transport
extracellular region
catalytic activity, acting on a protein
0.10
0.15
0.20
0.25
0.30
Gene Ratio
log10 p-value
-10
-20
-30
-40
-50
Count
200
400
600
800

**g**

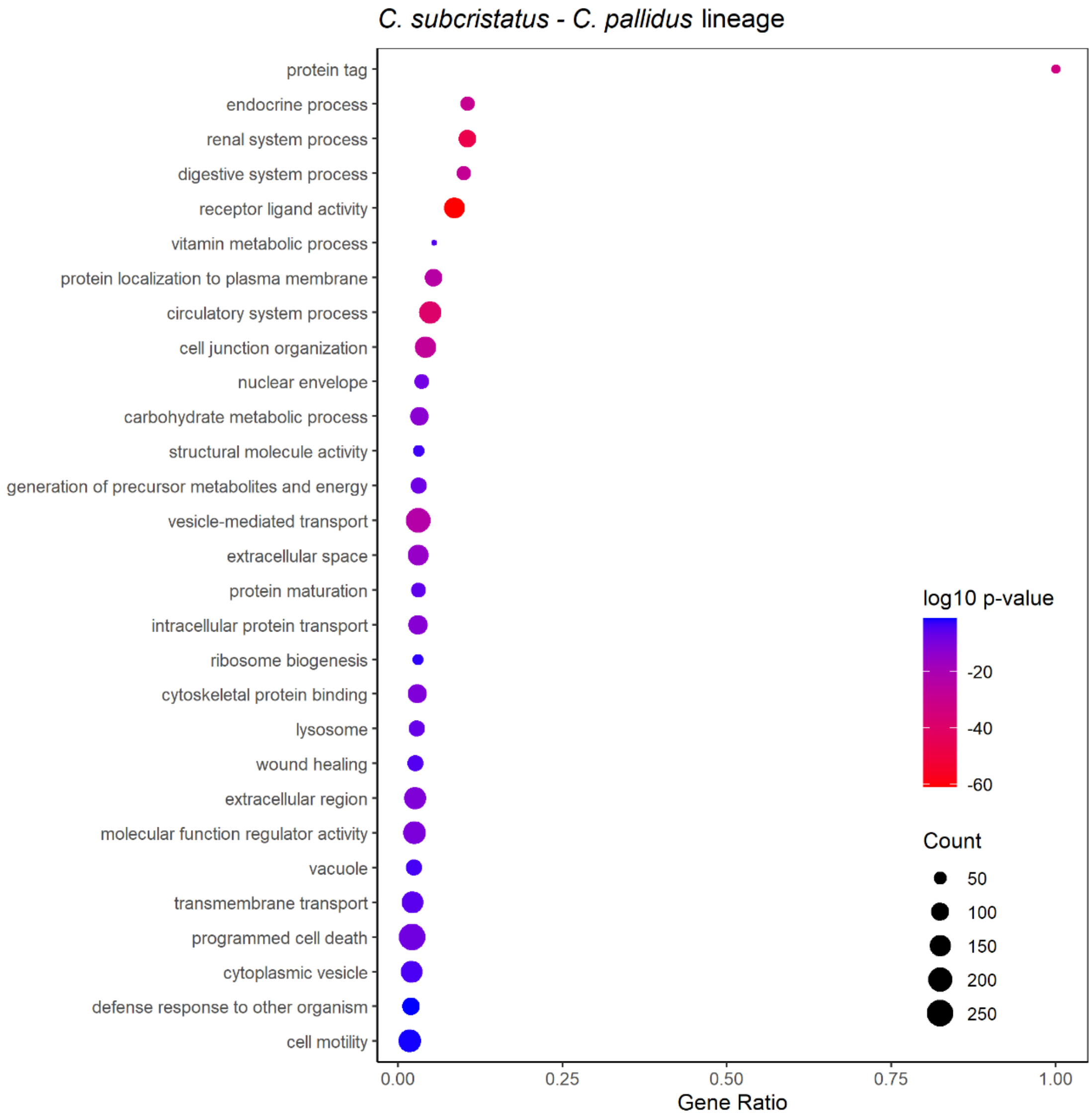


**Figure S2: Enriched GO slim terms across the positively selected genes.** GO slim term enrichment was tested for each lineage: **(a)** *Amblyrhynchus cristatus*, **(b)** *Conolophus marthae*, **(c)** *C. pallidus*, **(d)** *C. subcristatus*, **(e)** the Galápagos lineage, **(f)** the land lineage **(g)** the C. subcristatus – C. pallidus branch. 'Log10 p-value' is the log 10 of the Bonferroni-adjusted p-values. 'Count' refers to the number of RefSeq IDs enriched in a particular GO slim term. 'Gene ratio' is the occurrence of a given GO slim term in the positively selected genes set relative to the total occurrence of the term in the total gene set (including both selected and non-selected genes).

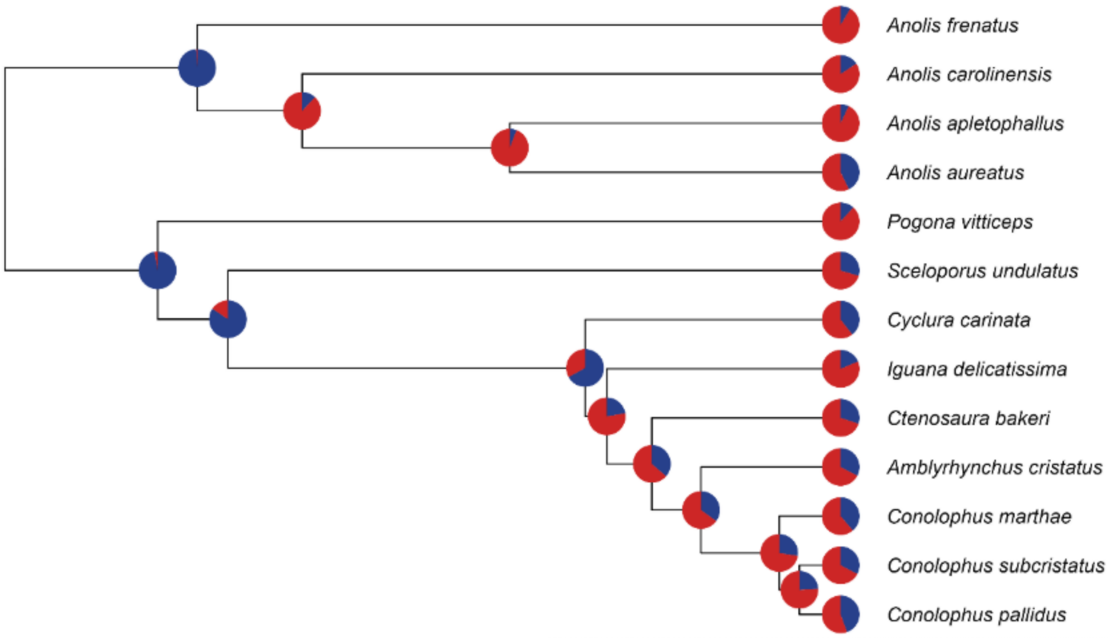


**Figure S3: Estimation of gene family expansion and contraction on each evolutionary branch.** The pie charts show the proportions of significantly expanded (blue) and contracted (red) gene families. Source data are provided in the Supplementary Data.

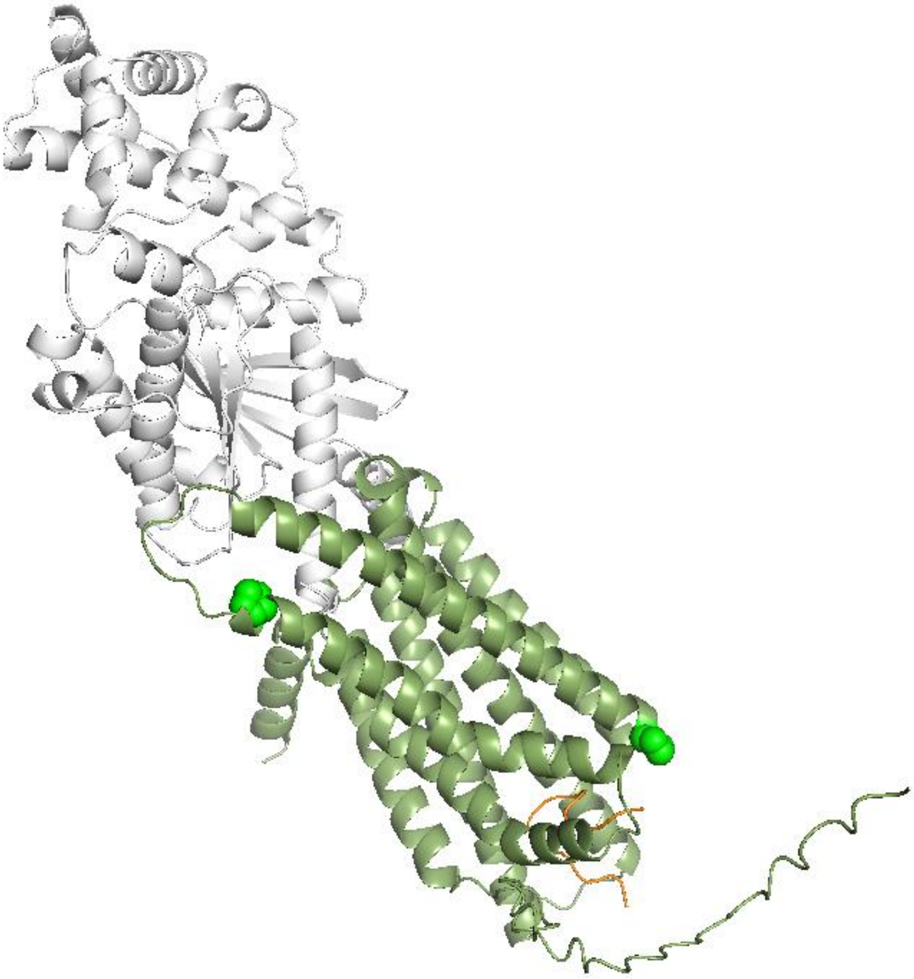

**Figure S4: 3D protein model of the MC1R (green), α-MSH (white) and Gα (orange) complex for the land iguanas, showing the A186P and A234P proline substitutions in bright green.**

### Supplementary Tables

**Table S1: Genome assembly statistics for all assembly pipelines tested**. The best performing pipeline per species is shown in bold. BUSCO scores correspond to the percentage of complete BUSCOs.

| Assembler | Features | *A. cristatus* | *C. marthae* | *C. pallidus* | *C. subcristatus* | *C. bakeri* | *C. carinata* | *I. delicatissima* |
|---|---|---|---|---|---|---|---|---|
| **Short-read assembly** | | | | | | | | |
| **SOAPdenovo** | **Genome size (Gbp)** | 3.6 | 2.32 | 2.28 | 2.49 | 2.22 | 2.25 | 2.20 |
| | **Scaffold N50 (kbp)** | 221 | 54 | 47 | 39 | 46 | 53 | 73 |
| | **Scaffold number** | 343,449 | 1,720,781 | 1,326,475 | 2,420,467 | 1,595,072 | 1,403,760 | 1,088,792 |
| | **BUSCO C (%)** | 76.5 | 69.5 | 73.8 | 72.5 | 71.0 | 74.0 | 75.3 |
| **MaSuRCA** | **Genome size (Gbp)** | 2.27 | 1.98 | 1.97 | 1.98 | 1.98 | **1.95** | 1.98 |
| | **Scaffold N50 (kbp)** | 445 | 95 | 70 | 93 | 64 | **83** | 71 |
| | **Scaffold number** | 28,166 | 44,521 | 56,278 | 46,323 | 66,871 | **48,324** | 53,352 |
| | **BUSCO C (%)** | 84.9 | 81.1 | 78.4 | 81.1 | 76.9 | **82.4** | 75.1 |
| **Sparse** | **Genome size (Gbp)** | 4.4 | 2.43 | 2.45 | 2.71 | 2.38 | 2.37 | 2.20 |
| | **Scaffold N50 (kbp)** | 1 | 2 | 4 | 1 | 3 | 3 | 5 |
| | **Scaffold number** | 3,400,000 | 7,892,079 | 8,424,092 | 11,722,634 | 7,168,493 | 7,232,523 | 5,401,287 |
| | **BUSCO C (%)** | 7.1 | 22.7 | 34.0 | 23.5 | 29.0 | 28.2 | 32.7 |
| **Hybrid assembly** | | | | | | | | |
| **dbg2olc** | **Genome size (Gbp)** | 1.97 | 2.04 | 2.02 | 2.03 | 2.16 | - | 2.00 |
| | **Scaffold N50 (kbp)** | 1,383 | 2,214 | 2,773 | 2,209 | 768 | - | 546 |
| | **Scaffold number** | 2,875 | 1824 | 1,528 | 1,795 | 5,382 | - | 6,978 |
| | **BUSCO C (%)** | 91.7 | 93.7 | 50.4 | 93.6 | 92.4 | - | 90.6 |
| **MaSuRCA** | **Genome size (Gbp)** | **1.98** | **2.02** | **2.01** | **1.99** | 2.06 | - | 1.99 |
| | **Scaffold N50 (kbp)** | **2,208** | **7,433** | **3,379** | **5,659** | 2,960 | - | 3,852 |
| | **Scaffold number** | **1,782** | **671** | **1,293** | **797** | 1,578 | - | 1,9101 |
| | **BUSCO C (%)** | **93.0** | **94.3** | **93.7** | **93.4** | 94.1 | - | 93.3 |
| **Wengan** | **Genome size (Gbp)** | 1.97 | 2.01 | 1.99 | 1.97 | **2.02** | - | **2.00** |
| | **Scaffold N50 (kbp)** | 1,431 | 5,468 | 1,112 | 1,657 | **8,875** | - | **4,405** |
| | **Scaffold number** | 3,129 | 1,085 | 3,737 | 2,874 | **1,522** | - | **1,522** |
| | **BUSCO C (%)** | 92.7 | 94.4 | 90.0 | 93.4 | **94.5** | - | **93.4** |

| | | | | | | | | |
|---|---|---|---|---|---|---|---|---|
| **Long-read assembly** | | | | | | | | |
| **Flye** | **Genome size (Gbp)** | 1.91 | 2.01 | 2.01 | 2.01 | 2.02 | - | 2.00 |
| | **Scaffold N50 (kbp)** | 106 | 1,971 | 915 | 5,987 | 6,805 | - | 3,583 |
| | **Scaffold number** | 36,858 | 2,451 | 5,626 | 1,303 | 2,157 | - | 2,678 |
| | **BUSCO C (%)** | 68.5 | 94.0 | 88.8 | 89.2 | 93.8 | - | 93.4 |
| **MECAT** | **Genome size (Gbp)** | 1.2 | 2.22 | 1.72 | 2.15 | 2.01 | - | 2.20 |
| | **Scaffold N50 (kbp)** | 10 | 415 | 67 | 511 | 460 | - | 331 |
| | **Scaffold number** | 26,798 | 10,633 | 33,084 | 8,703 | 11,231 | - | 13,710 |
| | **BUSCO C (%)** | 0.2 | 74.9 | 56.5 | 78.6 | 80.7 | - | 88.4 |
| **Miniasm** | **Genome size (Gbp)** | 0.27 | 2.03 | 1.79 | 2.05 | 2.04 | - | 2.04 |
| | **Scaffold N50 (kbp)** | 12 | 845 | 66 | 485 | 1,042 | - | 321 |
| | **Scaffold number** | 24,043 | 522 | 35,390 | 8,830 | 4,462 | - | 11,923 |
| | **BUSCO C (%)** | 0.1 | 3.9 | 10.1 | 4.3 | 37.2 | - | 30.4 |
| **Shasta** | **Genome size (Gbp)** | 0.2 | 1.99 | 2.4 | 1.89 | 2.08 | - | 1.10 |
| | **Scaffold N50 (kbp)** | 5 | 195 | 18 | 122 | 140 | - | 32 |
| | **Scaffold number** | 100 | 19,112 | 39,025 | 26,815 | 28,764 | - | 50,001 |
| | **BUSCO C (%)** | 0.0 | 61.6 | 5.4 | 51.6 | 75.7 | - | 24.4 |
| **wtdbg2** | **Genome size (Gbp)** | 1.89 | 1.87 | 1.97 | 1.95 | 1.90 | - | 1.91 |
| | **Scaffold N50 (kbp)** | 120 | 2,335 | 883 | 810 | 445 | - | 713 |
| | **Scaffold number** | 21,318 | 1831 | 4,566 | 3,395 | 9,618 | - | 9,495 |
| | **BUSCO C (%)** | 62.3 | 88.7 | 76.1 | 72.5 | 84.5 | - | 86.4 |

**Table S2: Sequencing summary statistics.** The tissue sampled is provided along with the platform, library types and instrument models. All sample, run and study accessions are detailed. The number of reads generated per run, and the total number of base pairs are detailed. Where possible we have also provided the N50 values.

| Species | Library type | Platform | Insert size (bp) | Instrument model | Sample | Run | Number of reads | Base pairs | Coverage | N50 (bp) |
|---|---|---|---|---|---|---|---|---|---|---|
| *Amblyrhynchus cristatus* | DNA-seq | Illumina | 500 | HiSeq 2000 | SAMN52407000 | SRR35754250 | 1,191,378,478 | 164,638,164,476 | 83.2 | NA |
| | DNA-seq | Illumina | 3,000 | HiSeq 2000 | | SRR35754253 | 232,478,948 | 22,934,718,569 | 11.6 | NA |
| | DNA-seq | Illumina | 8,000 | HiSeq 2000 | | SRR35754252 | 210,093,678 | 20,463,837,981 | 10.3 | NA |
| | DNA-seq | ONT | NA | MinION | | SRR35754251 | 9,515,006 | 24,955,769,248 | 12.6 | 3,706 |
| | RNA-seq | Illumina | 500 | NovaSeq 6000 | SAMN52407006 | SRR35754249 | 278,771,424 | 10,523,621,256 | NA | NA |
| *Conolophus marthae* | DNA-seq | Illumina | 500 | NovaSeq 6000 | SAMN52407002 | SRR35754262 | 821,432,964 | 124,036,377,564 | 61 | NA |
| | DNA-seq | ONT | NA | MinION | | SRR35754263 | 9,043,475 | 78,707,231,660 | 39 | 11,118 |
| *Conolophus pallidus* | DNA-seq | Illumina | 500 | NovaSeq 6000 | SAMN52407001 | SRR35754261 | 916,683,070 | 138,419,143,570 | 69 | NA |
| | DNA-seq | ONT | NA | MinION | | SRR35754248 | 4,258,083 | 26,879,582,859 | 13 | 8,032 |
| *Conolophus subcristatus* | DNA-seq | Illumina | 500 | NovaSeq 6000 | SAMN52407003 | SRR35754254 | 868,143,384 | 130,655,579,292 | 66 | NA |
| | DNA-seq | ONT | NA | MinION | | SRR35754255 | 5,485,962 | 45,510,110,239 | 23 | 10,317 |
| *Ctenosaura bakeri* | DNA-seq | Illumina | 500 | NovaSeq 6000 | SAMN52407004 | SRR35754259 | 66,137,090 | 133,209,153,424 | 66 | NA |
| | DNA-seq | ONT | NA | MinION | | SRR35754260 | 7,621,077 | 65,419,130,671 | 32 | 11,556 |
| *Cyclura carinata* | DNA-seq | Illumina | 500 | NovaSeq 6000 | SAMN52407005 | SRR35754258 | 67,801,930 | 135,964,564,882 | 70 | NA |
| *Iguana delicatissima* | DNA-seq | Illumina | 500 | NovaSeq 6000 | SAMN52407007 | SRR35754257 | 887,875,562 | 134,069,209,862 | 67 | NA |
| | DNA-seq | ONT | NA | MinION | | SRR35754256 | 16,041,008 | 71,404,239,969 | 36 | 6,354 |

**Table S3: Main repetitive element classes in total number and percentage of the genome.**

| | *A. cristatus* | | *C. marthae* | | *C. pallidus* | | *C. subcristatus* | | *C. bakeri* | | *C. carinata* | | *I. delicatissima* | |
|---|---|---|---|---|---|---|---|---|---|---|---|---|---|---|
| | **Number** | **%** | **Number** | **%** | **Number** | **%** | **Number** | **%** | **Number** | **%** | **Number** | **%** | **Number** | **%** |
| **Retroelements** | 1,350,316 | 20.39 | 1,307,860 | 20.04 | 1,240,992 | 20.64 | 1,340,014 | 20.89 | 1,461,000 | 20.53 | 1,313,797 | 18.77 | 1400,550 | 20.51 |
| **SINEs** | 423,817 | 3.37 | 338,356 | 2.49 | 392,491 | 3.18 | 436,470 | 3.29 | 423,727 | 3.23 | 383,721 | 3.06 | 491,347 | 3.67 |
| **LINEs** | 861,728 | 13.59 | 875,216 | 13.89 | 769,177 | 13.48 | 842,204 | 13.81 | 978,460 | 14.10 | 877,649 | 13.53 | 841,560 | 13.55 |
| **LTR elements** | 64,771 | 3.44 | 94,288 | 3.69 | 79,324 | 3.98 | 61,340 | 3.79 | 58,813 | 3.20 | 52,427 | 2.19 | 67,643 | 3.29 |
| **DNA transposons** | 258,875 | 2.93 | 271,348 | 3.17 | 280,003 | 3.24 | 214,670 | 2.91 | 272,797 | 2.85 | 210,780 | 2.01 | 225,204 | 2.32 |
| **Rolling-circles** | 1,033 | 0.02 | 20,665 | 0.17 | 3,696 | 0.02 | 1,226 | 0.01 | 1,722 | 0.02 | 939 | 0.01 | 986 | 0.01 |
| **Small RNA** | 27,082 | 0.15 | 21,184 | 0.09 | 35,880 | 0.18 | 25,310 | 0.18 | 26,702 | 0.12 | 35,602 | 0.15 | 32,953 | 0.19 |
| **Satellites** | 891 | 0.04 | 3,427 | 0.07 | 1,604 | 0.02 | 0 | 0.00 | 458 | 0.00 | 2,840 | 0.06 | 0 | 0.00 |
| **Simple repeats** | 490,745 | 0.91 | 467,668 | 0.87 | 474,383 | 0.91 | 465,496 | 0.85 | 483,094 | 1.02 | 463,344 | 0.92 | 454,640 | 0.87 |
| **Low complexity** | 56,479 | 0.15 | 55,994 | 0.16 | 55,338 | 0.14 | 54,343 | 0.14 | 55,270 | 0.14 | 53,025 | 0.14 | 53,637 | 0.13 |

**Table S4: Mean and 95% HPD interval of posterior divergence times in Mya under the node distributions for single-gene orthologs.**

| Node | Uniform distribution | | | Skewed-normal distribution | | | Cauchy distribution | | |
|---|---|---|---|---|---|---|---|---|---|
| | Posterior mean | 95% HPD | | Posterior mean | 95% HPD | | Posterior mean | 95% HPD | |
| 29 | 190.68 | 172.06 | 215.41 | 191.13 | 171.64 | 218.37 | 225.64 | 194.11 | 246.82 |
| 30 | 54.10 | 44.06 | 62.93 | 50.64 | 43.01 | 61.86 | 100.48 | 51.85 | 153.96 |
| 31 | 177.58 | 170.63 | 186.35 | 177.44 | 169.83 | 186.94 | 210.57 | 180.25 | 239.9 |
| 32 | 158.14 | 132.04 | 177.98 | 148.57 | 68.77 | 178.9 | 187.91 | 153.73 | 220 |
| 33 | 21.12 | 9.16 | 36.88 | 19.67 | 6.79 | 35.08 | 35.92 | 19.23 | 54.38 |
| 34 | 167.10 | 166.10 | 168.20 | 167.17 | 166.38 | 168.28 | 195.76 | 165.71 | 223.31 |
| 35 | 27.31 | 24.87 | 28.76 | 25.03 | 23.86 | 26.45 | 57.84 | 42.37 | 73.27 |
| 36 | 21.74 | 17.34 | 25.98 | 19.66 | 15.46 | 23.41 | 46.87 | 32.54 | 61.5 |
| 37 | 8.91 | 4.40 | 13.76 | 8.46 | 4.4 | 13.07 | 16.2 | 8.6 | 24.58 |
| 38 | 20.58 | 16.04 | 24.81 | 18.92 | 15.56 | 22.08 | 39.07 | 29.01 | 50.55 |
| 39 | 17.47 | 12.36 | 22.25 | 15.87 | 11.69 | 19.95 | 34.44 | 24.26 | 45.31 |
| 40 | 15.12 | 10.47 | 19.95 | 13.72 | 9.6 | 17.94 | 28.1 | 20.04 | 37.46 |
| 41 | 10.23 | 5.96 | 15.53 | 9.09 | 4.19 | 13.58 | 19.78 | 11.32 | 27.91 |
| 42 | 153.28 | 148.60 | 156.86 | 151.97 | 147.33 | 157.47 | 186.47 | 157.67 | 214.34 |
| 43 | 96.21 | 73.23 | 113.87 | 92.24 | 72.93 | 116.82 | 127.64 | 83.92 | 167.59 |
| 44 | 105.64 | 93.02 | 115.30 | 108.48 | 88.49 | 130.02 | 162.01 | 132.11 | 191.5 |
| 45 | 75.23 | 72.10 | 81.03 | 75.27 | 71.63 | 79.92 | 120.46 | 94.52 | 147.18 |
| 46 | 55.96 | 51.10 | 58.79 | 54.91 | 48.89 | 61.39 | 111.37 | 86.63 | 137.86 |
| 47 | 33.92 | 23.46 | 43.62 | 37.81 | 27.71 | 47.35 | 71.55 | 50.65 | 94.46 |
| 48 | 25.73 | 17.34 | 35.49 | 29.21 | 19.97 | 39.09 | 57.33 | 38.77 | 77.32 |
| 49 | 16.04 | 9.09 | 23.45 | 18.02 | 10.04 | 26.03 | 36.95 | 24.09 | 52.6 |
| 50 | 23.96 | 15.71 | 32.86 | 21.77 | 14.5 | 28.42 | 34.57 | 22.92 | 46.37 |
| 51 | 21.74 | 13.55 | 30.48 | 19.38 | 11.42 | 25.96 | 31.68 | 20.28 | 42.98 |
| 52 | 19.34 | 12.65 | 27.19 | 17.44 | 11.13 | 23.1 | 27.99 | 17.21 | 38.67 |
| 53 | 13.79 | 8.07 | 20.02 | 12.44 | 7.51 | 17.83 | 19.93 | 11.64 | 28.23 |
| 54 | 6.10 | 3.06 | 9.80 | 5.93 | 2.75 | 9.2 | 8.48 | 5.07 | 12.05 |
| 55 | 3.81 | 1.56 | 6.70 | 3.66 | 1.38 | 6.58 | 5.39 | 2.84 | 8.3 |

**Table S5: Mean and 95% HPD interval of posterior divergence times in Mya under skewed-normal distribution for the 95% UCE matrix.**

| | 4,153 UCEs | | | 500 UCEs | | | | | | 1,000 UCEs | | | | | |
|---|---|---|---|---|---|---|---|---|---|---|---|---|---|---|---|
| | No filter | | | Systematic bias filter | | | Default filter | | | Systematic bias filter | | | Default filter | | |
| | Mean | 95% HPD | | Mean | 95% HPD | | Mean | 95% HPD | | Mean | 95% HPD | | Mean | 95% HPD | |
| 29 | 196.09 | 176.23 | 219.54 | 193.43 | 173.39 | 228.52 | 193.75 | 175.31 | 228.55 | 195.77 | 174.13 | 231.28 | 190.74 | 173.03 | 219.31 |
| 30 | 52.8 | 43.36 | 70.41 | 51.53 | 43.86 | 63.74 | 51.17 | 43.86 | 62.61 | 51.58 | 43.83 | 63.84 | 52.59 | 44.15 | 67.41 |
| 31 | 175.02 | 169.16 | 181.99 | 176.63 | 169.8 | 188.07 | 178.1 | 171.43 | 190.18 | 177 | 170.22 | 188.36 | 177.28 | 170.87 | 187.86 |
| 32 | 154.56 | 110.59 | 172.43 | 152.96 | 94.99 | 172.79 | 157 | 121.93 | 177.28 | 160.7 | 142.21 | 171.77 | 152.51 | 91.87 | 171.85 |
| 33 | 20.14 | 14.4 | 31.77 | 17.39 | 8.27 | 26.34 | 17.38 | 9.61 | 24.9 | 18.42 | 8.67 | 31.07 | 17.72 | 8.97 | 33.06 |
| 34 | 167.13 | 166.45 | 168.43 | 167.11 | 166.45 | 168.38 | 167.13 | 166.46 | 168.41 | 167.09 | 166.44 | 168.34 | 167.11 | 166.45 | 168.35 |
| 35 | 25.55 | 24.1 | 27.95 | 24.92 | 23.92 | 26.47 | 25.08 | 23.95 | 26.83 | 25.16 | 23.98 | 26.89 | 25.27 | 24 | 27.17 |
| 36 | 20.99 | 16.75 | 25.66 | 19.58 | 13.77 | 23.31 | 19.22 | 13.34 | 23.49 | 20.35 | 16.41 | 23.56 | 19.91 | 15.25 | 24.59 |
| 37 | 4.8 | 2.41 | 9.31 | 6.44 | 3.03 | 13.1 | 4.78 | 2.19 | 8.52 | 6.08 | 2.77 | 11.84 | 4.9 | 1.8 | 9.73 |
| 38 | 20.73 | 18.9 | 25.72 | 17.25 | 12.1 | 20.92 | 17.8 | 13.89 | 23.67 | 18.67 | 15.7 | 22.66 | 17.98 | 14.25 | 24.82 |
| 39 | 15.91 | 8.75 | 23.69 | 13.66 | 7.57 | 18.38 | 13.92 | 8.86 | 20.66 | 15.41 | 10.52 | 20.44 | 14.29 | 9.72 | 19.69 |
| 40 | 12.47 | 9.69 | 15.9 | 11.58 | 6.74 | 15.98 | 11.54 | 7.44 | 16.29 | 12.15 | 8.18 | 17.07 | 11.65 | 9.04 | 15.23 |
| 41 | 9.11 | 7.64 | 11.43 | 7.02 | 3.37 | 11.03 | 6.92 | 3.53 | 11.89 | 7.33 | 4.03 | 12.01 | 7.28 | 4.78 | 11.05 |
| 42 | 153.06 | 147.98 | 160.02 | 155.92 | 148.68 | 164.78 | 154.3 | 148.36 | 161.62 | 155.68 | 148.68 | 163.92 | 154.62 | 148.44 | 162.18 |
| 43 | 90.18 | 73.2 | 108.94 | 88.8 | 74.62 | 111.58 | 88.27 | 73.97 | 114.16 | 87.43 | 73.35 | 114.15 | 88.87 | 75.27 | 107.75 |
| 44 | 105.02 | 94.66 | 124.37 | 111.96 | 93.29 | 134.89 | 112.53 | 97.92 | 133.82 | 110.27 | 90.02 | 137.25 | 113.05 | 91.16 | 132.87 |
| 45 | 75.01 | 72.16 | 80.3 | 75.39 | 72.09 | 81.11 | 75.07 | 72.01 | 80.25 | 75.01 | 72.05 | 80.21 | 75.07 | 72.02 | 80.29 |
| 46 | 54.96 | 49.03 | 63.09 | 53.39 | 47.99 | 60.71 | 53.81 | 48.25 | 59.84 | 54.6 | 49.07 | 61.34 | 54.04 | 48.25 | 60.25 |
| 47 | 36.09 | 27.95 | 49.05 | 30.63 | 20.64 | 44.43 | 31.44 | 21.66 | 42.04 | 35.31 | 23.67 | 42.59 | 29.08 | 17.53 | 43.19 |
| 48 | 22.33 | 16.66 | 28.29 | 22.21 | 14.14 | 35.4 | 22.86 | 15.34 | 32.52 | 25.5 | 16.77 | 35.3 | 21.12 | 12.37 | 30.19 |
| 49 | 10.74 | 5.89 | 17.11 | 13.25 | 6.62 | 22.17 | 12.93 | 6.94 | 20.46 | 15.63 | 9.71 | 22.77 | 12.64 | 6.77 | 21.95 |
| 50 | 24.33 | 21.02 | 28.89 | 19.82 | 13.61 | 24.95 | 18.75 | 12.54 | 34.79 | 21.16 | 15.07 | 31.98 | 17.08 | 11.33 | 28.61 |
| 51 | 22.99 | 19.8 | 27.23 | 18.62 | 12.43 | 23.71 | 16.92 | 10.95 | 31.73 | 19.64 | 14.03 | 30.41 | 15.56 | 10.14 | 26.71 |
| 52 | 19.23 | 14.16 | 22.83 | 13.53 | 8.47 | 19.07 | 12.02 | 7.02 | 24.76 | 15.5 | 10.55 | 27.12 | 11.78 | 7.05 | 25.41 |
| 53 | 15.28 | 10.64 | 20.64 | 8.99 | 4.8 | 14.67 | 8.05 | 4.34 | 19.79 | 10.18 | 5.64 | 22.06 | 7.53 | 4.06 | 20.21 |
| 54 | 2.44 | 1.37 | 4.66 | 3.07 | 1.55 | 5.66 | 2.38 | 1.22 | 4.32 | 3.1 | 1.46 | 6.51 | 2.48 | 1.27 | 5.33 |
| 55 | 0.85 | 0.36 | 1.67 | 1.04 | 0.41 | 2.31 | 0.65 | 0.3 | 1.29 | 1 | 0.39 | 2.26 | 0.71 | 0.31 | 1.52 |

**Table S6: Mean and 95% HPD interval of posterior divergence times in Mya under skewed-normal distribution for the 75% UCE matrix.**

| | 4,663 UCEs | | | 500 UCEs | | | | | | 1,000 UCEs | | | | | |
|---|---|---|---|---|---|---|---|---|---|---|---|---|---|---|---|
| | No filter | | | Systematic bias filter | | | Default filter | | | Systematic bias filter | | | Default filter | | |
| | Mean | 95% HPD | | Mean | 95% HPD | | Mean | 95% HPD | | Mean | 95% HPD | | Mean | 95% HPD | |
| 29 | 188.47 | 175.56 | 209.89 | 192.93 | 174.79 | 227.01 | 200.23 | 176.22 | 249.95 | 194.13 | 174.73 | 227.21 | 188.42 | 174.93 | 210.43 |
| 30 | 51.82 | 43.79 | 64.37 | 52.06 | 44.07 | 66.65 | 51.37 | 43.88 | 63.3 | 51.49 | 43.75 | 66.85 | 52.65 | 43.96 | 69.57 |
| 31 | 176.63 | 170.72 | 183.71 | 176.42 | 170.21 | 184.54 | 179.43 | 171.38 | 192.24 | 177.77 | 170.4 | 193.85 | 177.68 | 170.83 | 189.7 |
| 32 | 158.37 | 141.86 | 168.34 | 159.03 | 123.33 | 172.84 | 155.26 | 105.16 | 174.24 | 155.23 | 113.13 | 173.39 | 155.68 | 111.62 | 170.72 |
| 33 | 26.99 | 12.92 | 50.71 | 17.05 | 7.97 | 24.99 | 19.21 | 8.05 | 48.26 | 18.34 | 7.77 | 33.75 | 18.65 | 8.52 | 38.03 |
| 34 | 167.14 | 166.46 | 168.43 | 167.09 | 166.45 | 168.33 | 167.13 | 166.45 | 168.41 | 167.1 | 166.45 | 168.35 | 167.13 | 166.46 | 168.44 |
| 35 | 25.42 | 24.17 | 27.16 | 25.04 | 23.97 | 26.59 | 25.04 | 23.93 | 26.73 | 25.08 | 23.93 | 26.86 | 24.85 | 23.9 | 26.28 |
| 36 | 20.03 | 15.12 | 24.69 | 20.08 | 14.26 | 23.65 | 19.4 | 15.17 | 22.82 | 19.13 | 14.12 | 23.55 | 19.36 | 15.71 | 23.92 |
| 37 | 4.84 | 2.75 | 7.69 | 6.55 | 2.89 | 12.94 | 5.07 | 2.29 | 9.49 | 5.95 | 3.08 | 10.23 | 4.8 | 2.52 | 8.1 |
| 38 | 17.89 | 13.27 | 24.62 | 17.83 | 13.59 | 21.59 | 17.54 | 14.61 | 22.74 | 18.1 | 14.16 | 23.08 | 18.62 | 14.65 | 22.77 |
| 39 | 14.28 | 9.61 | 21.57 | 14.5 | 8.63 | 19.57 | 13.57 | 9.09 | 18.1 | 14.98 | 10.3 | 20.58 | 14.72 | 9.1 | 20.27 |
| 40 | 11.4 | 9.38 | 13.36 | 11.8 | 7.57 | 16.35 | 11.38 | 7.59 | 15.35 | 12.17 | 8.13 | 16.47 | 11.09 | 7.17 | 15.34 |
| 41 | 8.21 | 7.18 | 9.7 | 6.86 | 3.52 | 11.42 | 6.86 | 3.37 | 10.84 | 7.46 | 3.8 | 12.98 | 6.38 | 3.14 | 10.25 |
| 42 | 153.14 | 148.24 | 160.25 | 156.46 | 148.68 | 165.29 | 153.84 | 148.31 | 160.8 | 155.26 | 148.61 | 164 | 154.4 | 148.37 | 162.03 |
| 43 | 87.05 | 72.73 | 107.88 | 88.42 | 73.99 | 112.25 | 89.09 | 73.94 | 111.9 | 89.53 | 75.01 | 110.46 | 90.19 | 74.1 | 119.11 |
| 44 | 108.43 | 93.53 | 127.7 | 112.45 | 93.3 | 136.13 | 108.15 | 89.56 | 128.96 | 115.25 | 96.51 | 134.28 | 118.31 | 102.41 | 142.53 |
| 45 | 74.74 | 72.01 | 79.07 | 75.29 | 72.11 | 80.96 | 74.98 | 72.03 | 79.74 | 75.3 | 72.09 | 81.38 | 75.18 | 72.04 | 80.92 |
| 46 | 52.45 | 47.87 | 62.12 | 53.48 | 47.61 | 60.72 | 54.73 | 48.18 | 62.11 | 53.29 | 48.41 | 60.12 | 54.35 | 48.43 | 60.36 |
| 47 | 36.85 | 29.92 | 45.89 | 26.75 | 19.66 | 37.22 | 28.36 | 19.17 | 40.19 | 28.44 | 20.36 | 39.1 | 31.55 | 21.72 | 41.92 |
| 48 | 22.9 | 18.1 | 26.68 | 20.2 | 14.24 | 27.5 | 21.36 | 12.72 | 32.78 | 20.72 | 14.04 | 31.77 | 21.63 | 16.66 | 25.5 |
| 49 | 14.2 | 7.66 | 19.54 | 12.18 | 6.29 | 19.86 | 12.81 | 6.76 | 20.43 | 13.1 | 6.89 | 23.08 | 12.88 | 7.1 | 19.36 |
| 50 | 22.98 | 19.37 | 27.18 | 19.61 | 14.06 | 25.52 | 20.31 | 14.73 | 25.83 | 22.45 | 14.79 | 32.32 | 18.9 | 12.85 | 29.54 |
| 51 | 21.59 | 17.42 | 25.07 | 18.46 | 13.08 | 24.46 | 18.47 | 13.18 | 23.88 | 20.94 | 13.55 | 30.11 | 17.14 | 11.65 | 25.78 |
| 52 | 16.13 | 12.81 | 21.75 | 13.71 | 9.17 | 19.27 | 13.88 | 8.24 | 20.53 | 16.06 | 9.36 | 26.84 | 12.83 | 7.42 | 23.45 |
| 53 | 10.96 | 7.99 | 18.95 | 9.15 | 5.04 | 14.61 | 9.33 | 5.11 | 15.74 | 10.06 | 5.07 | 18.35 | 8.03 | 4 | 16.31 |
| 54 | 3.16 | 1.27 | 8.9 | 2.96 | 1.5 | 5.62 | 2.63 | 1.3 | 4.89 | 2.97 | 1.24 | 5.78 | 2.53 | 1.17 | 4.57 |
| 55 | 0.93 | 0.36 | 2.22 | 0.98 | 0.4 | 1.96 | 0.68 | 0.3 | 1.4 | 0.98 | 0.36 | 2.05 | 0.7 | 0.3 | 1.52 |

**Table S7: Mean and 95% HPD interval of posterior divergence times in Mya under uniform distribution for the 95% UCE matrix.**

| | 4,153 UCEs | | | 500 UCEs | | | | | | 1,000 UCEs | | | | | |
|---|---|---|---|---|---|---|---|---|---|---|---|---|---|---|---|
| | No filter | | | Systematic bias filter | | | Default filter | | | Systematic bias filter | | | Default filter | | |
| | Mean | 95% HPD | | Mean | 95% HPD | | Mean | 95% HPD | | Mean | 95% HPD | | Mean | 95% HPD | |
| 29 | 205.12 | 183.12 | 223.13 | 201.61 | 176.15 | 238.39 | 197.69 | 177.51 | 233.12 | 200.69 | 174.82 | 238.7 | 203.59 | 179.25 | 235.15 |
| 30 | 55.25 | 44.57 | 63.43 | 55.08 | 44.8 | 63.47 | 55.03 | 44.7 | 63.38 | 52.69 | 44.38 | 63.18 | 55.91 | 44.83 | 63.64 |
| 31 | 178.49 | 171.65 | 186.47 | 178.24 | 170.65 | 187.78 | 180.02 | 172.17 | 188.86 | 177.97 | 170.42 | 187.34 | 179.99 | 171.52 | 188.08 |
| 32 | 160.22 | 129.46 | 173.35 | 160.41 | 132.02 | 176.54 | 151.61 | 81.9 | 174.13 | 160.91 | 133.97 | 174.12 | 107.02 | 36.93 | 172.81 |
| 33 | 19.09 | 9.83 | 26.98 | 17.69 | 8.37 | 27.66 | 17.35 | 8.81 | 26.48 | 17.54 | 8.85 | 27.21 | 15.26 | 7.61 | 26.15 |
| 34 | 167.08 | 166.14 | 168.27 | 167.01 | 166.13 | 168.25 | 167.04 | 166.14 | 168.26 | 167.01 | 166.13 | 168.25 | 167.03 | 166.13 | 168.26 |
| 35 | 26.2 | 23.29 | 28.55 | 26.61 | 23.42 | 28.52 | 27.16 | 23.91 | 28.58 | 26.41 | 23.36 | 28.47 | 27.03 | 24.68 | 28.52 |
| 36 | 20.81 | 18.42 | 24.19 | 21.02 | 14.41 | 25.76 | 21.41 | 16.57 | 25.21 | 21.39 | 17.51 | 25.25 | 21.15 | 14.84 | 25.09 |
| 37 | 5.59 | 0.77 | 9.93 | 6.63 | 2.97 | 12.1 | 5.33 | 2.63 | 9.66 | 6.14 | 2.99 | 10.99 | 5.46 | 2.44 | 9.21 |
| 38 | 20.15 | 17.88 | 23.67 | 18.41 | 13.77 | 23.17 | 18.75 | 15.02 | 23.27 | 17.86 | 14.56 | 21.79 | 19.4 | 15.63 | 24.05 |
| 39 | 17.03 | 10.52 | 21.15 | 15.02 | 10.2 | 20.87 | 14.67 | 11.43 | 19.85 | 14.47 | 10.04 | 19.04 | 15.13 | 9.9 | 21.86 |
| 40 | 11.25 | 9 | 13.91 | 12.25 | 7.25 | 17.31 | 12.15 | 8.59 | 17.17 | 11.03 | 8.43 | 14.07 | 12.65 | 8.62 | 17.24 |
| 41 | 4.74 | 3.2 | 8.53 | 7.37 | 3.44 | 12.34 | 7.48 | 4.23 | 13.15 | 7.01 | 4.61 | 10.84 | 8.08 | 5.04 | 11.58 |
| 42 | 153.96 | 148.71 | 156.84 | 154.49 | 149.2 | 156.91 | 154.22 | 148.95 | 156.86 | 154.62 | 149.25 | 156.93 | 154.16 | 148.84 | 156.87 |
| 43 | 94.35 | 71.65 | 113.53 | 91.31 | 72.18 | 112.31 | 86.17 | 71.69 | 110.74 | 89.56 | 71.72 | 112.5 | 92.96 | 72.02 | 113 |
| 44 | 107.75 | 100.29 | 113.94 | 107.25 | 93.33 | 114.87 | 107.84 | 97.48 | 114.89 | 107.25 | 94.74 | 114.49 | 108.15 | 96.22 | 114.85 |
| 45 | 75.26 | 72.19 | 82.76 | 75.59 | 72.2 | 82.51 | 74.95 | 72.18 | 81.43 | 75.69 | 72.2 | 82.51 | 75.99 | 72.21 | 83.17 |
| 46 | 55.24 | 50 | 58.43 | 55.13 | 48.86 | 58.31 | 55.43 | 49.38 | 58.35 | 56.23 | 51.98 | 58.4 | 55.61 | 49.63 | 58.34 |
| 47 | 32.29 | 20.9 | 48.59 | 28.21 | 18.53 | 41.98 | 27.74 | 20.43 | 41.65 | 38.11 | 27.82 | 48.76 | 31.24 | 23.05 | 47.46 |
| 48 | 22.57 | 16.11 | 28.99 | 21.67 | 13.1 | 35.72 | 20.04 | 13.96 | 27.03 | 26.61 | 17.72 | 36.29 | 23.57 | 17.59 | 37.15 |
| 49 | 11.76 | 7.71 | 16.9 | 13.04 | 6.23 | 22.73 | 11.54 | 6 | 18.6 | 14.8 | 8.32 | 26.02 | 12.43 | 7.39 | 18.31 |
| 50 | 29.4 | 20.12 | 37.52 | 20.29 | 16.27 | 30.15 | 19.93 | 14.8 | 26.2 | 21.57 | 15.34 | 34.53 | 21.86 | 14.64 | 31.44 |
| 51 | 27.71 | 19.21 | 35.54 | 19.12 | 15.14 | 29.32 | 18.09 | 13.16 | 23.74 | 20.05 | 13.92 | 33.35 | 19.91 | 12.53 | 28.65 |
| 52 | 23.48 | 16.18 | 31.89 | 14.63 | 9.1 | 22.58 | 12.34 | 7.91 | 18.13 | 14.59 | 8.3 | 29.6 | 14.09 | 8.03 | 21.34 |
| 53 | 19.31 | 11.5 | 28.97 | 9.98 | 5.48 | 16.78 | 7.71 | 4.43 | 15.75 | 10.39 | 5.77 | 23.28 | 9.19 | 4.93 | 16.53 |
| 54 | 2.64 | 1.13 | 6.09 | 3.34 | 1.66 | 6.31 | 2.51 | 1.27 | 4.84 | 2.95 | 1.46 | 5.9 | 2.75 | 1.22 | 5.89 |
| 55 | 0.95 | 0.35 | 2.99 | 1.11 | 0.44 | 2.25 | 0.66 | 0.3 | 1.31 | 0.91 | 0.37 | 1.71 | 0.77 | 0.31 | 1.64 |

**Table S8: Mean and 95% HPD interval of posterior divergence times in Mya under uniform distribution for the 75% UCE matrix.**

| | 4,663 UCEs | | | 500 UCEs | | | | | | 1,000 UCEs | | | | | |
|---|---|---|---|---|---|---|---|---|---|---|---|---|---|---|---|
| | No filter | | | Systematic bias filter | | | Default filter | | | Systematic bias filter | | | Default filter | | |
| | Mean | 95% HPD | | Mean | 95% HPD | | Mean | 95% HPD | | Mean | 95% HPD | | Mean | 95% HPD | |
| 29 | 190.34 | 174.23 | 222.26 | 201.4 | 177.49 | 237.36 | 205 | 179.76 | 239.9 | 199.33 | 175.3 | 236.83 | 208.33 | 176.26 | 240.96 |
| 30 | 54.75 | 44.71 | 63.51 | 54.04 | 44.49 | 63.27 | 55.41 | 44.92 | 63.46 | 53.95 | 44.45 | 63.49 | 53.47 | 44.34 | 63.33 |
| 31 | 179.82 | 170.96 | 188.54 | 178.53 | 170.73 | 187.82 | 180.25 | 171.79 | 188.88 | 177.62 | 169.79 | 187.05 | 179.97 | 172.01 | 188.04 |
| 32 | 155.61 | 129.13 | 175.64 | 147.21 | 86.51 | 174.5 | 155.89 | 110.96 | 172.19 | 158.07 | 105.88 | 174.36 | 159.88 | 131.77 | 173.11 |
| 33 | 19.36 | 10.31 | 29.61 | 19.8 | 6.32 | 53.79 | 16.38 | 8.21 | 25.79 | 19.69 | 11.07 | 33.26 | 16.64 | 8.67 | 25.7 |
| 34 | 167.02 | 166.13 | 168.24 | 167.02 | 166.13 | 168.25 | 167.04 | 166.13 | 168.27 | 167.02 | 166.13 | 168.25 | 167.04 | 166.13 | 168.26 |
| 35 | 27.59 | 25.76 | 28.66 | 26.9 | 23.81 | 28.55 | 26.15 | 23.18 | 28.47 | 26.51 | 23.61 | 28.5 | 27.26 | 24.71 | 28.63 |
| 36 | 22.46 | 19.43 | 25.76 | 21.3 | 15.36 | 26.13 | 20.03 | 14.9 | 25.16 | 20.84 | 15.09 | 25.1 | 21.31 | 15.07 | 25.75 |
| 37 | 6.1 | 2.75 | 9.31 | 6.42 | 2.96 | 11.45 | 4.72 | 2.22 | 7.97 | 5.89 | 2.64 | 10.61 | 5.39 | 2.02 | 9.8 |
| 38 | 21.05 | 16.48 | 25.93 | 18.66 | 13.97 | 22.67 | 17.16 | 12.25 | 21.81 | 18.6 | 13.86 | 22.76 | 18.48 | 13.62 | 24.91 |
| 39 | 17.32 | 11.79 | 22.03 | 14.97 | 10.02 | 19.31 | 12.72 | 6.87 | 18.29 | 15.22 | 10.1 | 19.61 | 13.88 | 8.62 | 18.4 |
| 40 | 12.99 | 9.7 | 17.21 | 12.24 | 8.2 | 17.22 | 10.86 | 7.44 | 15.08 | 12.58 | 8.9 | 17.1 | 12.1 | 8.2 | 19.93 |
| 41 | 7.78 | 5.5 | 11.1 | 7.37 | 3.96 | 11.94 | 6.33 | 3.58 | 9.82 | 7.33 | 4.22 | 11.89 | 7.76 | 3.95 | 15.28 |
| 42 | 154.32 | 149.22 | 156.85 | 154.41 | 149.01 | 156.9 | 154.19 | 148.85 | 156.87 | 154.56 | 149.22 | 156.94 | 154.07 | 148.78 | 156.87 |
| 43 | 88.87 | 71.74 | 111.85 | 92.03 | 71.96 | 112.67 | 91.21 | 72.11 | 112.44 | 88.26 | 71.55 | 111.67 | 90.2 | 71.8 | 111.26 |
| 44 | 106.51 | 92.91 | 115.05 | 107.26 | 90.39 | 114.89 | 107.97 | 96.39 | 114.91 | 106.98 | 94.48 | 114.74 | 108.14 | 97.35 | 115.24 |
| 45 | 74.62 | 72.18 | 79.07 | 76.01 | 72.21 | 82.89 | 75.22 | 72.18 | 82.33 | 75.2 | 72.2 | 81.52 | 75.05 | 72.19 | 81.65 |
| 46 | 55.86 | 50.78 | 58.54 | 55.35 | 49.6 | 58.3 | 55.8 | 49.65 | 58.37 | 55.86 | 50.8 | 58.35 | 55.56 | 50.25 | 58.31 |
| 47 | 41.19 | 31.65 | 48.36 | 29.79 | 21.11 | 41.36 | 28.54 | 20.43 | 43.65 | 31.24 | 18.86 | 45.01 | 32.11 | 21.91 | 40.9 |
| 48 | 24.82 | 20.06 | 32.64 | 22.59 | 15.47 | 33.53 | 20.49 | 14.08 | 27.92 | 22.4 | 14.19 | 31.17 | 23.99 | 15.59 | 36.01 |
| 49 | 13.66 | 7.13 | 18.46 | 14.16 | 7.82 | 22.99 | 11.91 | 7.21 | 16.71 | 12.83 | 6.31 | 20.42 | 12.06 | 6.26 | 17.98 |
| 50 | 25.75 | 20.74 | 29.04 | 26.14 | 17.02 | 41.59 | 18.1 | 12.05 | 33.12 | 22.55 | 12.69 | 37.56 | 25.95 | 16.04 | 40.85 |
| 51 | 24.23 | 19.91 | 27.58 | 24.79 | 16 | 40.19 | 16.13 | 10.3 | 30 | 21.19 | 11.77 | 36.14 | 24.2 | 14.39 | 38.36 |
| 52 | 18.32 | 14.61 | 23.87 | 16.34 | 9.48 | 35.33 | 11.03 | 6.44 | 19.68 | 16.12 | 9.05 | 33.54 | 18.55 | 10.72 | 32.52 |
| 53 | 12.76 | 6.5 | 20.14 | 11.11 | 5.76 | 24.62 | 6.84 | 3.51 | 12.61 | 11.28 | 6.15 | 30.34 | 13.38 | 4.83 | 26.54 |
| 54 | 3.18 | 1.55 | 5.95 | 3.3 | 1.45 | 5.91 | 2.35 | 1.22 | 4.29 | 3.23 | 1.64 | 5.91 | 2.85 | 1.17 | 6.15 |
| 55 | 1.17 | 0.42 | 2.55 | 1.09 | 0.42 | 2.31 | 0.65 | 0.29 | 1.32 | 0.97 | 0.37 | 2.01 | 0.77 | 0.32 | 1.85 |

**Table S9: Mean and 95% HPD interval of posterior divergence times in Mya under Cauchy distribution for the 95% UCE matrix.**

| | 4,153 UCEs | | | 500 UCEs | | | | | | 1,000 UCEs | | | | | |
|---|---|---|---|---|---|---|---|---|---|---|---|---|---|---|---|
| | No filter | | | Systematic bias filter | | | Default filter | | | Systematic bias filter | | | Default filter | | |
| | Mean | 95% HPD | | Mean | 95% HPD | | Mean | 95% HPD | | Mean | 95% HPD | | Mean | 95% HPD | |
| 29 | 228.1 | 195.26 | 246.49 | 226.37 | 192.76 | 244.95 | 229.56 | 198.43 | 245.81 | 226.3 | 191.34 | 245.15 | 226.18 | 191.01 | 245.17 |
| 30 | 108.21 | 68.23 | 144.99 | 102.66 | 51.98 | 156.18 | 119.18 | 78.13 | 165.52 | 77.22 | 45.55 | 143.83 | 103.48 | 70.51 | 147.02 |
| 31 | 215.74 | 182.72 | 241.85 | 207.98 | 177.49 | 237.85 | 208.27 | 179.6 | 237.79 | 211.25 | 178.97 | 238.38 | 213.95 | 180.31 | 239.63 |
| 32 | 198.54 | 166.41 | 230.04 | 186.79 | 143.83 | 220 | 181.99 | 150.55 | 213.64 | 178.44 | 136.03 | 215 | 174.68 | 116.61 | 219.23 |
| 33 | 29.87 | 17.8 | 53.56 | 28.27 | 14.57 | 49.8 | 25.31 | 14.2 | 41.66 | 28.25 | 11.38 | 57.55 | 30.08 | 10.37 | 53.94 |
| 34 | 197.75 | 167.4 | 229.98 | 193.25 | 165.82 | 222.91 | 190.07 | 164.57 | 220.93 | 197.61 | 166.69 | 226.47 | 198.34 | 167.3 | 225.76 |
| 35 | 86.51 | 69.26 | 111.83 | 57.02 | 40.28 | 83.77 | 54.21 | 38.72 | 102.31 | 62.14 | 33.31 | 102.24 | 51.44 | 34.27 | 92.66 |
| 36 | 70.49 | 47.81 | 95.62 | 46.08 | 29.65 | 64.74 | 41.4 | 27.64 | 61.74 | 47.98 | 25.82 | 80.9 | 39.65 | 25.17 | 68.66 |
| 37 | 13.81 | 5.88 | 41.3 | 9.93 | 5.04 | 18.07 | 8 | 3.93 | 16.83 | 12.92 | 4.91 | 32.72 | 11.07 | 4.09 | 36.71 |
| 38 | 62.18 | 49.13 | 78.3 | 34.96 | 24.55 | 57.31 | 32.78 | 23.96 | 46.49 | 40.84 | 22.65 | 62.44 | 31.37 | 20.7 | 44.75 |
| 39 | 41.03 | 9.71 | 68.65 | 30.06 | 20.56 | 48.56 | 25.71 | 14.89 | 35.1 | 32.02 | 16.57 | 52.05 | 22.51 | 11.6 | 30.94 |
| 40 | 53.85 | 42.77 | 66.24 | 22.33 | 13.98 | 37.27 | 20.42 | 12.34 | 35.24 | 27.12 | 14.3 | 45.97 | 19.64 | 11.49 | 35.04 |
| 41 | 15.54 | 5.29 | 37.75 | 13.46 | 7.15 | 23.5 | 11.47 | 5.83 | 23.17 | 17.12 | 6.8 | 31.94 | 10.44 | 4.49 | 20.61 |
| 42 | 191.44 | 160.6 | 225.02 | 190.52 | 162.92 | 220.2 | 184.45 | 158.16 | 215.15 | 193.95 | 163.33 | 222.64 | 192.33 | 161.66 | 220.16 |
| 43 | 114.06 | 68.28 | 174.8 | 116.47 | 74.31 | 168.49 | 93.19 | 70.7 | 135.34 | 112.94 | 71.4 | 159.79 | 103.36 | 71.8 | 141.67 |
| 44 | 172.4 | 137.53 | 219.41 | 165.21 | 137.4 | 196.22 | 153.41 | 115.03 | 196.41 | 177.12 | 148.21 | 206.21 | 167.58 | 133.04 | 198.94 |
| 45 | 137.98 | 113.89 | 161.22 | 119.9 | 91.84 | 165.07 | 104.15 | 77.03 | 151.33 | 154.9 | 121.93 | 188.69 | 132.7 | 98.57 | 164.01 |
| 46 | 131.38 | 107.33 | 156.84 | 110.87 | 83.49 | 157.47 | 94.62 | 68.42 | 141.22 | 146.89 | 115.42 | 182.04 | 124.1 | 93.16 | 156.53 |
| 47 | 122.98 | 100.54 | 148.9 | 68.88 | 44.19 | 139.91 | 65.62 | 42.08 | 124.81 | 124.98 | 89.22 | 163.69 | 113.6 | 82.07 | 150.4 |
| 48 | 118.92 | 96.41 | 147.85 | 55 | 33.39 | 120.51 | 53.18 | 32.12 | 112.23 | 112.71 | 80.11 | 151.52 | 104.6 | 72.5 | 146.44 |
| 49 | 53.99 | 20.23 | 85.17 | 30.49 | 17.97 | 53.29 | 26.84 | 15.85 | 43.51 | 46.05 | 23.37 | 77.6 | 37.05 | 18.11 | 54.63 |
| 50 | 66.16 | 37.16 | 97.09 | 30.37 | 18.95 | 73.29 | 26.97 | 18.89 | 36.65 | 44.71 | 15.86 | 80.14 | 34.78 | 21.97 | 53.86 |
| 51 | 62.78 | 34.6 | 92.31 | 28.89 | 17.86 | 69.84 | 24.68 | 17.1 | 33.81 | 41.66 | 14.39 | 76.4 | 31.58 | 19.55 | 48.43 |
| 52 | 45.92 | 22.15 | 80.79 | 21.49 | 13.14 | 49.48 | 17.17 | 11.11 | 25.16 | 28.94 | 9.51 | 66.91 | 22.04 | 12.73 | 36.37 |
| 53 | 27.38 | 11.83 | 65.6 | 14.93 | 8.75 | 39.12 | 11.18 | 6.9 | 18.06 | 19.26 | 6.21 | 52.3 | 13.94 | 7.1 | 25.23 |
| 54 | 5.77 | 2.6 | 10.33 | 4.42 | 2.48 | 8.7 | 3.37 | 1.92 | 6.04 | 5.34 | 2.15 | 12.84 | 3.87 | 1.93 | 8.1 |
| 55 | 2.11 | 0.66 | 5.46 | 1.48 | 0.73 | 3.01 | 0.88 | 0.44 | 1.71 | 1.57 | 0.53 | 3.76 | 1.14 | 0.46 | 2.45 |

**Table S10: Mean and 95% HPD interval of posterior divergence times in Mya under Cauchy distribution for the 75% UCE matrix.**

| | 4,663 UCEs | | | 500 UCEs | | | | | | 1,000 UCEs | | | | | |
|---|---|---|---|---|---|---|---|---|---|---|---|---|---|---|---|
| | No filter | | | Systematic bias filter | | | Default filter | | | Systematic bias filter | | | Default filter | | |
| | Mean | 95% HPD | | Mean | 95% HPD | | Mean | 95% HPD | | Mean | 95% HPD | | Mean | 95% HPD | |
| 29 | 226.95 | 192.11 | 246.49 | 225.91 | 190.68 | 244.8 | 227.43 | 194.69 | 245.12 | 226.61 | 192.73 | 245.07 | 228.42 | 196.11 | 245.5 |
| 30 | 92.88 | 58.83 | 145.76 | 124.97 | 84.44 | 167.04 | 112.76 | 62.7 | 205.05 | 100.85 | 59.56 | 149.22 | 73.81 | 45.2 | 125.47 |
| 31 | 215.21 | 181.79 | 239.67 | 210.64 | 177.5 | 239.5 | 210.33 | 179.81 | 238.64 | 211.66 | 179.47 | 239.06 | 211.66 | 181.32 | 238.3 |
| 32 | 190.84 | 157.66 | 217.46 | 186.7 | 145.52 | 219.14 | 173.17 | 124.16 | 211.02 | 185.71 | 137.29 | 220.54 | 176.45 | 123.89 | 216.74 |
| 33 | 43.1 | 26.55 | 73.77 | 29.45 | 15.26 | 51.24 | 24.54 | 12.49 | 43.94 | 30.14 | 15.45 | 53.29 | 32.75 | 11.41 | 68.44 |
| 34 | 200.54 | 168.86 | 226.62 | 196.96 | 166.55 | 226.91 | 192.99 | 165.87 | 222.51 | 196.41 | 166.78 | 226.4 | 190.79 | 165.57 | 220.75 |
| 35 | 75.34 | 54.32 | 105.72 | 59.86 | 38.89 | 102.58 | 55.99 | 32.04 | 119.04 | 52.79 | 28.23 | 102.65 | 81.06 | 47.28 | 111.88 |
| 36 | 64.8 | 45.92 | 93.78 | 49.69 | 30.6 | 89.72 | 42.52 | 22.59 | 71.47 | 43.48 | 21.88 | 90.16 | 64.61 | 38.29 | 91.88 |
| 37 | 11.13 | 4.22 | 28.37 | 10.45 | 5.6 | 17.77 | 8.02 | 3.57 | 16.77 | 11.99 | 4.19 | 37.79 | 13.21 | 5.22 | 36.87 |
| 38 | 40.55 | 28.74 | 52.5 | 36.12 | 23.67 | 52.14 | 30.57 | 19.74 | 49.78 | 29.94 | 17.93 | 52.79 | 51.02 | 28.47 | 70.03 |
| 39 | 27.8 | 12.15 | 41.65 | 30.66 | 18.64 | 46.06 | 21.34 | 8.7 | 32.48 | 23.91 | 12.17 | 43.15 | 39.28 | 14.89 | 61.04 |
| 40 | 25.29 | 16.83 | 40.06 | 22.29 | 13.92 | 34.29 | 18.93 | 11.18 | 39.36 | 20.07 | 10.02 | 41.05 | 32.85 | 18.71 | 47.94 |
| 41 | 16.56 | 10.37 | 32.98 | 13.37 | 6.64 | 22.81 | 10.24 | 5.22 | 19.07 | 11.27 | 4.89 | 25.44 | 17.02 | 6.86 | 32.06 |
| 42 | 195.16 | 164.31 | 220.37 | 194.41 | 164.32 | 224.22 | 186.73 | 159.09 | 215.44 | 192.74 | 163.59 | 222.36 | 183.07 | 157.6 | 213.7 |
| 43 | 96.11 | 68.03 | 124.06 | 103.13 | 71.47 | 154.47 | 98.57 | 72.2 | 133.62 | 101.91 | 71.07 | 148.06 | 101.49 | 72.16 | 137.78 |
| 44 | 175.27 | 142.29 | 212.36 | 175.29 | 145.92 | 204.25 | 169.93 | 141.66 | 199.37 | 174.84 | 145.78 | 208.12 | 164.35 | 138.28 | 194.59 |
| 45 | 132.07 | 109.1 | 154.34 | 138.87 | 103.93 | 165.47 | 143.77 | 117.85 | 175.69 | 145.16 | 116.71 | 178.14 | 140.47 | 117.85 | 167.53 |
| 46 | 125.04 | 102.49 | 150.87 | 128.63 | 91.49 | 156.44 | 135.42 | 109.66 | 167.71 | 137.5 | 109.21 | 172.45 | 131.11 | 109.01 | 160.04 |
| 47 | 115.77 | 92.29 | 145.62 | 87.23 | 49.41 | 135.4 | 114.15 | 83.67 | 145.61 | 111.48 | 82.35 | 164.65 | 112.43 | 86.53 | 149.63 |
| 48 | 108.09 | 84.91 | 141.8 | 72.17 | 35.43 | 128.25 | 102.53 | 76.51 | 135.11 | 99.48 | 68.25 | 156.42 | 99.2 | 71.03 | 140.63 |
| 49 | 52.8 | 24.63 | 77.98 | 36.16 | 18.26 | 58.33 | 36.54 | 15.2 | 60.85 | 38.36 | 15.85 | 69.58 | 50.71 | 23.14 | 87.64 |
| 50 | 31.43 | 23.54 | 40.62 | 28.02 | 18.36 | 48.74 | 33.11 | 19.47 | 71.1 | 30.26 | 19.17 | 57.99 | 57.41 | 35.23 | 91.59 |
| 51 | 29.39 | 22.16 | 37.34 | 26.48 | 17.19 | 45.54 | 29.97 | 17.25 | 64.94 | 28.14 | 17.74 | 54.28 | 53.1 | 32 | 87.51 |
| 52 | 23.88 | 17.48 | 32.88 | 19.54 | 12.31 | 33.75 | 19.96 | 12.1 | 36.93 | 21.49 | 12.14 | 46.63 | 36.1 | 15.03 | 68.93 |
| 53 | 17.47 | 10.3 | 27.33 | 13.36 | 8.01 | 22.61 | 12.7 | 7.14 | 24.18 | 14.6 | 7.87 | 33.19 | 17.56 | 7.96 | 36.9 |
| 54 | 4.83 | 1.94 | 14.49 | 4.57 | 2.54 | 8.47 | 4.06 | 1.94 | 8.14 | 4.44 | 2.32 | 9.46 | 4.57 | 2.08 | 8.97 |
| 55 | 1.53 | 0.58 | 3.4 | 1.49 | 0.71 | 2.93 | 1.05 | 0.43 | 2.48 | 1.45 | 0.63 | 3.03 | 1.37 | 0.49 | 2.99 |

For reference, node numbers are in the following figure:

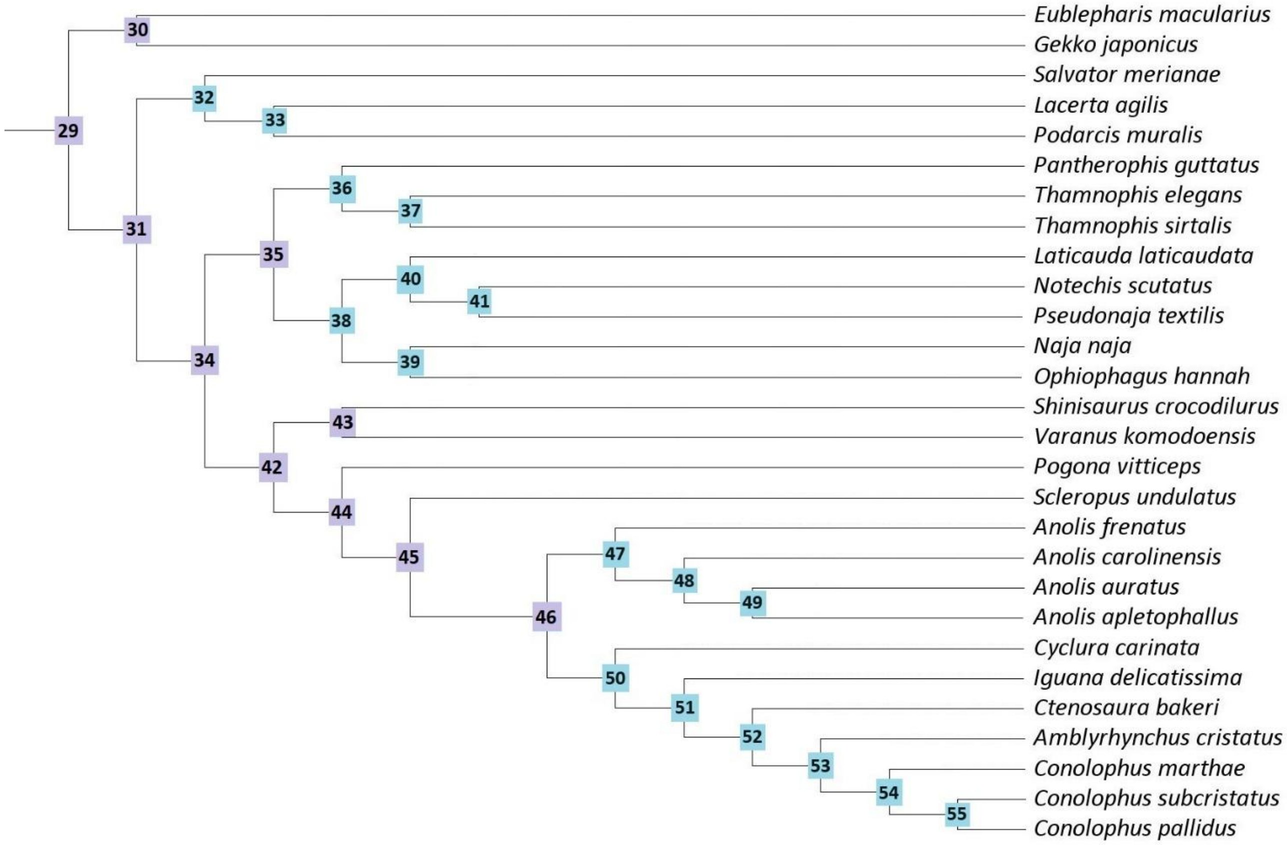

**Table S11: Genome-wide heterozygosity (θ), average length of ROH (with minimum and maximum values) and the proportion of the genome in ROH using a 500 kbp, 1 Mbp and 2 Mbp window sizes.**

| Species | Window | θ | ROH length (Mbp) | ROH (%) |
|---|---|---|---|---|
| ***Amblyrhynchus cristatus*** | 500 kbp | 0.00207 | 1.00 (1.00 – 1.00) | 0.13 |
| | 1 Mbp | 0.00207 | 1.00 (1.00 – 1.00) | 0.17 |
| | 2 Mbp | 0.00206 | 1.00 (1.00 – 1.00) | 0.13 |
| ***Conolophus marthae*** | 500 kbp | 0.00105 | 2.28 (2.21 – 2.36) | 14.71 |
| | 1 Mbp | 0.00105 | 2.27 (2.22 – 2.29) | 14.67 |
| | 2 Mbp | 0.00106 | 2.29 (2.22 – 2.35) | 14.76 |
| ***Conolophus pallidus*** | 500 kbp | 0.00020 | 1.46 (1.25 – 1.48) | 1.73 |
| | 1 Mbp | 0.00020 | 1.46 (1.25 – 1.48) | 1.73 |
| | 2 Mbp | 0.00020 | 1.46 (1.25 – 1.48) | 1.73 |
| ***Conolophus subcristatus*** | 500 kbp | 0.00185 | 3.25 (2.00 – 4.20) | 0.53 |
| | 1 Mbp | 0.00186 | 3.25 (2.00 – 3.67) | 0.53 |
| | 2 Mbp | 0.00185 | 3.25 (2.00 – 3.67) | 0.53 |
| ***Ctenosaura bakeri*** | 500 kbp | 0.00037 | 2.50 (1.65 – 5.00) | 1.04 |
| | 1 Mbp | 0.00037 | 2.64 (1.70 – 5.00) | 1.01 |
| | 2 Mbp | 0.00037 | 2.64 (1.71 – 5.00) | 1.01 |
| ***Cyclura carinata*** | 500 kbp | 0.00100 | 1.00 (1.00 – 1.00) | 1.24 |
| | 1 Mbp | 0.00101 | 1.00 (1.00 – 1.00) | 1.26 |
| | 2 Mbp | 0.00100 | 1.00 (1.00 – 1.00) | 1.73 |
| ***Iguana delicatissima*** | 500 kbp | 0.00021 | 1.85 (1.52 – 2.07) | 33.25 |
| | 1 Mbp | 0.00021 | 1.89 (1.61 – 2.07) | 33.51 |
| | 2 Mbp | 0.00021 | 1.91 (1.65 – 2.07) | 33.19 |

**Table S12: Mutational load summary of functional variants annotated with SnpEff.** The columns correspond to: proportion of derived alleles across all sites (relative load), the total number of loss-of-function (LoF) variants, LoF variants with two derived alleles (realized), LoF variants with one derived allele (masked), and percentage of realized LoF variants relative to total LoF variants.

| **Species** | **Relative load** | **Total LoF** | **Realized LoF** | **Masked LoF** | **Realized Ratio (%)** |
|---|---|---|---|---|---|
| *Amblyrhynchus cristatus* | 1.06661773 | 10,363 | 712 | 9,651 | 6.87 |
| *Conolophus marthae* | 1.07833844 | 3,341 | 272 | 3,069 | 8.14 |
| *Conolophus pallidus* | 1.13763066 | 1,654 | 227 | 1,427 | 13.72 |
| *Conolophus subcristatus* | 1.05691883 | 6,516 | 389 | 6,127 | 5.97 |

**Table S13: Squamate species used in the analyses. The species with an asterisk were not included in the positive selection analysis.**

| Species | Common name | Suborder | Database | Accession |
|---|---|---|---|---|
| *Amblyrhynchus cristatus* | Marine iguana | Iguania | López-Delgado et al. | Acri_T1 |
| *Anolis apletophallus* | Slender anole | Iguania | Harvard Dataverse | 10.7910/DVN/NGSGCG |
| *Anolis auratus* * | Grass anole | Iguania | Harvard Dataverse | 10.7910/DVN/NGSGCG |
| *Anolis carolinensis* * | Green anole lizard | Iguania | UniProt | UP000001646 |
| *Anolis frenatus* * | Bridled anole | Iguania | Harvard Dataverse | 10.7910/DVN/NGSGCG |
| *Conolophus marthae* | Pink iguana | Iguania | López-Delgado et al. | Cmar_T22 |
| *Conolophus pallidus* | Santa Fe land Iguana | Iguania | López-Delgado et al. | Cpal_T1 |
| *Conolophus subcristatus* | Galápagos land iguana | Iguania | López-Delgado et al. | Csub_T2 |
| *Ctenosaura bakeri* | Útila spiny-tailed iguana | Iguania | López-Delgado et al. | Cbak_T1 |
| *Cyclura carinata* | Turks and Caicos rock iguana | Iguania | López-Delgado et al. | Ccar_T1 |
| *Eublepharis macularius* | Leopard gecko | Gekkota | pers comm | E_macularius_v1.0 |
| *Gekko japonicus* | Japanese gecko | Gekkota | NCBI RefSeq | GCF_001447785.1 |
| *Iguana delicatissima* | Lesser Antillean iguana | Iguania | López-Delgado et al. | Idel_T1 |
| *Lacerta agilis* | Sand lizard | Laterata | VGP | GCA_009819535.1 |
| *Laticauda laticaudata* * | Blue-ringed sea krait | Serpentes | UniProt | UP000694406 |
| *Naja naja* | Indian cobra | Serpentes | UniProt | UP000694559 |
| *Notechis scutatus* | Tiger snake | Serpentes | UniProt | UP000504612 |
| *Ophiophagus hannah* * | King cobra | Serpentes | UniProt | UP000018936 |
| *Pantherophis guttatus* | Corn snake | Serpentes | UniProt | UP000515157 |
| *Podarcis muralis* | Common wall lizard | Laterata | UniProt | UP000472272 |
| *Pogona vitticeps* | Australian dragon lizard | Iguania | UniProt | UP000504619 |
| *Pseudonaja textilis* | Eastern brown snake | Serpentes | UniProt | UP000472273 |
| *Salvator merianae* * | Tegu lizard | Laterata | UniProt | UP000694421 |
| *Sceloporus undulatus* | Eastern fence lizard | Iguania | AUrora | GCA_019175285.1 |
| *Shinisaurus crocodilurus* | Chinese crocodile lizard | Anguimorpha | GigaDB | PRJNA353147 |
| *Thamnophis elegans* | Garter snake | Serpentes | VGP | GCA_009769535.1 |
| *Thamnophis sirtalis* | Common garter snake | Serpentes | UniProt | UP000504617 |
| *Varanus komodoensis* | Komodo dragon | Anguimorpha | UniProt | UP000694545 |

**Table S14: Number of positively selected genes in each foreground lineage**, including the genes that were uniquely under positive selection in their lineage and those that were not exclusive to a particular lineage.

| Foreground lineage | Lineage-exclusive | Non-exclusive |
|---|---|---|
| *Amblyrhynchus cristatus* | 421 | 2,413 |
| *Conolophus marthae* | 100 | 1,114 |
| *Conolophus pallidus* | 59 | 781 |
| *Conolophus subcristatus* | 59 | 784 |
| Galápagos lineage | 108 | 1,265 |
| Land lineage | 248 | 2,894 |
| *C. pallidus – C. subcristatus* lineage | 51 | 646 |

**Table S15: Summary of gene family expansions and contractions obtained with CAFE across the Iguanidae family.**

| Clade | Expansions | Contractions |
|---|---|---|
| *Amblyrhynchus cristatus* | 1,040 | 1,701 |
| *Anolis apletophallus* | 537 | 7,408 |
| *Anolis aureatus* | 1,033 | 1,230 |
| *Anolis carolinensis* | 618 | 3,136 |
| *Anolis frenatus* | 864 | 5,460 |
| *Conolophus marthae* | 870 | 1,223 |
| *Conolophus pallidus* | 894 | 1,013 |
| *Conolophus subcristatus* | 536 | 1,031 |
| *Ctenosaura bakeri* | 982 | 1,853 |
| *Cyclura carinata* | 2,313 | 2,791 |
| *Iguana delicatissima* | 952 | 3,415 |
| *Pogona vitticeps* | 627 | 3,263 |
| *Sceloporus undulatus* | 1,826 | 4,543 |

**Table S16: Substitution rates per 100 million years estimated with MCMCtree for single-gene orthologs and UCEs across different posterior distributions and subsets.**

| Distribution | Marker | Completeness | Subset | Filtering | mu |
|---|---|---|---|---|---|
| Skewed-normal | SGOs | All | All | Saturation + hidden paralogy + phylogenetic signal | 0.0980 |
| | UCEs | 75% matrix | All | No | 0.1012 |
| | | | 500 UCEs | Systematic bias | 0.0480 |
| | | | | genesortR default | 0.0933 |
| | | | 1,000 UCEs | Systematic bias | 0.0577 |
| | | | | genesortR default | 0.1012 |
| | | 95% matrix | All | No | 0.1010 |
| | | | 500 UCEs | Systematic bias | 0.0484 |
| | | | | genesortR default | 0.0969 |
| | | | 1,000 UCEs | Systematic bias | 0.0573 |
| | | | | genesortR default | 0.0989 |
| Uniform | SGOs | All | All | Saturation + hidden paralogy + phylogenetic signal | 0.0939 |
| | UCEs | 75% matrix | All | No | 0.0958 |
| | | | 500 UCEs | Systematic bias | 0.0450 |
| | | | | genesortR default | 0.1007 |
| | | | 1,000 UCEs | Systematic bias | 0.0561 |
| | | | | genesortR default | 0.0948 |
| | | 95% matrix | All | No | 0.1023 |
| | | | 500 UCEs | Systematic bias | 0.0468 |
| | | | | genesortR default | 0.0935 |
| | | | 1,000 UCEs | Systematic bias | 0.0565 |
| | | | | genesortR default | 0.0955 |
| Cauchy | SGOs | All | All | Saturation + hidden paralogy + phylogenetic signal | 0.0562 |
| | UCEs | 75% matrix | All | No | 0.0646 |
| | | | 500 UCEs | Systematic bias | 0.0279 |
| | | | | genesortR default | 0.0663 |
| | | | 1,000 UCEs | Systematic bias | 0.037 |
| | | | | genesortR default | 0.0571 |
| | | 95% matrix | All | No | 0.0613 |
| | | | 500 UCEs | Systematic bias | 0.029 |
| | | | | genesortR default | 0.0625 |
| | | | 1,000 UCEs | Systematic bias | 0.0362 |
| | | | | genesortR default | 0.0637 |

**Table S17: Fossil calibration points and distributions constructed from minimum and maximum constraints in the three strategies.** The uniform distribution in Strategy 1 includes the minimum age, maximum age, and upper and lower bound probabilities. The skewed-normal distribution in Strategy 2 sets the location, scale and shape parameters. The Cauchy distribution in Strategy 3 sets the mode as the minimum age and no maximum age, the tail parameter and left tail probability.

| **Split** | **Fossil** | **Source** | **Min age** | **Max age** | **Strategy 1** | **Strategy 2** | **Strategy 3** |
|---|---|---|---|---|---|---|---|
| Root | Lepidosaurian jaw, *Eichstaettisaurus schroederi* | 1,2 | 150 | 241.4 | B(1.5,2.414,0.0,0.025) | SN(1.55,0.41,9) | B(1.5,2.414,0.0,0.025) |
| *Gekko japonicus – Eublepharis macularius* | *Yantarogekko balticus* | 3 | 44 | 63.3 | B(0.44,0.633,0.0,0.025) | SN(0.44,0.09,10) | L(0.44,0,10,0.001) |
| *Salvator merianae - Pantherophis guttatus* | *Ptilodon* | 4 | 111 | 187.3 | B(1.11,1.873,0.0,0.025) | SN(1.15,0.34,11) | L(1.11,0,10,0.001) |
| *Pantherophis guttatus - Shinisaurus crocodilurus* | *Eophis underwoodi* | 5 | 166.1 | 168.3 | B(1.661,1.683,0.0,0.025) | SN(1.665,0.01,10) | L(1.661,0,10,0.001) |
| *Shinisaurus crocodilurus - Varanus komodoensis* | *Palaeosaniwa* | 6 | 71 | 113 | B(0.71,1.13,0.0,0.025) | SN(0.75,0.19,8) | L(0.71,0,10,0.001) |
| *Pantherophis guttatus – Naja naja* | *Rukwanyoka holmani* | 7 | 23 | 28.4 | B(0.23,0.284,0.0,0.025) | SN(0.24,0.01,5) | L(0.23,0,10,0.001) |
| *Shinisaurus crocodilurus - Pogona vitticeps* | *Dorsetisaurus* | 8 | 148 | 156.6 | B(1.48,1.566,0.0,0.025) | SN(1.48,0.04,5) | L(1.48,0,10,0.001) |
| *Pogona vitticeps – Sceloporus undulatus* | *Saichangurvel davidsoni* | 9 | 71 | 113 | B(0.71,1.13,0.0,0.025) | SN(0.75,0.19,8) | L(0.71,0,10,0.001) |
| *Sceloporus undulatus – Anolis frenatus* | *Phrynosomatidae* | 10 | 72.1 | 83.6 | B(0.721,0.836,0.0,0.025) | SN(0.73,0.05,6) | L(0.721,0,10,0.001) |
| *Anolis frenatus – Cyclura carinata* | *Afairiguana avius* | 11 | 48 | 58 | B(0.48,0.58,0.0,0.025) | SN(0.48,0.04,3) | L(0.48,0,10,0.001) |

**Supplementary Information References**
